# Characterisation of human hair follicle development

**DOI:** 10.1101/2024.07.26.605346

**Authors:** Zoe R. Sudderick, James D. Glover, Cameron Batho-Samblas, Barbara Bo-Ju Shih, Denis J. Headon

**Affiliations:** The Roslin Institute and Royal (Dick) School of Veterinary Studies, University of Edinburgh, Edinburgh, UK; Division of Biomedical and Life Sciences, Faculty of Health and Medicine, Lancaster University, Lancaster, UK

## Abstract

Humans have a characteristic distribution of hair across the body. Visible, relatively long and thick terminal hair fibres are present on the scalp and eyebrows in childhood, and are stimulated to grow on other parts of the body, such as the beard and armpits, by hormones during puberty. The short and fine vellus hairs, in contrast, are not readily visible and cover most of the body, including the face. Here we report quantification of the timing and characteristics of hair follicle development in human embryogenesis, from gestational weeks 8 to 19, and compare this to mouse hair follicle development. We find that human hair follicles develop first on the head, where we identify several distinct initiation sites, followed by the torso. Although terminal and vellus hair follicles have clear differences in the adult, both hair types initially develop from placodes and dermal condensates of similar size. Once their development is initiated, we find that human hair follicles grow and mature at the same rate, regardless of anatomical location, but have different density at different body sites. These findings suggest that regional hair differences in human skin, such as the distinction between scalp and forehead, are largely caused by processes acting after the initial hair follicle morphogenesis. Efforts to understand the evolution of human ‘hairlessness’ should, therefore, focus on genetic and cellular events that take place after hair follicle morphogenesis. Finally, we compared human skin appendages, including eccrine sweat glands, with those in mouse. We found that molecular markers, such as EDA, EDAR, SOX2 and WNT pathway components, are broadly similar in expression between both species, although specific differences do exist. Together with comparison of morphology and gene expression, these results support the use of embryonic mouse primary hair follicles as a model for human hair follicle development.

## Introduction

Hair follicle development initiates prior to birth in both mouse and human. Hair follicles begin as an epidermal thickening, termed a placode (PC), which acquires a dermal condensate (DC) composed of mesenchymal fibroblasts. This composite structure grows down into the dermis, driven by rapid epithelial proliferation and develops though defined stages to become a mature, fibre-producing, follicle (Schneider et al., 2009). This initial morphogenesis phase is followed by lifelong cycling through three different stages of variable duration; regression (catagen), resting (telogen), and regeneration (anagen), each cycle producing a new hair fibre (Botchkarev and Paus, 2003).

Human and mouse skin both carry multiple different types of hair. In human development, fine unpigmented lanugo hair fibres are produced first and cover the fetus, usually being shed prior to or around birth (Domagala et al., 2017, Verhave et al., 2022). The adult human body carries two hair types; long, thick terminal hair fibres and tiny unpigmented vellus hairs, each present in clearly demarcated body regions, often with a sharp boundary or ‘hairline’ between them. The scalp and eyebrows produce terminal hair in children, with beard and axillary terminal hair produced under hormonal influences beginning at puberty (Randall, 2007). In contrast, in the mouse coat the different hair types of straight guard (also called monotrich or tylotrich), awl, kinked auchene, and zigzag are interspersed across the body (Dry, 1926). Mouse hair types develop in sequence through the last third of gestation (Mann, 1962), with primary guard hairs developing first followed by secondary awl and auchene, and finally tertiary zigzag hairs (Duverger and Morasso, 2009). There is no equivalent of a lanugo hair cycle in mouse.

All hair follicle types in mouse rely on WNT/β-catenin signalling in both epithelium and mesenchyme to form (Saxena et al., 2019). However, the primary hair follicles selectively require EDA/EDAR signalling (Headon and Overbeek, 1999), while secondary hair follicles rely instead on LEF1 (van Genderen et al., 1994) and Noggin (Botchkarev et al., 2002). FGF20 is required by both the primary and secondary waves of hair follicles and has a particular role in the recruitment of the dermal condensate (Biggs et al., 2018) The transcription factor SOX2 is a marker of the dermal condensate and dermal papilla in primary and secondary hair follicles, but is not expressed in the last-forming and smallest tertiary hair follicles (Driskell et al., 2009), thus serving as a molecular marker to distinguish between these hair types in mouse skin.

Here we describe hair follicle formation during human development at molecular and morphological levels, and compare these processes to those operating during development of the primary hair follicles in the mouse.

## Results

### Human hair follicle initiation sites and growth

Through analysis of human fetal skin sections from weeks 10 to 19 estimated gestational age (EGA), we identified stages of hair follicle morphogenesis in human that match those reported in mice (Figure 1a) (Saxena et al., 2019, Paus et al., 1999). To define the locations on the body at which hair follicle development initiates we used whole mount RNA *in situ* hybridisation to visualise expression of the hair placode marker *EDAR*, together with histological assessment of the skin. We first observed punctate, regular and bilaterally symmetrical spots of expression of *EDAR* at 10 weeks EGA on the medial part of each eyebrow, close to the nose, and the lateral edges of the upper lip. At 11 weeks, the organised array of spots on the eyebrows had extended both laterally and medially, filling the space between the eyebrows and the formation of placodes on the upper lip had also extended medially, towards the philtrum. New areas of punctate *EDAR* expression were present bilaterally below the eyes on the upper cheeks, and at the lateral edges of the lower lip. By 12 weeks hair placodes had extended to cover the cheeks and filled the upper lip. New areas of placode initiation were also observed on the scalp, temples, forehead, and chin, thereby covering most of the face (Figure 1b). The early initiation sites of supraorbital eyebrow, the chin, and upper and lower lips correspond to clearly demarcated sites of eyebrow and beard in adult. The observed discrete placode initiation sites below each eye, on the other hand, are not associated with any obvious feature present in adult skin. Areas of punctate *EDAR* expression were always symmetric on the right and left of the head (Figure 1c).

**Figure 1:**
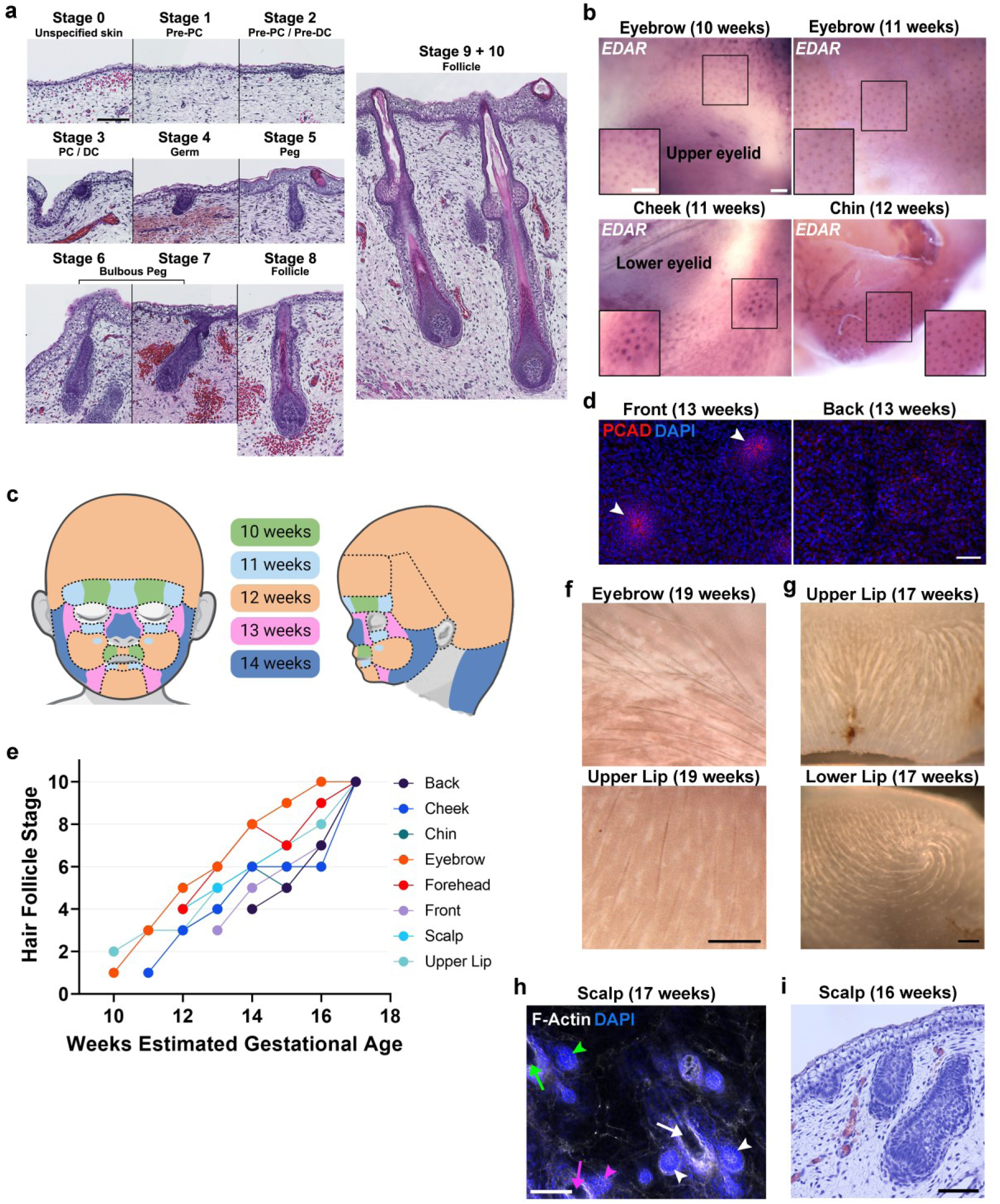
Hair follicle initiation and development. **(a)** Developmental stages of human hair follicles from H&E stained fetal skin sections. **(b)** Detection of *EDAR* transcripts by wholemount RNA in situ hybridisation of human fetal skin showing sites of hair follicle placode initiation. **(c)** Schematic of hair follicle placode initiation sites and corresponding estimated gestational age (EGA) on the human head. **(d)** Wholemount immunofluorescence detecting P-CADHERIN (PCAD) on week 13 back and front skin. Hair placodes are indicated by white arrowheads. **(e)** Quantification of the most mature hair follicle morphogenesis stage observed by EGA for eight areas of skin. **(f)** Brightfield images of hair fibre pigmentation. **(g)** Brightfield images showing local hair fibre orientation. **(h)** Wholemount human scalp skin stained with phalloidin (to detect F-actin) showing formation of follicular units with groups of hair follicles of different developmental stages present. More mature central hair follicles are indicated with arrows, younger accessory follicles with arrowheads. Each follicular unit is indicated by different coloured arrows/arrowheads (green, purple, and white) **(i)** H&E stained section showing follicles of different developmental stage in close proximity. PC = Placode, DC = Dermal Condensate. Scale bars: **a, h, i =** 100 μm; **b =** 250 μm; **c =** 50 μm; **f, g =** 500 μm

The first hair follicles produced in mouse are the sensory vibrissae on the face. Primary hair placode formation then initiates at the flanks, adjacent to the mammary gland rudiments under the forelimbs, and spreads as a rapid wave dorsally and posteriorly (Painter et al., 2021). In contrast, in human development we find that on the torso the placodes form on the front (ventrum) before the back (dorsum) (Figure 1d).

By quantifying the most advanced hair follicle morphogenesis stage observed for each area of skin we find that all regions undergo downgrowth and maturation at the same rate, with follicles taking approximately six weeks to reach stage 10 and display hair fibres emerging from the skin surface. As hair follicles on all skin regions grow and mature at the same rate, differences in hair follicle stage arise only from earlier or later initiation of hair follicle formation in a given region (Figure 1e).

Pigmentation of hair fibres was first observed at 19 weeks in the eyebrow, upper lip and forehead, which is expected as these follicles are the earliest to appear and thus the most mature at this age (Figure 1f). Orientation of lanugo hair fibres varies between areas of the face but appears to be related to the specific topography of a region, such as the whorls often present at the corners of the lower lip and the changes in orientation across the medial cleft of the upper lip (Figure 1g). Hair follicles in the adult scalp are arranged in clusters termed follicular units that contain two to four follicles (Headington, 1984). This arrangement of a mature central follicle and flanking less mature follicles can be observed from week 17 on the scalp (Figure 1h). We observed continuous formation of placodes in all areas of human fetal skin analysed, with some new placodes observed at all ages, resulting in different ages of follicles being observed in close proximity (Figure 1i).

### Skin regional differences in human hair follicle characteristics

In mice, different hair follicle types can be distinguished by several features. Primary hair follicle placodes have a larger diameter than those of the later-forming secondary and tertiary follicles, as well as a greater hair fibre width, lower placode density, and a longer mature follicle (Duverger and Morasso, 2009). Whisker follicle placodes are larger at inception than the primary hair placodes, and the whiskers are longer and thicker than the guard hairs that the primary hair follicles produce (Mou et al., 2008). Thus, in mouse, the size of the placode upon its formation is closely related to the size of the hair follicle and hair fibre that it produces. This could occur through an inductive templating, whereby placodes recruit an underlying dermal condensate in proportion to the former’s area via epithelial signals such as FGF20 and SHH (Huh et al., 2013, Glover et al., 2017, Qu et al., 2022).

To determine whether human hair follicle primordia differ in size in different regions of the skin, potentially corresponding to the distinct sizes of the hair fibres at these locations in postnatal skin, we assessed early hair placode diameter from H&E sections and compared these to mouse back skin primary follicles. We found that these range from 26 to 94 μm, with a mean placode diameter for each region of approximately 50 μm. No statistically significant difference in placode size was detected between human skin regions, or between the sizes of human and mouse primary follicles (Ordinary one-way ANOVA, P = 0.5160, F (8,156) = 0.9025) (Figure 2a). Thus, the size of the hair placode does not predict skin region differences in hair characteristics in human development.

**Figure 2:**
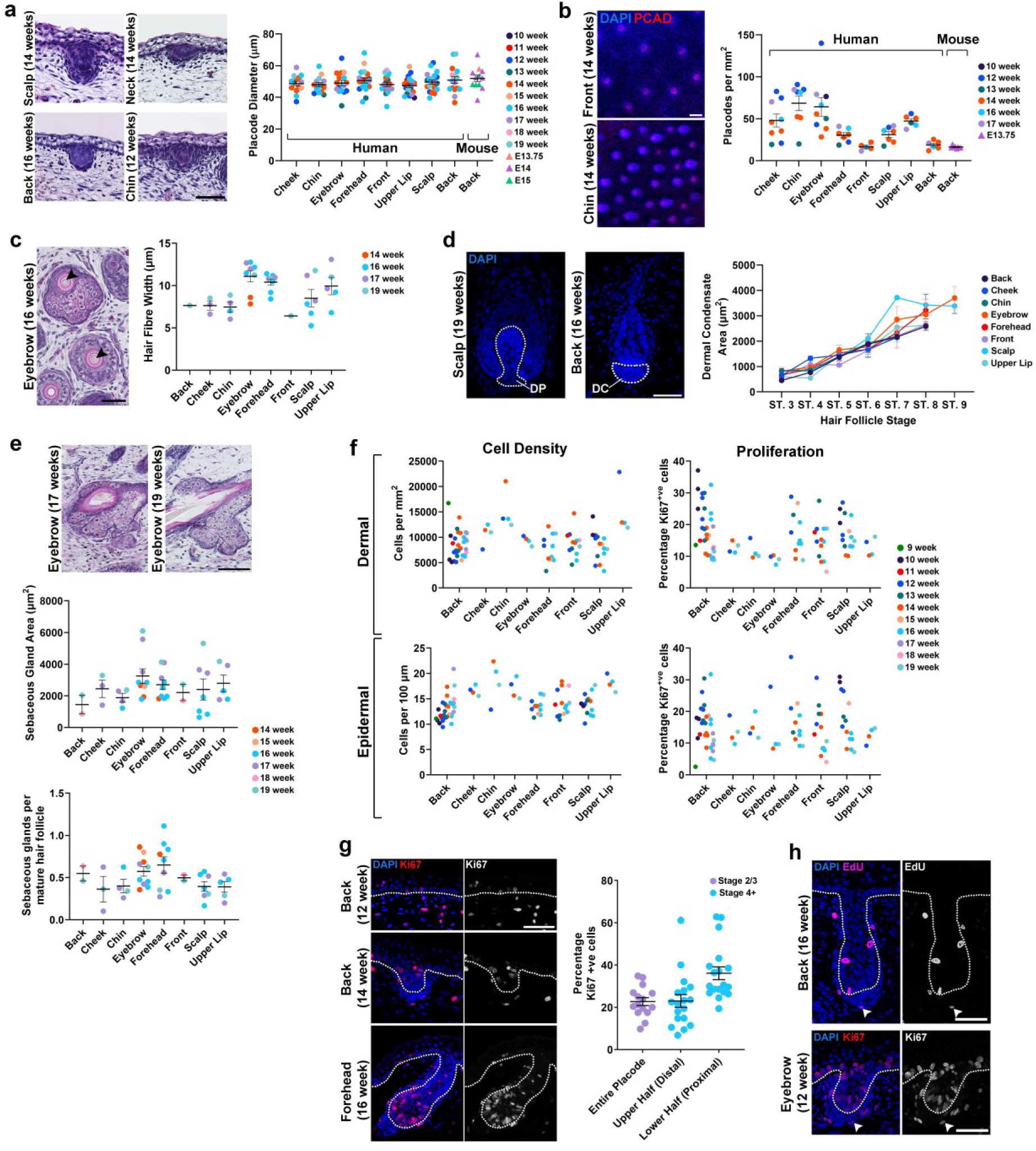
Morphometrics of the developing human hair follicle and skin. **(a)** H&E stained early hair placodes from different developmental stages and skin regions (left). Placode diameter quantification in mouse and human skin (right). Embryonic stages as indicated. **(b)** Wholemount immunofluorescent detection of P-CADHERIN (PCAD) on 14 week EGA front and chin skin (left). Quantification of placode density from eight human skin areas and mouse back skin (right). **(c)** H&E stained section of 16 week eyebrow skin showing hair follicle cross sections. Arrowheads indicate hair fibres (left). Hair fibre width on different skin areas coloured by age (right). **(d)** DAPI staining showing a dermal papilla (DP) and a dermal condensate (DC); dotted lines outline the area measured (left). Quantification of dermal condensate area on tissue sections by hair follicle stage for eight areas of skin; each dot represents the mean for each skin area (right). **(e)** H&E stained eyebrow sections showing sebaceous glands attached to a hair follicle (above). Mean sebaceous gland area from tissue sections of eight areas of skin at different ages (middle). Number of sebaceous glands detectable per mature hair follicle on different areas of human skin (below). **(f)** Quantification of cell density and proliferation (defined by Ki67 immunofluorescence) in the epidermis and dermis at different ages and across anatomical sites. **(g)** Immunofluorescent detection of Ki67 in unpatterned skin, an early hair follicle, and a later stage follicle (left). Quantification of proliferation in early hair follicles and in the upper (distal) or lower (proximal) half of the epithelial component of follicles at stages 5-7. Each dot represents an individual hair follicle (right). **(h)** Fluorescent detection of incorporated EdU and of Ki67 protein showing proliferating cells in hair follicle dermal condensates (arrowheads). A dotted line indicates the epidermal-dermal boundary. For graphs each dot represents an individual specimen unless otherwise stated. Scale bars: **a, c, d, e, g, h =** 50 μm; **b =** 100 μm

We find that the density of human hair placodes varies across the body (Ordinary one-way ANOVA, P= <0.0001, F (8,56) = 8.872), with the eyebrow and chin showing the highest densities and the trunk the lowest. On some facial regions there appears to be a link between developmental stage and placode density, with younger developmental stages often having higher follicle density than older ones. This suggests that the pattern of follicles may be fixed early on the face, with subsequent growth and expansion of skin area decreasing the density of the hair follicle primordia already present. The torso, scalp and forehead do not appear to show this phenomenon and instead have similar placode densities at all stages, suggesting continued insertion of new hair placodes as the skin expands at these sites (Figure 2b). We found that mouse back skin primary hair follicles are laid out at approximately the same density as follicle primordia in human fetal back skin (Tukey’s multiple comparisons test, P = >0.9999).

Unlike placode size, hair fibre diameter does display some significant differences between skin regions (Ordinary one-way ANOVA, P= 0.0127, F (7,28) = 3.203). Hair fibres on the eyebrow and upper lip are the largest, potentially as these are the first areas to form follicles so are the most developed at the stages of analysis. The chin, however, is the next area to form follicles and shows a smaller fibre width than the scalp and forehead, which develop follicles slightly later. Follicles on the torso show the lowest hair fibre width, with back similar to that of cheek and chin (Figure 2c).

Smaller fibre widths in the back, front, and cheek compared to the coarser fibres of the eyebrow and scalp are consistent with these areas forming only vellus hairs in adult life. The larger fibre widths observed in the forehead are, however, not consistent with this relationship as the forehead forms only vellus hairs in adult skin.

Given the importance of the dermal papilla in defining hair follicle characteristics (Van Scott and Ekel, 1958, Jahoda, 1992), differences in the size of its embryonic precursor, the dermal condensate, could underlie later-emerging differences in size between human hair follicle types. However, we find that dermal condensate size on tissue sections increases steadily as the follicles grow through stages of morphogenesis, with significant differences only observed between the regions with the smallest (front skin) and two largest (eyebrow and scalp) dermal condensate areas (Tukey’s multiple comparisons test, Front vs Eyebrow p = 0.0471, Front vs Scalp p = 0.0408) (Figure 2d). These modest differences in dermal condensate and placode size suggest that neither predict whether a follicle will later produce a terminal or a vellus hair, but that the major distinctions in follicle size between vellus and terminal hairs emerge later in development.

The epithelial placode undergoes downgrowth into the dermis to produce the entire pilosebaceous unit of hair follicle and associated sebaceous gland. Postnatally, sebaceous glands are larger and more densely packed on the face and scalp than the torso or other body sites, making skin at these sites oilier (Tsatsou and Zouboulis, 2014). At the developmental stages examined we did not detect a statistically significant difference in the size or density of sebaceous glands between different areas of skin (Ordinary one-way ANOVA, size P= 0.5260, F (7,34) = 0.8887, and density P = 0.1393, F (7,34) = 1.712), when measured as mean sebaceous gland area, or as the number of sebaceous glands in relation the number of mature hair follicles observed per unit length of skin (Figure 2e).

The placode and dermal condensate are formed from cells recruited from the basal epidermis and upper dermis, respectively. We quantified basal epidermal and upper dermal cell density and proliferation in different body sites and ages (Figure 2f) using the proliferation marker Ki67 to determine whether there are distinct patterns of cell division associated with hair follicle initiation and development. When comparing areas of skin, we detected significant differences in dermal and epidermal cell density and in the proliferation of the dermis (Ordinary one-way ANOVA: epidermal cell density P = <0.0001, F (7,68) = 5.777, dermal cell density P = <0.0001, F (7,68) = 6.275, dermal proliferation P = 0.0004, F (7,67) = 2.170) while no significant difference was found between regions in the proliferation of the epidermis (Ordinary one-way ANOVA, P = 0.6982, F (7,67) = 0.2011). Although significant differences in proliferation and cell density of the epidermis and in proliferation of the dermis were detected when comparing across different ages, no obvious patterns related to hair follicle formation could be discerned (Ordinary one-way ANOVA: epidermal cell density P = 0.0204, F (10,68) = 2.322, epidermal proliferation P = <0.0001, F (10,67) = 5.623, dermal cell density P = 0.0525, F (10,68) = 1.954, dermal proliferation P = <0.0001, F (10,67) = 4.749).

In general, basal epidermal cell density increases while proliferation decreases as the skin matures, but we did not observe a discrete change coincident with the onset of hair follicle formation in different skin regions. Upper dermal cell density remains quite constant across skin development while proliferation decreases slightly, but, as observed for the epidermis, there is no discrete change in gross cell density at the time of follicle formation. This suggests that, unlike the events during feather formation (Ho et al., 2019), there is no clear demarcation of dense and loose dermis into tracts that might define the characteristics of different human hair follicle types. However, the eyebrow and beard regions do have a high density of dermal cells at the time of hair follicle formation, compared to other skin regions (Figure 2f), and these areas may be comparable to the feather tracts in avian skin. More broadly, the epidermal and dermal cell density of skin of facial regions (cheek, chin, eyebrow, and upper lip) appears to be higher than that of the trunk.

Within developing hair follicles, the number of proliferating cells is greater in the lower half of the epithelial component of the follicle (stage 4+) than the upper half (Tukey’s multiple comparisons test: P = 0.0038). The percentage in the lower half of these (stage 4+) follicles is also significantly greater than that of stage 3 and below follicles (Tukey’s multiple comparisons test: P = 0.0050) (Figure 2g). Proliferation in developing human hair follicles thus occurs mainly in the epithelial cells at the leading edge of the growing follicle closest to the dermal condensate, as in mouse (Saxena et al., 2019). In the dermal condensate itself, approximately 40% of follicles examined in sections showed at least one Ki67 positive cell, most of which were present at the periphery of the condensate (Figure 2h).

### Epidermal differentiation in relation to hair follicle formation

At the time of hair placode formation, the epidermis is multi-layered, with periderm as the most superficial layer, at least one suprabasal keratinocyte layer beneath it, and the more densely packed basal keratinocytes as the lowest layer. The periderm is a continuous layer of nucleated cells at week 14, these cells being larger than the suprabasal or basal keratinocytes. Periderm cells have complex Actin cytoskeletal structures at this stage, focalised in particular on the basal side of the cells, but unlike the basal and suprabasal layers they do not express α-catenin (Figure 3a).

**Figure 3:**
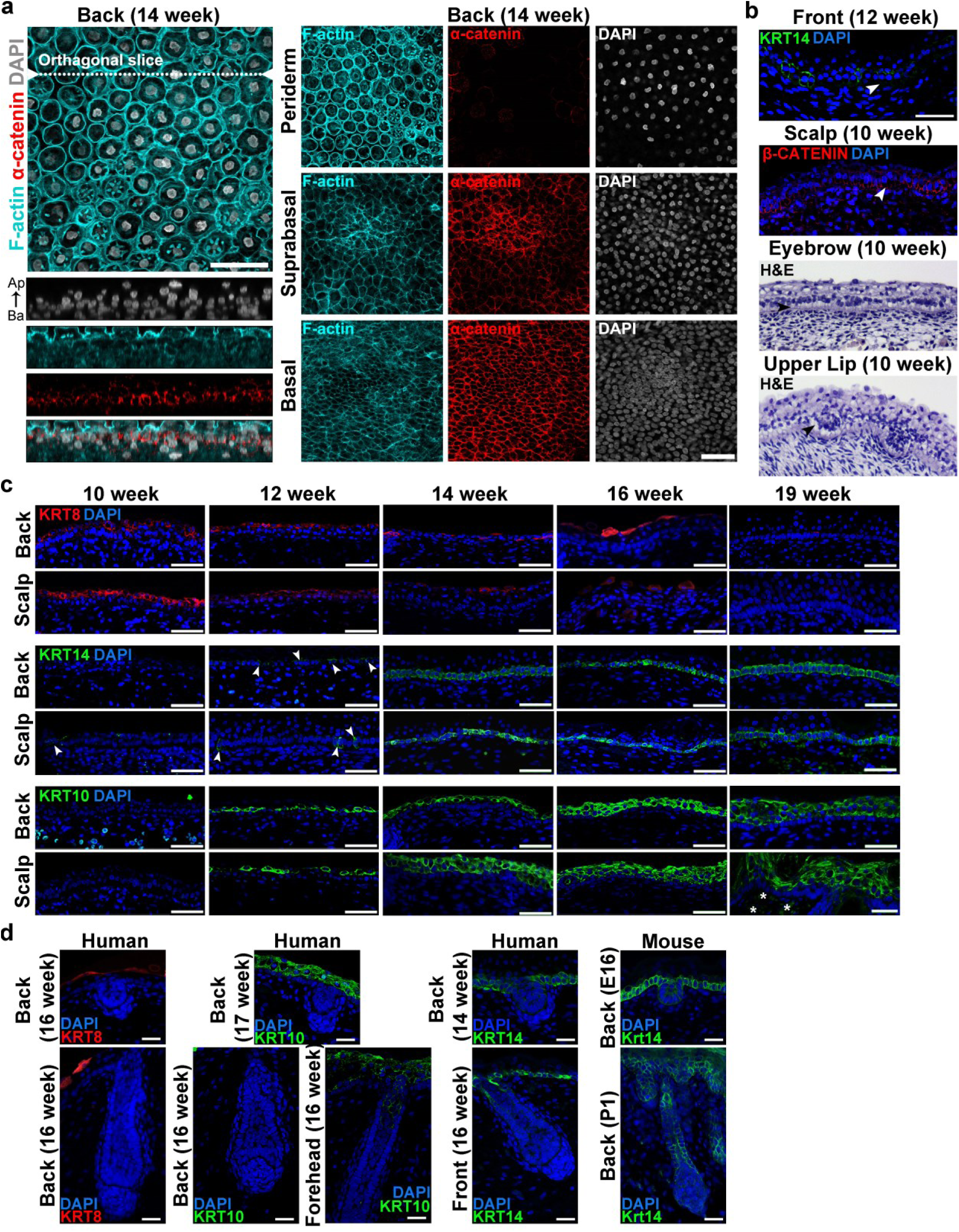
Timing of epidermal differentiation relative to hair follicle development. **(a)** Wholemount immunofluorescent detection of α-catenin and F-actin (stained with phalloidin) in different layers of 14 week back skin. Orthogonal projection, 3 μm z-plane intervals. **(b)** Immunofluorescent detection of KRT14 and β-catenin, and H&E staining showing apical localisation of nuclei in the basal epithelium with arrowheads indicating the cytoplasm in the basal side of the epithelial cells. **(c)** Immunofluorescent detection of Keratins (KRTs) 8, 14 and 10 in human back and scalp skin at 10 to 19 weeks EGA. Arrowheads indicate isolated positive cells; asterisks indicate autofluorescent blood cells. Immunofluorescent detection of KRT14 and β-catenin, and H&E staining showing apical localisation of nuclei in the basal epithelium with arrowheads indicating the cytoplasm in the basal side of the epithelial cells. **(d)** Immunofluorescent detection of Keratins 8, 10, and 14 in early and late human hair follicles, and KRT14 in mouse early and late primary hair follicles. Ap, apical; Ba, basal. Scale bars: **a, c, d** = 50 μm; **b** = 25 μm

Some sections of human skin displayed a clear gap between epidermis and dermis, visible on H&E stained and immunofluorescent tissue sections. This apparent gap arose from basal epidermal cells becoming columnar, with apically located nuclei and a large basal cytoplasm (Figure 3b). We observed this arrangement of basal epidermal cells at many skin sites at weeks 10 and 12 EGA, including around early hair placodes, but very rarely at later ages, and never on volar skin. We did not observe this configuration of epidermal cells in mouse skin.

We assessed the first appearance of specific epidermal keratins across the face, head and torso to assess relative rates of epidermal maturation and their timing relative to hair placode development. Keratins 8, 14 and 10 (KRT8, KRT14, KRT10) were assessed as markers of simple epidermis/periderm, basal epidermis and intermediate/suprabasal layers, respectively. KRT8 was detected as a continuous suprabasal layer in the back and scalp from 10 weeks EGA, becoming restricted to the periderm, and then lost as the periderm regresses. KRT14 is first detected in occasional isolated individual cells beneath the KRT8 expressing layer, at 10 weeks in the scalp (Figure 3c) and volar skin on the distal digit (Glover et al., 2023). Similar sporadic, isolated KRT14+ve cells are present at 11 weeks in the front skin (Supplementary Figure 1) and 12 weeks in the back skin. The number of KRT14 expressing cells steadily increases in the basal layer until week 14, when all regions examined had a continuous KRT14-expressing basal layer. KRT10 expression is also detected in suprabasal epidermis of the scalp and back skin from 12 weeks, with more layers of KRT10 positive cells observed as the epidermis thickens (Figure 3c). Thus, there are relatively small differences in epidermal maturation between different regions of hair-bearing skin, but K14 expression especially follows the same order as the development of hair follicle placodes, first the scalp, followed by the front, then the back skin.

In the vicinity of the hair follicles, KRT8 is not found within the follicle epidermis while KRT10 is found in the suprabasal layers above the developing placode and later marks an internal core of follicle epidermis in the infundibulum near the exit point of the shaft (Figure 3d). In human skin there is a significant decrease in the intensity of KRT14 expression in placodes and hair follicles compared to the interfollicular epidermis. This contrasts with mouse skin development, where KRT14 expression remains high in the developing follicles (Figure 3d).

### Cell signalling across the period of follicle development

In areas of skin where hair follicles develop, immunoreactivity for LEF1, a component and a readout of the canonical WNT pathway (Gupta et al., 2019), was stronger in epidermis than in the dermis. In general, as skin development proceeded across stages the dermal LEF1 signal reduced while the epidermal signal intensified. Dermal LEF1 was greater in facial skin, particularly in the upper dermis, than in skin of the scalp and the torso (Figure 4a). Of all body sites examined, the torso skin consistently had the least dermal LEF1 expression. The future beard and eyebrow regions had a high density of LEF1 positive, presumably WNT-activated, dermal cells, and these areas may be comparable to the high cell density dermal tracts of avian skin, which are also identifiable by increased LEF1 dermal expression (Supplementary Figure 2). LEF1 expression is strong in the early forming placode before it becomes restricted to the leading epithelial edge of downgrowing follicles. The dermal condensate exhibits LEF1 expression at levels greater than that of the surrounding dermis.

**Figure 4:**
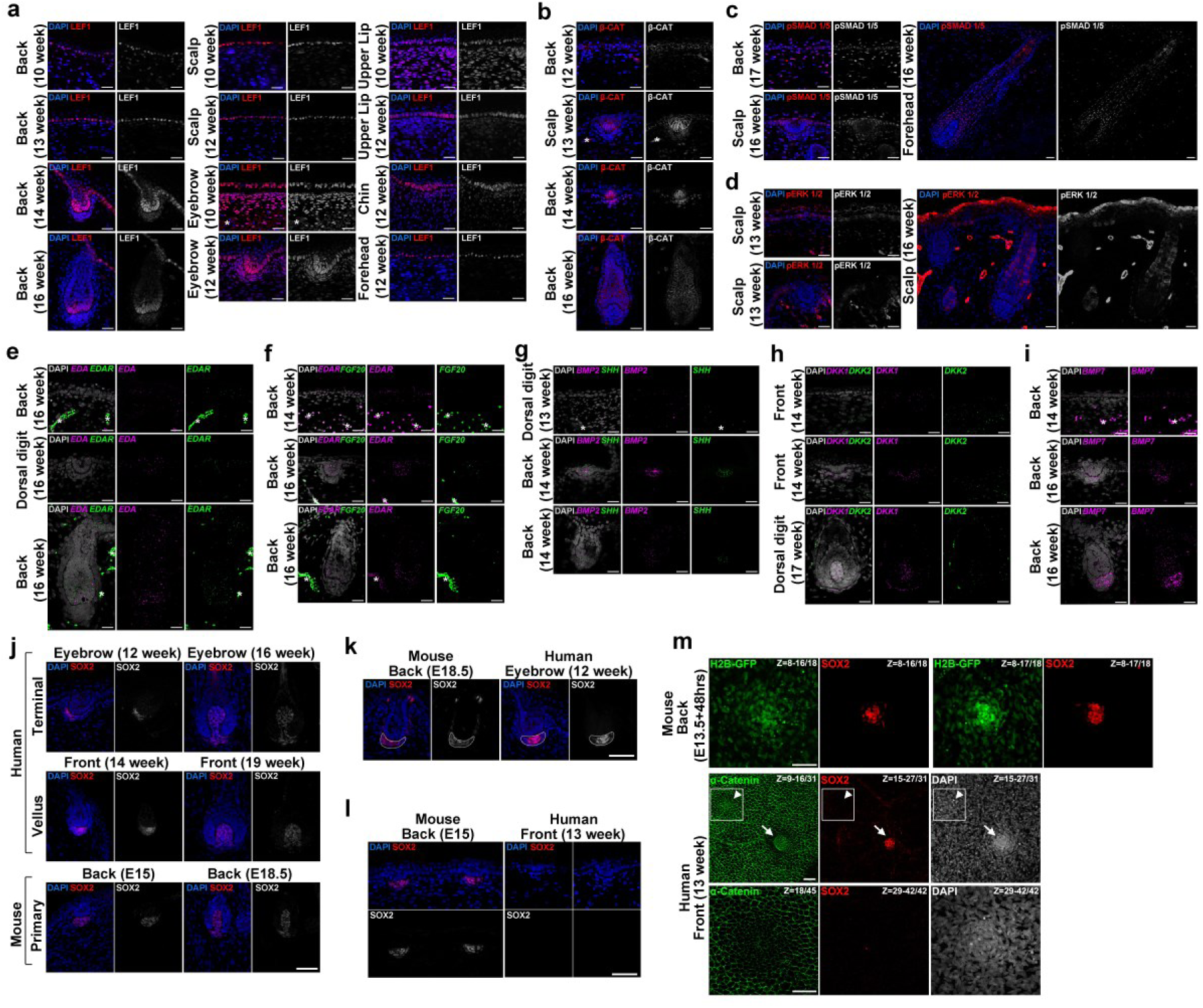
Expression of intercellular signalling factors, signal transducers, and transcriptional regulators during human hair follicle development. (a-d) Immunofluorescent detection of **(a)** LEF1, **(b)** β-catenin, **(c)** phospho-SMAD1/5, and **(d)** phospho-ERK1/2 in unpatterned skin, early hair placodes, and later stage follicles. LEF1 is detected in developing hair follicles and there is strong dermal LEF1 immunoreactivity in eyebrow, upper lip and chin, but not in scalp, forehead or back skin prior to and during early follicle formation. **(e-i)** RNA in situ hybridisation detecting expression of (e) *EDA* and *EDAR,* (f) *EDAR* and *FGF20,* (g) *BMP2* and *SHH*, (h) *DKK1* and *DKK2*, and (i) *BMP7*, in unpatterned skin, early hair follicles, and later stage follicles. **(j)** Immunofluorescent detection of SOX2 positive dermal condensates and dermal papillae in human terminal and vellus hair areas, and mouse primary hair follicles. **(k)** Immunofluorescent detection of mouse and human later stage follicles showing SOX2 localisation in dermal condensates. **(l)** Immunofluorescent detection of SOX2 in human and mouse early hair follicle primordia, showing earlier SOX2 expression in cells that are forming mouse dermal condensates as compared to human, where expression is detected only after condensate compaction. **(m)** Wholemount immunofluorescent detection of SOX2 in TCF/Lef::H2B-GFP mouse back skin and fetal human back skin showing earlier SOX2 expression in mouse dermal condensates than human. **a, b, e, f, g + i** = asterisks indicate appearance of autofluorescent blood cells. Scale bars: **a-i** = 25 μm; **j-m** = 50 μm

β-catenin expression is patchy in the epidermis of both trunk and scalp at 10 weeks EGA. This becomes continuous across the entire epidermis by 13 weeks, with strong signal in the basal layer and weaker signal in the suprabasal layers. Expression is consistently strong in the hair follicle epidermis, with evidence of nuclear localisation in the dermal condensate of early follicles on the torso (Figure 4b).

BMP signalling inhibits hair follicle formation in mouse (Mou et al., 2006, Cheng et al., 2014, Botchkarev et al., 1999). We visualised BMP signalling locations by detecting the active, phosphorylated, form of the BMP signal transducer SMAD1/5. This was present in both the epidermis and dermis at all areas and locations assessed. Phospo-SMAD1/5 (pSMAD1/5) was decreased in the developing placode, with low epidermal detection persisting as the follicle develops, while it was lost from the dermal condensate as it becomes a dermal papilla (Figure 4c).

Phospho-ERK1/2, which is stimulated in response to several growth factors including EGF and FGF, was detected in the developing epidermis and in mesenchymal cells surrounding early follicles on the scalp, but in more mature follicles phospho-ERK1/2 was detected in the upper/distal basal epithelium likely to be the developing outer root sheath. No signal was detected in the dermal condensate at any follicle stage examined, though dermal phospho-ERK1/2 was detected surrounding the condensates of late stage follicles (Figure 4d).

Prior to placode formation, *EDAR* transcript is detected at low levels throughout the epidermis, before becoming highly enriched in the developing placode. As hair follicle development continues, *EDAR* expression becomes restricted to the leading edge of the downgrowing follicle (Figure 4e and f). *EDA* transcript was detected within the placodes, with limited expression in the remainder of the epidermis and in the dermis both before and during placode formation (Figure 4e). Expression increases in the later stage follicles with the most intense signal present at the leading edge of the epidermal component. No expression was observed in the initial dermal condensate, but *EDA* transcript was detected in the condensate of more mature stage follicles as well as in the developing dermal sheath. For *EDAR*, these results match those reported during mouse primary hair follicle development (Headon and Overbeek, 1999), but *EDA* expression differs in that it is reduced in expression in mouse primary hair placodes (Laurikkala et al., 2002) and in feather primordia (Ho et al., 2019), but we find it to be intensified in expression in human hair follicle placodes.

*FGF20* expression was rarely detected in the skin before the formation of follicles and is expressed selectively in the hair placode as these emerge. *FGF20* expression decreased as follicle downgrowth progressed (Figure 4f). *BMP2* and *SHH* expression was detected in a similar pattern in the initiating placodes, and these genes remain expressed in the more proximal epithelial component of the downgrowing hair follicles (Figure 4g). At the time of hair follicle formation *DKK2* is detected broadly at low levels throughout the dermis, but not in the epidermis. *DKK1* is observed more sparsely in the dermis, increasing in the dermal condensates and then the dermal papillae of the developing hair follicles (Figure 4h). *BMP7* was detected in the epidermis before follicle formation, with expression intensifying in both epidermal placode and dermal condensate. In later follicles very strong expression is seen in the dermal condensate with slightly less in the epidermal compartment of the follicle (Figure 4i). These results are generally consistent with the expression profiles of these genes during mouse primary hair follicle formation (Saxena et al., 2019).

The transcription factor SOX2 is a marker of the dermal condensate and dermal papilla of primary and secondary hair follicles, but not tertiary follicles, in mouse (Saxena et al., 2019). We identified SOX2 immunofluorescence in the dermal condensates of hair follicles in all areas of skin and all ages examined in human (Figure 4j), therefore SOX2 does not mark different follicle types in human as it does in mouse. SOX2 expression is uniformly distributed across the entirety of mouse dermal primary dermal condensates, whereas in human hair follicles SOX2 was expressed in the core of the dermal condensate with a layer of SOX2 negative cells at the periphery (Figure 4j and k, Supplementary Figure 3). When compared to developing mouse skin, the dermal condensates emerging under human hair follicle placodes appear to be more loosely arranged, and SOX2 was not detected in the very early stages of condensate development (Figure 4l and m, Supplementary Figure 4). Occasional SOX2 positive Merkel cells are observed in the interfollicular epidermis, appearing first in the facial regions at week 12 and the torso by week 16 (not shown).

### Comparison of gene expression in developing hair follicles of mouse and human fetal skin

To compare early human hair follicle development to that of the primary hair follicles in mouse, we analysed existing single nucleus RNA sequencing (snRNAseq) data of week 14 human back skin (Glover et al., 2023) and single cell RNA sequencing (scRNAseq) of E14.5 mouse back skin (Qu et al., 2022). Cell/nucleus clusters representing the placode, interfollicular epidermis (IFE), dermal condensate and general dermis were identified in both datasets. Expression of *BMP7* and *INHBA* identified the dermal condensates; *PTCH1* the general dermis; *EDAR* and *FGF20* the placode; and *KRT14* (without high *EDAR* or *FGF20*) the basal interfollicular epidermis (Figure 5a-d). Using the same thresholds as reported in Sulic et al. (Sulic et al., 2023) and Sennett et al. (Sennett et al., 2015) to define significantly upregulated genes (Fold change > 1.5, adjusted p-value <0.05), we identify many known markers of placodes and dermal condensates in both mouse and human. However, key markers such as *Shh* in the placode and *Sox2* in the dermal condensate were not found to be significantly upregulated in mouse placode or human dermal condensate using this simple threshold (Figure 5e-h). We therefore assessed whether genes were significantly up or down regulated in placode/condensates relative to the IFE/general dermis, taking into account both the level of expression and the proportion of cells in which the gene is expressed, which we refer to as uniqueness (U) score (Glover et al., 2023). We assessed genes with a U score >1.5 in both mouse and human, and identified 18 genes as placode enriched and 39 as dermal condensate enriched (Supplementary tables), with the top 5 enriched genes and two known markers shown in Figure 5i and j. All mouse placode enriched genes we identified in the mouse dataset were also identified as placode-upregulated in mouse by Sulic et al. and some by Sennett et al. (Sulic et al., 2023, Sennett et al., 2015), with many of those in the dermal condensate also identified by Sennett et al. (Sennett et al., 2015).

**Figure 5:**
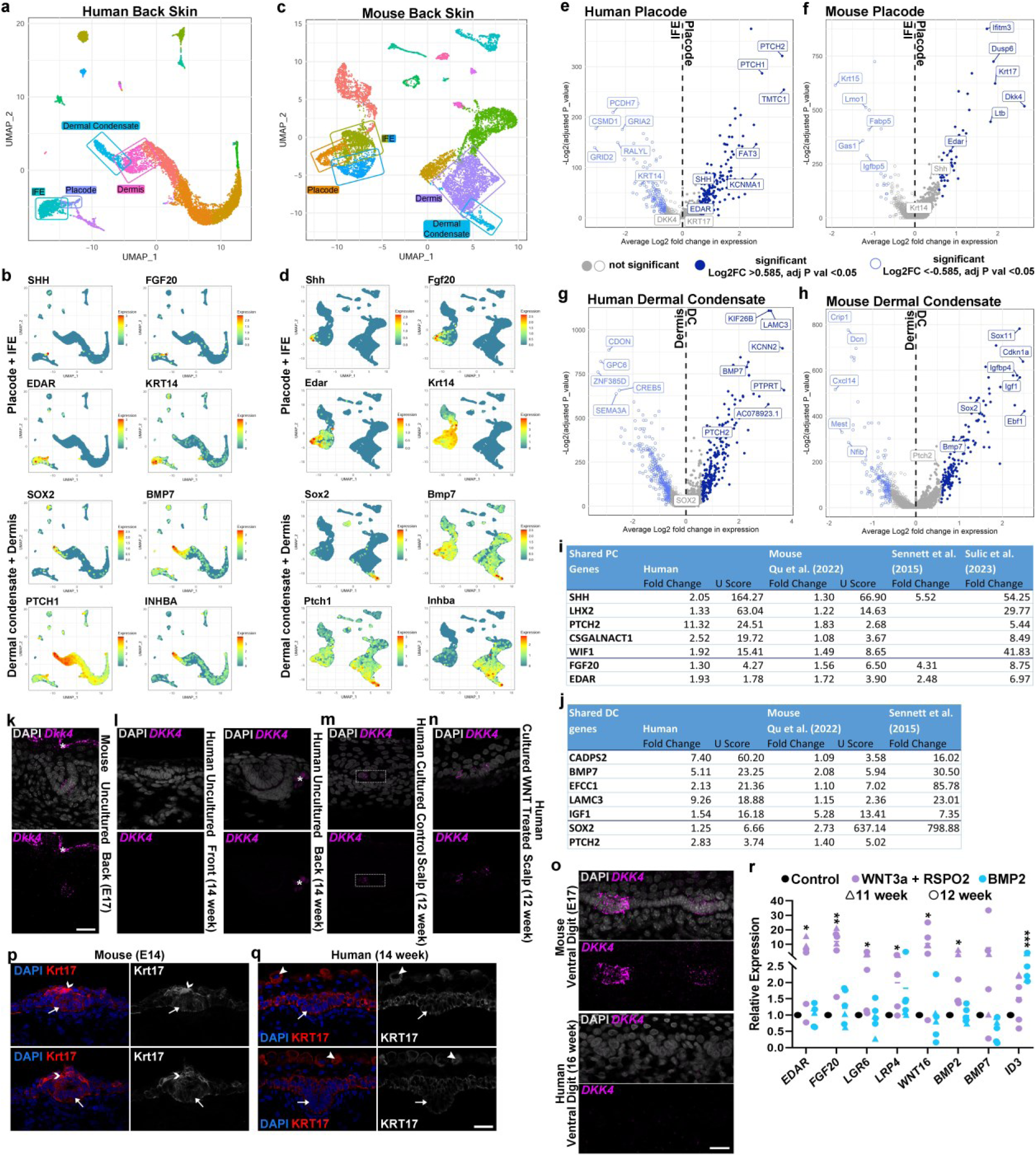
Transcriptome profiling of early mouse and human hair follicles. **(a)** Unbiased clustering of snRNAseq from human 14 week back skin. Identified clusters are placode, interfollicular epidermis (IFE), dermal condensate, and dermis. Here dermis is defined as those fibroblasts closest in expression profile to the dermal condensate (Supplementary Figure 5). **(b)** Feature plots from human data of signature genes used to identify the placode, IFE, dermal condensate, and dermis. **(c)** Unbiased clustering of scRNAseq from published E14.5 mouse data (Qu et al., 2022). Identified clusters are placode, IFE, dermal condensate, and dermis. Here dermis is defined as those fibroblasts closest in expression profile to the dermal condensate (Supplementary Figure 5). **(d)** Feature plots from mouse data of signature genes used to identify the placode, IFE, dermal condensate, and dermis. **(e-h)** Volcano plots of genes expressed in the epidermis and dermis of human and mouse. The top five genes upregulated in the placode/dermal condensate (dark blue) or downregulated (light blue), and known marker genes of placode, basal epidermis, and dermal condensate, are indicated. Significance thresholds correspond to the fold change >1.5 or <-1.5 and adjusted p-value <0.05. **(i and j)** Tables of enriched genes shared between mouse and human during early hair follicle development as determined by U score (U = avg_log2foldchange x proportion of detection in cluster1/proportion of detection in cluster2). Shared upregulated genes were defined as those with a U score > 1.5 in both species. **(i)** Genes with high expression in placodes compared to basal epidermal cells. Results for E14.5 mouse skin from other studies shown in right of table. **(j)** Genes with high expression in dermal condensate cells compared to general dermal cells. Results for E14.5 mouse skin from other studies shown in right of table. **(k-n)** RNA in situ hybridisation detecting *DKK4* transcripts in early mouse and human hair follicles; **(k)** a primary hair follicle from uncultured mouse embryonic day 17 (E17) back skin, **(l)** early hair follicles found on uncultured 14 week human back and front of torso skin, **(m)** an early placode on untreated cultured human 12 week scalp skin, and **(n)** an early placode on human 12 week scalp skin cultured with recombinant WNT3A and RSPO2. Asterisks indicate autofluorescent blood cells. **(o)** RNA in situ hybridisation to detect *DKK4* transcripts in dematoglyph ridges found on the ventral digit of E17 mouse and 16 week human volar skin. **(p-q)** Immunofluorescent detection of K17 in stage 2/3 placodes from **(p)** E14 mouse back skin, and **(q)** 14 week human back skin. Arrows indicate the basal epidermis, arrowheads indicate periderm cells, and chevrons indicate suprabasal cells above the basal placode. **(r)** RT-qPCR of hair follicle related genes in human back and scalp skin treated for 24 hours with recombinant WNT3A+RSPO2 or BMP2 proteins. Each point represents an individual piece of skin from back or scalp from 11 week (triangles) or 12 week (dots) embryos. *GAPDH* was used for normalisation. **\***p<0.05; **p<0.01; ***p<0.001. Student’s paired t-test. Scale bars: **k-n, p, q =** 25 μm; **o =** 20 μm

We identified genes specifically upregulated in the human placode and dermal condensate using a two-step process. First, with thresholds of U score >1.5 in the human and <0.5 in the mouse, and secondly removing any genes that had been found to be upregulated in the mouse placode or dermal condensate by Sulic et al. (2023) or Sennett et al. (2015). This approach identified genes that may be expressed specifically in human, but not mouse, early hair follicle development (Supplementary tables 2 and 5). However, the human skin analysed may carry hair follicle primordia that are more developmentally advanced than the tissue from which the mouse datasets were generated. Thus, further characterisation of this candidate gene set would be required to assess whether any might have species-specific expression patterns.

The observed differences in gene expression between mouse and human could be influenced by the different analyses used; the mouse study being conducted using single-cell RNAseq whereas the human study used single-nucleus RNAseq. Thus, we assessed by in situ hybridisation and immuofluorescent detection selected genes with notably different U scores between mouse and human, to assess the distinctiveness of their expression patterns.

*Dkk4* is a widely used marker of hair placode identity in mouse (Bazzi et al., 2007) and has been proposed to be a key regulator of hair follicle spacing (Sick et al., 2006). From snRNAseq using week 14 EGA back skin, we readily identified a placode cell cluster with high expression of known placode marker genes such as *EDAR*, *SHH*, and *FGF20* (Figure 5a and b, Supplementary tables). However, from a total of roughly 10,000 nuclei only a single read of the *DKK4* transcript was identified. This single detection event was in a mesenchymal cell cluster, while *Dkk4* expression reported in mouse skin is strictly epithelial (Bazzi et al., 2007). To assess the expression of *DKK4* in developing human skin in more detail, we performed RT-PCR experiments using freshly isolated, cultured and WNT-stimulated cultured human fetal skin. We found *DKK4* to be detectable by RT-PCR in all samples of human skin tested (Supplementary Figure 5). The majority of these samples (from 12 and 13 weeks back and scalp), were cultured skin maintained for 20 hours. Only one sample was uncultured skin (13 week front). When we used RNAscope in situ hybridisation to explore the localisation of *DKK4* expression, we found abundant expression in mouse hair placodes (Figure 5k) but detected essentially no signal in developing human epidermis or hair follicle placodes at any age when uncultured (Figure 5l). When human fetal scalp skin was cultured for 20 hours, a weak signal was detected in control (Figure 5m), with much higher expression in the sample treated with recombinant WNT3A and RSPO2 proteins (Figure 5n). Thus, in the absence of supplemented WNT stimulation, there was no detectable *DKK4* expression in intact native human fetal hair-bearing skin. Similar to the hair-bearing skin, *Dkk4* is also readily detected in mouse volar digit ridges, but not in human primary fingerprint ridges (Figure 5o).

Unlike the mouse scRNAseq, in human hair placodes the expression of *KRT14* was downregulated compared to that of the IFE (Figure 5e and f, Supplementary Figure 5) based on snRNAseq, and also apparent by immunofluorescent detection of the protein (Figure 4b). It is possible that the *KRT14* transcript and encoded protein have a long half-life, and the slower rate of human development may permit its disappearance from placodes more completely than in the faster-developing mouse skin. *KRT17* was also differentially expressed in mouse and human epidermal placodes (Figure 5e and f), with increased abundance in mouse placode vs. IFE but not in human placode vs. IFE. Immunofluorescent detection agreed with this finding, with significantly stronger K17 expression in the mouse placode and suprabasal cells above the placode than in the IFE, but very slightly higher expression in the IFE than the placode of human skin (Figure 5p-q).

In embryonic day 13 and 14 mouse skin *EDAR*, *FGF20* and *BMP2* have been reported to be rapidly upregulated by WNT signalling and downregulated by BMP signalling (Glover et al., 2017, Mou et al., 2006). We assessed the response of these genes to stimulation of the WNT pathway (using recombinant WNT3A + RSPO2 proteins) and BMP pathway (using recombinant human BMP2 protein). We find that WNT pathway stimulation increases abundance of *EDAR*, *FGF20* and *BMP2* transcripts on back and scalp skin at 11 and 12 weeks EGA, confirming conservation of WNT targets between species. We did not detect any decrease in EDAR expression elicited by BMP signalling, though the positive control for BMP treatment, *ID3*, did increase in abundance (Figure 5r). This suggests that, unlike the situation on mouse, *EDAR* is not a BMP target gene in developing human skin.

### EDA, EDAR expression and BMP and WNT activity in eccrine sweat gland development

Formation of eccrine sweat glands relies on EDAR signalling, causing hypohidrotic ectodermal dysplasia when mutations in *EDA*, *EDAR* or *EDARADD* abolish pathway function (Sadier et al., 2014). We assessed the expression of key components and transducers of the WNT, BMP and EDA/EDAR pathways in the development of human eccrine sweat glands of the volar skin. Sweat glands begin development on the volar skin (that of the ventral side of the hands and feet) from approximately 16 weeks EGA, budding off from the primary ridges. Morphologically, the eccrine sweat gland rudiments appear similar to an early hair follicle placode but are not associated with condensed dermal cells (Supplementary Figure 6). Like the hair follicles, the sweat glands show strong expression of LEF1, β-CATENIN, *EDAR*, and *BMP2* (Supplementary Figure 6), with the majority of these signals concentrated to the leading edge of the downgrowing sweat gland epithelial cord. Unlike hair follicles, there is very little expression of *FGF20*, increased pSMAD1/5, and no expression of *SHH* or SOX2 in developing sweat glands, at any stage of their development. Expression of *EDA* is very low in the downgrowing sweat gland bud (Supplementary Figure 6), unlike its prominent expression in the hair placode. Since EDA-EDAR signalling is required for sweat gland formation in mouse and human, and sustained EDA signalling is required to generate an eccrine sweat gland in mouse (Cui et al., 2009) it is possible that the extended growth of the sweat gland draws EDA ligand from non-epithelial sources.

## Discussion

This comparison of early human and mouse hair follicles reveals that their morphology, physical size, signalling pathway activity and gene expression are broadly comparable, supporting the use of mouse primary hair follicles as a model for human hair follicle development. The spacing and size of the hair follicle rudiments are strikingly similar between the species, suggesting similar mechanisms of embryonic pattern formation. However, the distinction between the large terminal and small vellus hair follicles in human is not apparent at early stages of hair follicle formation, and so appears to arise after their morphogenesis and the production of lanugo hair fibres. Expression of SOX2, which distinguishes the last-forming zigzag hair follicles from other types in mouse (Driskell et al., 2009) does not identify future vellus or terminal hair follicles in human, suggesting that these hair types in mouse do not have analogues in human skin. Thus, the study of early embryonic hair follicle formation in mouse appears to be limited as a model to understand the evolution of the human hair types and their anatomical distribution.

In human, profound hormonal regulation of hair follicles in some regions of the skin occurs throughout life, with axillary and beard hairs transitioning from vellus to terminal, and scalp hair follicles transitioning from terminal to vellus, under the influence of androgens (Randall, 2007). This illustrates the ability of human hair follicles to change size through hair cycles. It may be that the cellular processes driving follicle enlargement/minituarisation as a result of androgen stimulation from puberty are the same events that mediate the vellus-terminal transition during prenatal development (Whiting, 2001), though at this early stage of life operating without an androgen trigger.

Formation of feather buds in chicken skin has long been studied as a model for skin appendage development in general. In chicken skin a densely cellular dermis forms tracts, which will bear feathers, while loose dermis will not (Olivera-Martinez et al., 2004). Thus, the structure of the avian skin influences which regions will and will not be feather bearing. In mouse, however, the onset of primary hair follicle formation is not accompanied by an increase in dermal cell density (Makela and Mikkola, 2023), and we find that the density of dermal cells in human skin development does not closely align with the onset of hair follicle formation nor define areas with future terminal or vellus hair follicles. The closest observation to the feather tracts of avian embryos was the definition of a high density LEF1-positive dermis at the very first sites of hair follicle formation; the upper lip, chin and eyebrow, in human development. These could potentially reflect an influence of dermal structure and signalling on specific hair-bearing regions of human skin.

DKK4 has been proposed to be a central factor in defining hair follicle density and spacing (Sick et al., 2006), though no *Dkk4* mutant mouse hair patterning phenotype has yet been reported. In cats, a mutation in *DKK4* is associated with altered coat colour patterns, while the distribution of the hair follicles themselves is not altered by this mutation (Kaelin et al., 2021). In our investigations we did not detect *DKK4* expression in native human skin undergoing hair follicle development by in situ hybridisation or single nucleus RNA-sequencing, though we did detect the transcript by RT-PCR and upon WNT stimulation of cultured skin. Further technical assessment of *DKK4* expression in human skin is warranted, though the current findings suggest that this widely-employed placode marker gene in mouse may have species-specific expression in skin development.

*Edar* regulation by BMP signalling has also been proposed as a driver of primary hair follicle pattern formation and spacing in mouse (Mou et al., 2006). However, the key regulatory effect of BMPs in suppressing *Edar* expression that was observed in mouse we failed to detect in human, suggesting that this interaction is not relevant to human hair patterning. Thus the two simple models proposed for hair follicle patterning based on mouse models (Mou et al., 2006, Sick et al., 2006) do not appear to apply in human skin, suggesting that human hair follicle spacing either relies on decisive regulatory interactions not yet observed, or upon a complex interweaving of a set of regulatory pathways with multiple regulatory connections between them (Glover et al., 2017). The longer time taken to produce a pattern in human may permit the engagement of more signalling pathways and interactions in this process.

In conclusion, while there are some key differences to be recognised, the development of mouse primary hair follicles generally serves as a good model for human hair follicle development. However, an understanding of the evolution of human-specific hair characteristics and human ‘hairlessness’ needs to consider events taking place subsequent to initial hair follicle patterning and morphogenesis.

## Acknowledgements

We thank Richard Anderson for his help with the study.

## Materials and Methods

### Tissues

Human fetal tissue was obtained following elective termination with informed maternal consent from the Royal Infirmary of Edinburgh. Approval for the collection of tissue was granted by the Lothian Research Ethics Committee (ref: 08/S1101/1). Gestational age was estimated by an ultrasound scan and foot length measurements.

Maintenance of transgenic mouse lines was approved by the Roslin Institute Animal Welfare and Ethical Review Board and carried out under UK Home Office license (P682B81E4). TCF/Lef::H2B-GFP (Ferrer-Vaquer et al., 2010) mice were maintained on the FVB background. For timed matings noon on the day of detection of a copulatory plug was assigned E0.5.

### Sample Preparation and Histology

Dissected skin samples stored at 4°C for up to 24 hours were fixed overnight in 4% PFA or NBF, then dehydrated through an ethanol (EtOH) series from 25% to 100% with additional clearing steps in 1:1 chloroform: 100% EtOH and 100% chloroform. Samples were then incubated in paraffin at 65°C overnight and embedded in paraffin blocks. 6 μm sections were cut on a Thermo Microtome HM325 for histology and immunofluorescence. For histology, slides were dewaxed and stained for haematoxylin and eosin (H&E) using a Leica Autostainer XL and imaged on a Nikon Ni brightfield microscope or Hamamatsu Nanozoomer XR slide scanner. Whole unprocessed fixed samples were imaged with an Olympus SZX10 stereo microscope.

### Immunofluorescence

Paraffin sections on slides were dewaxed using a Leica Autostainer XL and immersed in citrate buffer for antigen retrieval using an Antigen Retriever 2100 (Aptum Biologics). Slides were washed with Tris-buffered saline (TBS) containing 0.1% Tween (TBST) and 0.1% Triton, and blocked for 1 hr at room temperature in 5% goat serum/TBST before primary antibodies were added and slides incubated overnight at 4°C. Sections were washed in TBST, then secondary antibody (1:500, Life Technologies A-11035, A-32723) applied for 1 hr at room temperature. Slides were washed twice in TBST and once in TBS (no detergent) then stained for 30 seconds with TrueBlack (Biotium). Slides were rinsed with TBS and counterstained with DAPI (1:5000, Sigma) for 3 minutes before being mounted with Prolong Gold (Life Technologies). Imaging was carried out on a Zeiss LSM 880 confocal microscope or Leica DMLB upright microscope.

For EdU staining the Click-iT™ EdU Cell Proliferation Kit for Imaging, Alexa Fluor™ 555 dye (Thermo Fisher) was used according to manufacturer’s instructions. Following incubation with the EdU cocktail, sections were washed with PBS/3% BSA before undergoing immunofluorescence performed as detailed above from the blocking step.

For wholemount immunofluorescence, fixed tissue samples were washed in PBS with 0.5% Triton X100 (PBTx) three times for 30 minutes at RT then incubated in 10% goat serum/PBTx for two hours at RT. Samples were incubated with primary antibodies (Table 1) in 10% goat serum/PBTx for 24 hours at 4°C. Samples were washed three times for 45 minutes in PBTx at RT before incubation with secondary antibodies and/or phalloidin (Table 1) in 10% goat serum/PBTx for 24 hours at 4°C. Samples were washed again three times for 45 mins in 10% goat serum/PBTx then incubated in DAPI/PBTx for 30 mins and mounted in Prolong Gold. Samples were imaged using a Zeiss LSM 880 microscope.

### In situ hybridisation

RNA in situ hybridisation on tissue sections was performed using the RNAscope Fluorescent Multiplex Assay V2 kit (ACD biosciences) following the manufacturer’s protocol. Hybridisation was done on 6 µm formalin fixed paraffin sections and detected using Opal dyes (Akoya Biosciences) (Table 1). After in situ hybridisation, samples were washed in TBST and immunofluorescence performed as described above, from the initial blocking step, but excluding TrueBlack treatment.

For RNA in situ hybridisation on whole, unsectioned, specimens, tissue was fixed in 4% PFA at 4°C for 48 hours then dehydrated through a methanol series and bleached with 5% H_2_O_2_ in 100% methanol. After rehydration into PBS + 0.1% Tween (PBT), samples were permeabilised with 20 μg/ml proteinase K for 20 minutes, washed with 2 mg/ml glycine, and post-fixed with 0.2% glutaraldehyde. Pre-hybridisation was carried out over 2 days at 65°C followed by hybridisation with DIG-labelled human *EDAR* riboprobe (consisting of the entire open reading frame; nucleotides 280-1626 of NCBI Reference Sequence NM_022336.4) for a further 2 days. Samples were washed, blocked in 10% heat inactivated sheep serum (HISS) and incubated at 4°C with anti-DIG antibody (1:2000, Roche) in TBST with 1% HISS overnight. Staining was detected with 5-bromo-4-chloro-3’-indolylphosphate/nitro-blue-tetrazolium (BCIP/NBT, Sigma).

### Measurement of morphological and histological features Quantification and statistics

Unless otherwise stated, Fiji software (Schindelin et al., 2012) was used for measurements and image analysis. Graphpad Prism 9 was used for statistical analysis and graph production. Statistical tests are named in the text and supplementary statistical information.

### Single nucleus RNA sequencing and data processing

snRNAseq of the human 14 week back skin samples and the data processing of both the human and mouse datasets was carried out as in Glover et al. (Glover et al., 2023)

Human 14 week back skin snRNAseq data GEO: GSE195657

Mouse E14.5 back skin scRNAseq data GEO: GSE198487

A U score = avg_log2FC x pct.1/pct.2 was used to determine genes that were up/downregulated in the placode and dermal condensate, while U score = avg_log2FC x pct.2/pct.1 was used for the IFE and dermis, where pct.1 and pct.2 refer to the percentage of cells in cluster 1 and cluster 2 with detection of the transcript.

### Placode Measurements

Images from wholemount *EDAR* in situ hybridisations, P-CADHERIN wholemount immunofluorescence, and H&E stained tissue sections were analysed using ImagePro Plus 6.2, Fiji, Zen 2 (Blue edition), and Hamamatsu NDP.view 2 software to measure placode diameter and density. *EDAR* wholemount in situ hybridisation images were processed with a linear local equalisation filter to remove background signal before measurement.

Mouse measurements were taken from E14.5 TCF/Lef::H2B-GFP (Ferrer-Vaquer et al., 2010) mouse back skins imaged on a Zeiss Axiozoom V16 microscope, or H&E stained sections.

### Cell density and proliferation

Cell counts were carried out using Fiji software on immunofluorescent images of sectioned skin stained with Ki67 and counterstained with DAPI. Images were taken using Zeiss LSM 880 Confocal, Zeiss LSM 710 Confocal, or Leica DMLB upright fluorescent microscopes. Images were taken at x20 magnification and scaled appropriately. Counts were taken from the entire field of view encompassing 300-600 μm linear skin length. Damaged tissue or hair follicle structures were disregarded.

To assess dermal cell density, DAPI and Ki67 positive nuclei were counted to a depth of 40 μm from the basal epidermis. Epidermal counts of DAPI and Ki67 positive nuclei were taken from the basal layer only.

### Skin culture and quantitative RT-PCR

Dissected skin samples were incubated on methylcellulose filters (Millipore) at 37°C in DMEM containing 5% FBS and either control (vehicle), recombinant human WNT3A (250 ng/ml, R&D Systems) + recombinant human RSPO2 (500 ng/ml, R&D Systems), or recombinant human/mouse/rat BMP2 (500 ng/ml, R&D Systems). Samples were collected after 24 hours and RNA was extracted using TriReagent (Sigma-Aldrich) according to manufacturer’s instructions. cDNA was synthesised using Superscript III Reverse Transcriptase (Invitrogen) with random primers. All qRT-PCR reactions were carried out in 20 μl using SYBR Green Universal Master Mix (Roche) including Rox reference dye. Each reaction was performed in triplicate.

### Oligonucleotide sequences for quantitative RT-PCR

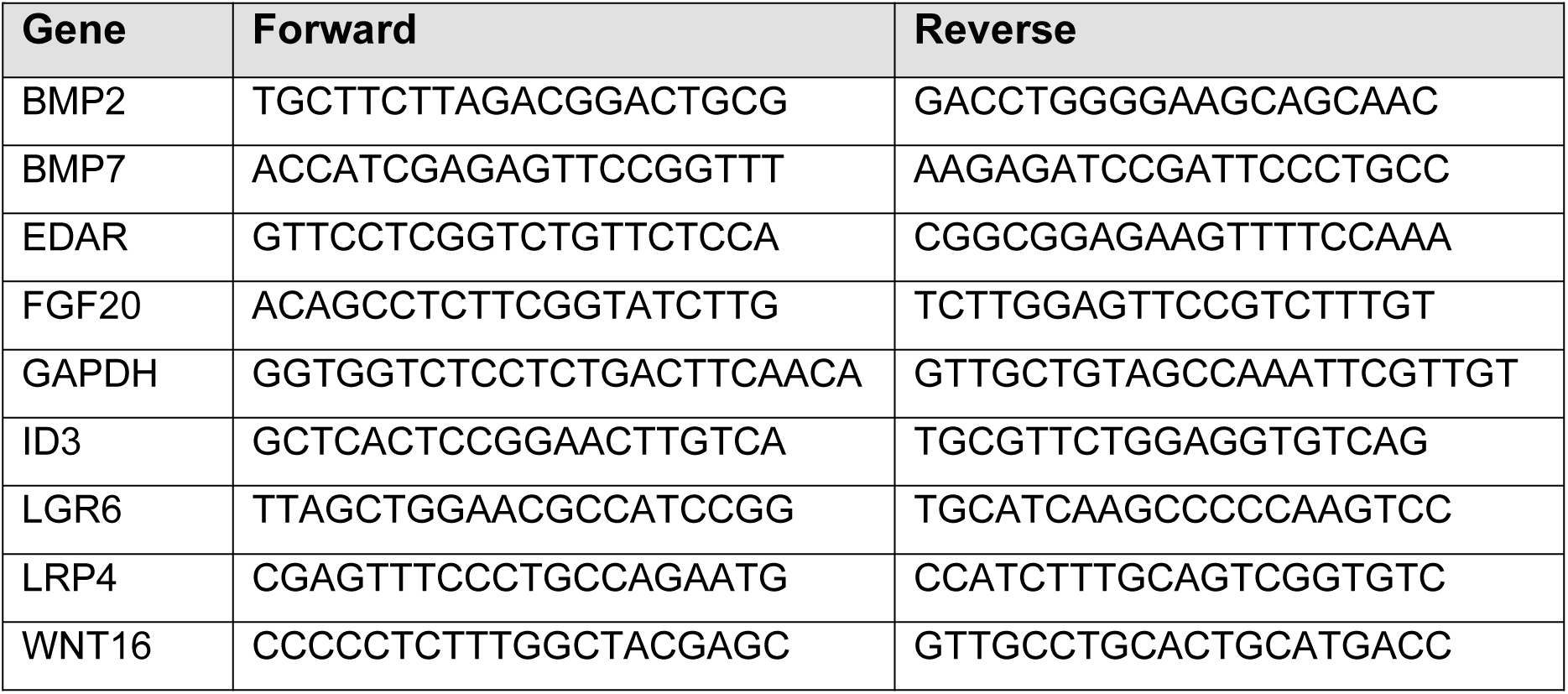

### Reagents used for staining tissues

| Reagent | Species | Supplier | Cat. No. | Dilution | Retrieval |
| --- | --- | --- | --- | --- | --- |
| **Primary antibodies** |  |  |  |  |  |
| Anti-Cytokeratin 14 antibody [LL002] | Mouse | Abcam | ab7800 | 1:1000 | Citrate |
| Recombinant Anti-Ki67 antibody [SP6] | Rabbit | Abcam | ab16667 | 1:200 | Citrate |
| Anti-Cytokeratin 10 antibody [RKSE60] | Mouse | Abcam | ab9025 | 1:600 | Citrate |
| Recombinant Anti-beta catenin antibody [E247] | Rabbit | Abcam | ab32572 | 1:500 | Citrate |
| Anti-α E-catenin Antibody (G-11) | Mouse | Santa Cruz | sc-9988 | 1:100 | Citrate |
| Recombinant Anti-LEF1 antibody [EPR2029Y] | Rabbit | Abcam | ab137872 | 1:500 | Citrate |
| Phospho-SMAD1/SMAD5 (Ser463, Ser465)<br>Recombinant Rabbit Monoclonal Antibody (31H14L11) | Rabbit | Thermo Fisher | 700047 | 1:200 | Citrate |
| Anti-SOX2 antibody | Rabbit | Abcam | ab97959 | 1:500 | Citrate |
| Anti-Cytokeratin 8 antibody | Rabbit | Abcam | ab59400 | 1:200 | Citrate |
| P-cadherin Polyclonal Antibody | Rabbit | Invitrogen |  | 1:1000 | Citrate |
| **Secondary antibodies, dyes and detection reagents** |  |  |  |  |  |
| Goat anti-Rabbit IgG (H+L) Highly Cross-Adsorbed Secondary Antibody, Alexa Fluor 546 | Goat | Thermo Fisher | A11035 | 1:500 |  |
| Goat anti-Rabbit IgG (H+L) Highly Cross-Adsorbed Secondary Antibody, Alexa Fluor 647 | Goat | Thermo Fisher | A21245 | 1:500 |  |
| Goat anti-Mouse IgG (H+L)<br>Highly Cross-Adsorbed<br>Secondary Antibody, Alexa<br>Fluor 488 | Goat | Thermo<br>Fisher | A11029 | 1:500 |  |
| 4',6-Diamidino-2-phenylindole<br>dihydrochloride (DAPI) |  | Sigma-Aldrich | D9542 | 1:5000 |  |
| Alexa Fluor™ 647<br>Phalloidin |  | Thermo<br>Fisher | A22287 | 1:200 |  |
| Opal Dye 520 |  | Akoya<br>biosciences | OP-001001 | 1:1500 |  |
| Opal Dye 570 |  | Akoya<br>biosciences | OP-001003 | 1:1500 |  |
| Opal Dye 620 |  | Akoya<br>biosciences | OP-001004 | 1:1500 |  |
| Opal Dye 690 |  | Akoya<br>biosciences | OP-001006 | 1:1500 |  |
| Click-iT™ EdU Cell<br>Proliferation Kit for Imaging,<br>Alexa Fluor™ 555 dye |  | Invitrogen | C10338 |  |  |
| **RNAscope probes** |  |  |  |  |  |
| Hs-BMP2 |  | ACD | 430641 |  |  |
| Hs-FGF20-C2 |  | ACD | 460511-C2 |  |  |
| Hs-EDAR |  | ACD | 494601 |  |  |
| Hs-DKK1 |  | ACD | 421411 |  |  |
| Hs-DKK2-C2 |  | ACD | 531131-C2 |  |  |
| Hs-SHH-C2 |  | ACD | 600951-C2 |  |  |
| Hs-EDA-C2 |  | ACD | 535971-C2 |  |  |
| Hs-DKK4-C3 |  | ACD | 429371-C3 |  |  |
| Hs-BMP7 |  | ACD | 424401 |  |  |
| Mm-DKK4 |  | ACD | 404851 |  |  |
| Hs-DKK4 |  | ACD | 429371 |  |  |

## References

Bazzi, H., Fantauzzo, K. A., Richardson, G. D., Jahoda, C. A. & Christiano, A. M. 2007. The Wnt inhibitor, Dickkopf 4, is induced by canonical Wnt signaling during ectodermal appendage morphogenesis. Dev Biol, 305, 498–507.

Biggs, L. C., Makela, O. J., Myllymaki, S. M., DAS Roy, R., Narhi, K., Pispa, J., Mustonen, T. & Mikkola, M. L. 2018. Hair follicle dermal condensation forms via Fgf20 primed cell cycle exit, cell motility, and aggregation. Elife, 7.

Botchkarev, V. A., Botchkareva, N. V., Roth, W., Nakamura, M., Chen, L. H., Herzog, W., Lindner, G., Mcmahon, J. A., Peters, C., Lauster, R., Mcmahon, A. P. & Paus, R. 1999. Noggin is a mesenchymally derived stimulator of hair-follicle induction. Nat Cell Biol, 1, 158–64.

Botchkarev, V. A., Botchkareva, N. V., Sharov, A. A., Funa, K., Huber, O. & Gilchrest, B. A. 2002. Modulation of BMP signaling by noggin is required for induction of the secondary (nontylotrich) hair follicles. J Invest Dermatol, 118, 3–10.

Botchkarev, V. A. & Paus, R. 2003. Molecular biology of hair morphogenesis: development and cycling. J Exp Zool B Mol Dev Evol, 298, 164–80.

Cheng, C. W., Niu, B., Warren, M., Pevny, L. H., LOVELL-Badge, R., Hwa, T. & Cheah, K. S. 2014. Predicting the spatiotemporal dynamics of hair follicle patterns in the developing mouse. Proc Natl Acad Sci U S A, 111, 2596–601.

Cui, C. Y., Kunisada, M., Esibizione, D., Douglass, E. G. & Schlessinger, D. 2009. Analysis of the temporal requirement for eda in hair and sweat gland development. J Invest Dermatol, 129, 984–93.

Domagala, Z., Dabrowski, P., Kurlej, W., Porwolik, M., Wozniak, S., Kacala, R. R. & Gworys, B. 2017. The sequence of lanugo pattern development on the trunk wall in human fetuses. Adv Clin Exp Med, 26, 967–972.

Driskell, R. R., Giangreco, A., Jensen, K. B., Mulder, K. W. & Watt, F. M. 2009. Sox2-positive dermal papilla cells specify hair follicle type in mammalian epidermis. Development, 136, 2815–23.

Dry, F. W. 1926. The coat of the mouse (Mus musculus). Journal of Genetics, 16, 287–340.

Duverger, O. & Morasso, M. I. 2009. Epidermal Patterning and Induction of Different Hair Types During Mouse Embryonic Development. Birth Defects Research Part C-Embryo Today-Reviews, 87, 263–272.

FERRER-Vaquer, A., Piliszek, A., Tian, G., Aho, R. J., Dufort, D. & Hadjantonakis, A. K. 2010. A sensitive and bright single-cell resolution live imaging reporter of Wnt/ss-catenin signaling in the mouse. BMC Dev Biol, 10, 121.

Glover, J. D., Sudderick, Z. R., Shih, B. B., BATHO-Samblas, C., Charlton, L., Krause, A. L., Anderson, C., Riddell, J., Balic, A., Li, J., Klika, V., Woolley, T. E., Gaffney, E. A., Corsinotti, A., Anderson, R. A., Johnston, L. J., Brown, S. J., Wang, S., Chen, Y., Crichton, M. L. & Headon, D. J. 2023. The developmental basis of fingerprint pattern formation and variation. Cell, 186, 940–956 e20.

Glover, J. D., Wells, K. L., Matthaus, F., Painter, K. J., Ho, W., Riddell, J., Johansson, J. A., Ford, M. J., Jahoda, C. A. B., Klika, V., Mort, R. L. & Headon, D. J. 2017. Hierarchical patterning modes orchestrate hair follicle morphogenesis. PLoS Biol, 15, e2002117.

Gupta, K., Levinsohn, J., Linderman, G., Chen, D., Sun, T. Y., Dong, D., Taketo, M. M., Bosenberg, M., Kluger, Y., Choate, K. & Myung, P. 2019. Single-Cell Analysis Reveals a Hair Follicle Dermal Niche Molecular Differentiation Trajectory that Begins Prior to Morphogenesis. Dev Cell, 48, 17–31 e6.

Headington, J. T. 1984. Transverse microscopic anatomy of the human scalp. A basis for a morphometric approach to disorders of the hair follicle. Arch Dermatol, 120, 449–56.

Headon, D. J. & Overbeek, P. A. 1999. Involvement of a novel Tnf receptor homologue in hair follicle induction. Nat Genet, 22, 370–4.

Ho, W. K. W., Freem, L., Zhao, D., Painter, K. J., Woolley, T. E., Gaffney, E. A., Mcgrew, M. J., Tzika, A., Milinkovitch, M. C., Schneider, P., Drusko, A., Matthaus, F., Glover, J. D., Wells, K. L., Johansson, J. A., Davey, M. G., Sang, H. M., Clinton, M. & Headon, D. J. 2019. Feather arrays are patterned by interacting signalling and cell density waves. PLoS Biol, 17, e3000132.

Huh, S. H., Narhi, K., Lindfors, P. H., Haara, O., Yang, L., Ornitz, D. M. & Mikkola, M. L. 2013. Fgf20 governs formation of primary and secondary dermal condensations in developing hair follicles. Genes Dev, 27, 450–8.

Jahoda, C. A. 1992. Induction of follicle formation and hair growth by vibrissa dermal papillae implanted into rat ear wounds: vibrissa-type fibres are specified. Development, 115, 1103–9.

Kaelin, C. B., Mcgowan, K. A. & Barsh, G. S. 2021. Developmental genetics of color pattern establishment in cats. Nat Commun, 12, 5127.

Laurikkala, J., Pispa, J., Jung, H. S., Nieminen, P., Mikkola, M., Wang, X., SAARIALHO-Kere, U., Galceran, J., Grosschedl, R. & Thesleff, I. 2002. Regulation of hair follicle development by the TNF signal ectodysplasin and its receptor Edar. Development, 129, 2541–53.

Makela, O. J. M. & Mikkola, M. L. 2023. Mesenchyme governs hair follicle induction. Development, 150.

Mann, S. J. 1962. Prenatal formation of hair follicle types. Anatomical Record, 144, 135–141.

Mou, C., Jackson, B., Schneider, P., Overbeek, P. A. & Headon, D. J. 2006. Generation of the primary hair follicle pattern. Proc Natl Acad Sci U S A, 103, 9075–80.

Mou, C., Thomason, H. A., Willan, P. M., Clowes, C., Harris, W. E., Drew, C. F., Dixon, J., Dixon, M. J. & Headon, D. J. 2008. Enhanced ectodysplasin-A receptor (EDAR) signaling alters multiple fiber characteristics to produce the East Asian hair form. Hum Mutat, 29, 1405–11.

Olivera-Martinez, I., Viallet, J. P., Michon, F., Pearton, D. J. & Dhouailly, D. 2004. The different steps of skin formation in vertebrates. Int J Dev Biol, 48, 107–15.

Painter, K. J., Ptashnyk, M. & Headon, D. J. 2021. Systems for intricate patterning of the vertebrate anatomy. Philos Trans A Math Phys Eng Sci, 379, 20200270.

Paus, R., MULLER-Rover, S., VAN DER Veen, C., Maurer, M., Eichmuller, S., Ling, G., Hofmann, U., Foitzik, K., Mecklenburg, L. & Handjiski, B. 1999. A comprehensive guide for the recognition and classification of distinct stages of hair follicle morphogenesis. J Invest Dermatol, 113, 523–32.

Qu, R., Gupta, K., Dong, D., Jiang, Y., Landa, B., Saez, C., Strickland, G., Levinsohn, J., Weng, P. L., Taketo, M. M., Kluger, Y. & Myung, P. 2022. Decomposing a deterministic path to mesenchymal niche formation by two intersecting morphogen gradients. Dev Cell, 57, 1053–1067 e5.

Randall, V. A. 2007. Hormonal regulation of hair follicles exhibits a biological paradox. Semin Cell Dev Biol, 18, 274–85.

Sadier, A., Viriot, L., Pantalacci, S. & Laudet, V. 2014. The ectodysplasin pathway: from diseases to adaptations. Trends Genet, 30, 24–31.

Saxena, N., Mok, K. W. & Rendl, M. 2019. An updated classification of hair follicle morphogenesis. Exp Dermatol, 28, 332–344.

Schneider, M. R., SCHMIDT-Ullrich, R. & Paus, R. 2009. The hair follicle as a dynamic miniorgan. Curr Biol, 19, R132–42.

Sennett, R., Wang, Z., Rezza, A., Grisanti, L., Roitershtein, N., Sicchio, C., Mok, K. W., Heitman, N. J., Clavel, C., MA’ayan, A. & Rendl, M. 2015. An Integrated Transcriptome Atlas of Embryonic Hair Follicle Progenitors, Their Niche, and the Developing Skin. Dev Cell, 34, 577–91.

Sick, S., Reinker, S., Timmer, J. & Schlake, T. 2006. WNT and DKK determine hair follicle spacing through a reaction-diffusion mechanism. Science, 314, 1447–50.

Sulic, A. M., DAS Roy, R., Papagno, V., Lan, Q., Saikkonen, R., Jernvall, J., Thesleff, I. & Mikkola, M. L. 2023. Transcriptomic landscape of early hair follicle and epidermal development. Cell Rep, 42, 112643.

Tsatsou, F. & Zouboulis, C. C. 2014. Anatomy of the sebaceous gland. In: Zouboulis, C. C., Katsambas, A. & Kligman, A. (eds.) Pathogenesis and Treatment of Acne and Rosacea. Berlin, Heidelberg: Springer.

VAN Genderen, C., Okamura, R. M., Farinas, I., Quo, R. G., Parslow, T. G., Bruhn, L. & Grosschedl, R. 1994. Development of Several Organs That Require Inductive Epithelial-Mesenchymal Interactions Is Impaired in Lef-1-Deficient Mice. Genes & Development, 8, 2691–2703.

VAN Scott, E. J. & Ekel, T. M. 1958. Geometric relationships between the matrix of the hair bulb and its dermal papilla in normal and alopecic scalp. J Invest Dermatol, 31, 281–7.

Verhave, B. L., Nassereddin, A. & Lappin, S. L. 2022. Embryology, Lanugo. StatPearls. Treasure Island (FL).

Whiting, D. A. 2001. Possible mechanisms of miniaturization during androgenetic alopecia or pattern hair loss. J Am Acad Dermatol, 45, S81–6.

