## Supplemental figures and tables for "Characterisation of human hair follicle development"

### Supplementary figures

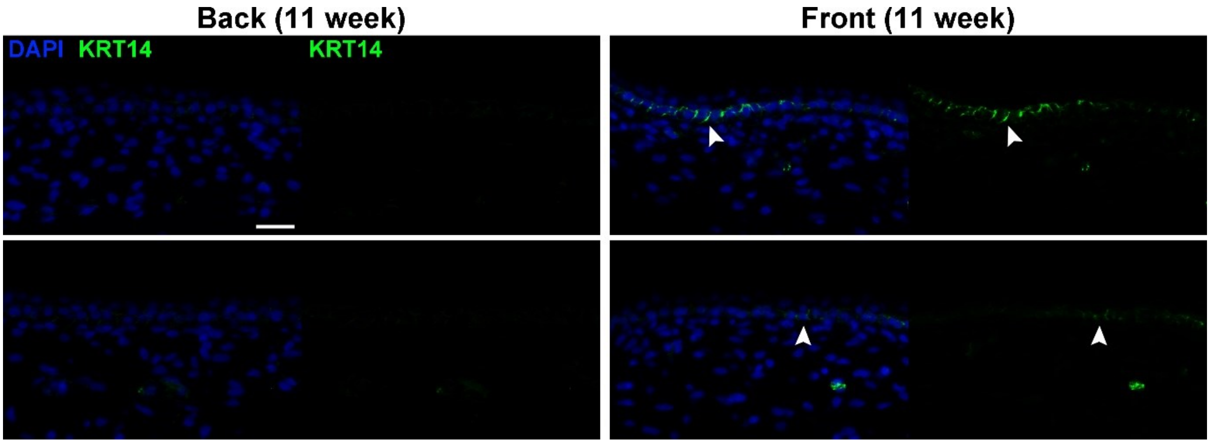

**Supplementary Figure 1: Keratin 14 is expressed in ventral skin prior to its expression dorsally.** Immunofluorescent detection of Keratin 14 in 11 week human torso skin. No expression was detected in the epidermis of the back skin but was detected in the front skin at this stage of development. Scale bar = 25 µm

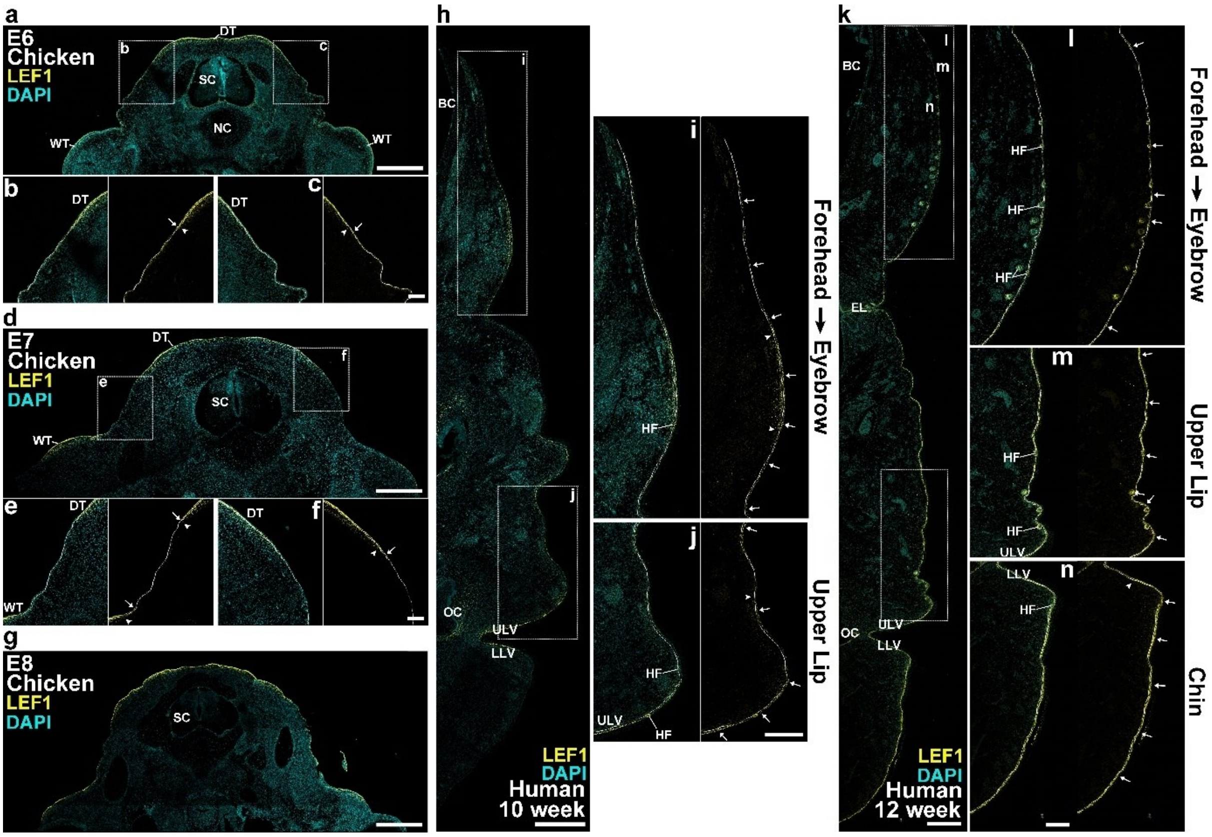

**Supplementary Figure 2: LEF1 detection in chicken feather tracts and the developing human facial skin.** Immunofluorescent detection of LEF1, counterstained with DAPI, in transverse sections of chicken embryos and saggital sections of human facial skin at the embryonic days or weeks indicated. LEF1 is detected in the chicken dorsal and wing skin tracts at **a-c)** embryonic day 6 (E6), **d-f)** E7, and **g)** E8, and in human fetal facial skin at **h-j)** 10 weeks EGA, and **k-n)** 12 weeks EGA. **a-g)** Arrows and arrowheads indicate boundaries of epidermal and mesenchymal LEF1 expression, denoting the boundaries of the presumptive feather tracts in these regions of the chicken embryos. **h-n)** Areas of epidermal LEF1 expression are indicated by arrows and strong mesenchymal LEF1 detection by arrowheads. In chicken embryonic skin strong LEF1 expression is present in the epidermis at the site of presumptive feather tracts with strong mesenchymal LEF1 in the upper dermis in the dorsal and humeral feather tracts. expression in the space between feather tracts is absent from the epidermis. **e & f)** Enlarged image of the gap between the dorsal and wing tracts. No epidermal LEF1 is seen in the gap, and only weak mesenchymal LEF1 is detected. Mesenchymal LEF1 in the tracts is

In human facial skin at week 10 epidermal LEF1 is present across the face particularly in the presumptive eyebrow, upper lip, and lip vermilion. Low mesenchymal LEF1 is observed generally with stronger expression in the dermis underneath the areas of highest epidermal expression. Strong epidermal and mesenchymal LEF1 is present in the eyebrow, with limited epidermal and no mesenchymal LEF1 continuing in the forehead.

At 12 weeks LEF1 expression in the epidermis is detected across the face with strong mesenchymal expression detected in the upper lip, and chin surrounding developing hair follicles. Dermal LEF1 appears weaker than that at 10 weeks but extends deeper into the dermis. Epidermal LEF1 is strong in the eyebrow and all hair follicles, but is slightly weaker in the forehead. Strong mesenchymal LEF1 expression is present in the upper lip, reducing to moderate in the cheek. DT = Dorsal tract, WT = Wing tract, FF = Feather follicle, SC = Spinal cord, NC = Notochord, BC = Brain cavity, OC = Oral cavity, EL = Eyelid, LLV = Lower lip vermilion, ULV = Upper lip vermilion, HF = Hair follicle. Scale bars: **a**, **d**, **g**, **h**, & **k** = 500 μm; **b**, **c**, **e**, & **f** = 100 μm; **i**, **j**, **l**, **m**, & **n** = 250 μm

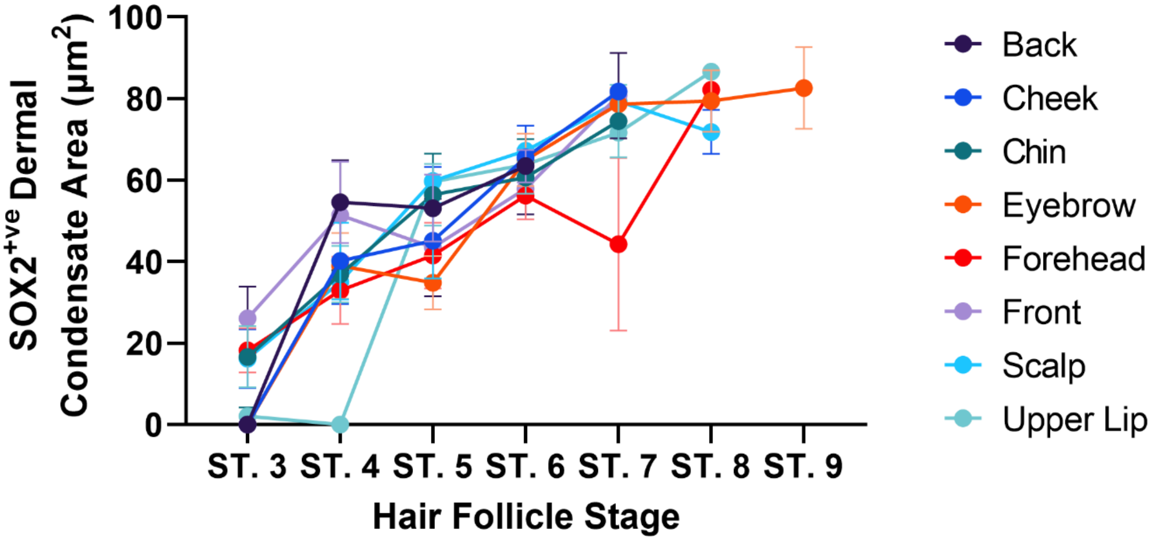

**Supplementary Figure 3: Proportion of SOX2+ve dermal condensate cells in developing human hair follicles.** The percentage of the dermal condensate that is SOX2+ve by area. All anatomical sites had SOX2+ve cells in the dermal condensate but in no hair follicle examined did all of the dermal condensate cells express detectable SOX2. A total of 258 hair follicles were measured from 15 individual specimens. Dots indicate the mean ± SEM at each anatomical site and hair follicle stage.

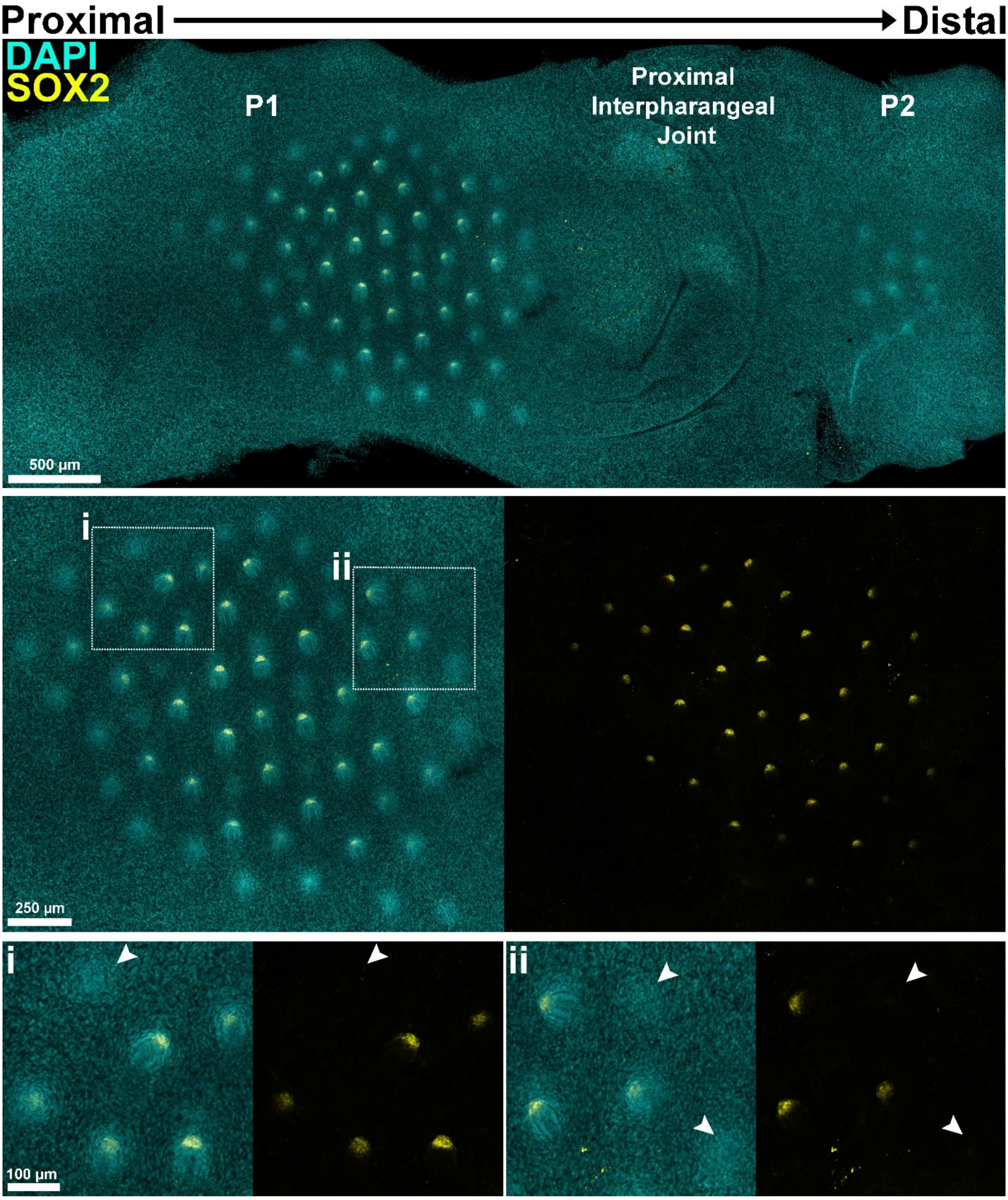

**Supplementary Figure 4: SOX2 expression is not detected at the earliest stages of human hair follicle morphogenesis.** Whole mount immunofluorescent detection of SOX2 in dorsal digit skin at 14 weeks. Various developmental stages of hair follicles are present on the skin above phalanx 1 (P1), in a cluster between the knuckle joints. The hair follicle primordia have a radial organisation, with the most advanced and deepest primordia at the centre of the cluster, giving way to progressively less developed primordia towards its periphery. From this arrangement we infer that hair follicle formation in this region is likely to occur in a spreading wave. SOX2 expression is observed in the dermal condensates of the more mature follicle primordia, but morphologically distinct mesenchymal condensations at the periphery of the cluster, and above P2, lack SOX2 immunoreactivity, demonstrating the late activation of its expression in human hair follicle development. Boxes **i** and **ii** are expanded in the lower panels. Arrowheads indicate dermal condensations of incipient follicles that are detectable by cell arrangement (DAPI stain) but that lack detectable SOX2 expression. Scale bars as indicated.

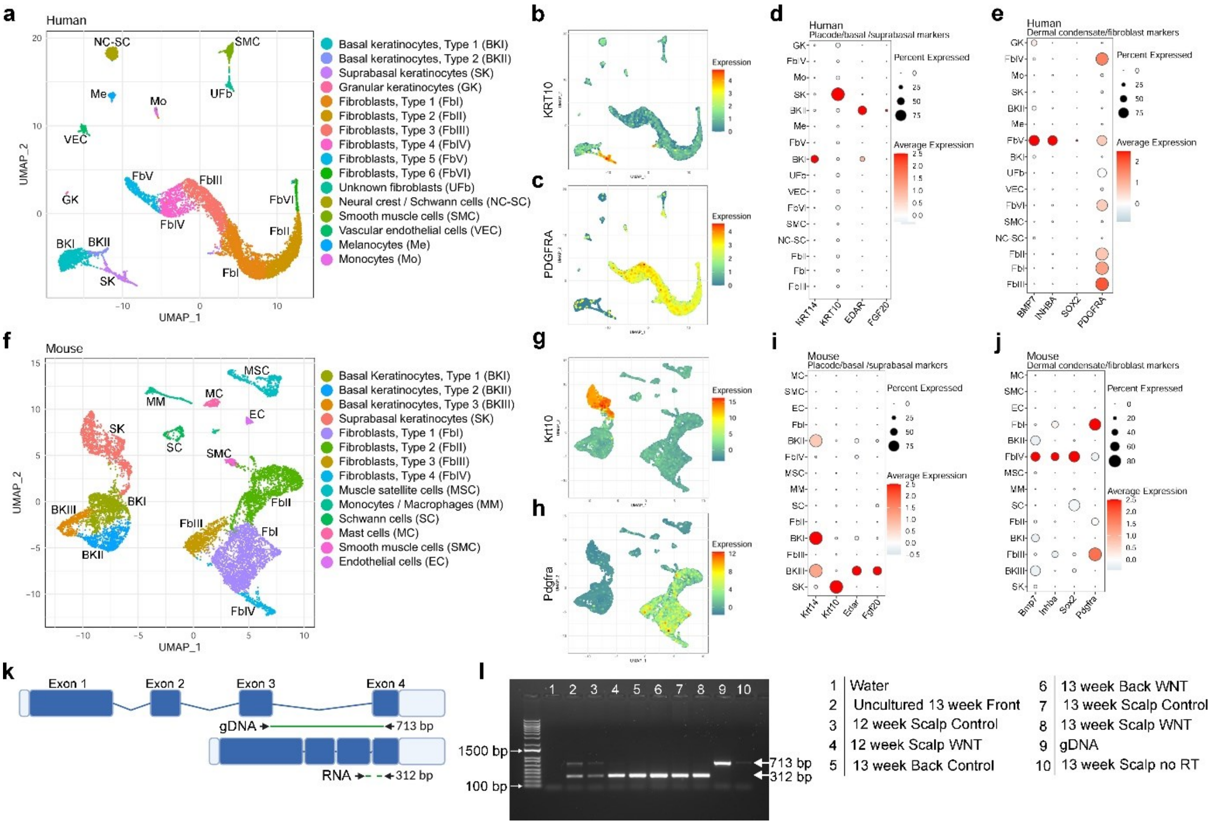

**Supplementary Figure 5: Gene expression cluster identification in snRNASeq and scRNASeq data, and *DKK4* transcript detection. a)** Unbiased clustering of snRNAseq from human 14 week back skin. All clusters are labelled based on differential gene expression and previously reported markers of different cell types. Populations for comparison were defined as follows: BKII = Placode, BKI = IFE, FbV = Dermal condensate, FbIV = Dermis. **b)** Feature plot for *KRT10* on human back skin identifying the suprabasal keratinocyte population. **c)** Feature plot for *PDGFRA* on human back skin identifying fibroblast populations. **d)** Dotplot of placode and epithelial cell markers from human back skin. **e)** Dotplot of dermal condensate and fibroblast markers in human back skin. **f)** Unbiased clustering of scRNAseq from published E14.5 mouse data (Qu et al., 2022). All clusters are labelled based on differential gene expression and known markers of cell types. Populations for comparison were defined as follows: BKIII = Placode, BKI + BKII = IFE, FbIV = Dermal condensate, FbI = Dermis. **g)** Feature plot for *Krt10* on mouse back skin identifying the suprabasal keratinocyte population. **h)** Feature plot for *Pdgfra* on mouse back skin identifying fibroblast populations. **i)** Dotplot of placode and epithelial cell markers from mouse back skin. **j)** Dotplot of dermal condensate and fibroblast markers in mouse back skin. **k)** Diagram of *DKK4* gene with primers used for PCR. Length of the gDNA fragment is shown with a solid green line, length of the RNA fragment with a dashed green line. **I)** PCR for *DKK4* on different ages and areas of human fetal skin. *DKK4* cDNA was amplified in all skin samples, both treated with recombinant WNT protein, and untreated. Only *DKK4* DNA was identified in the gDNA and no RT (no reverse transcriptase) reactions, as expected.

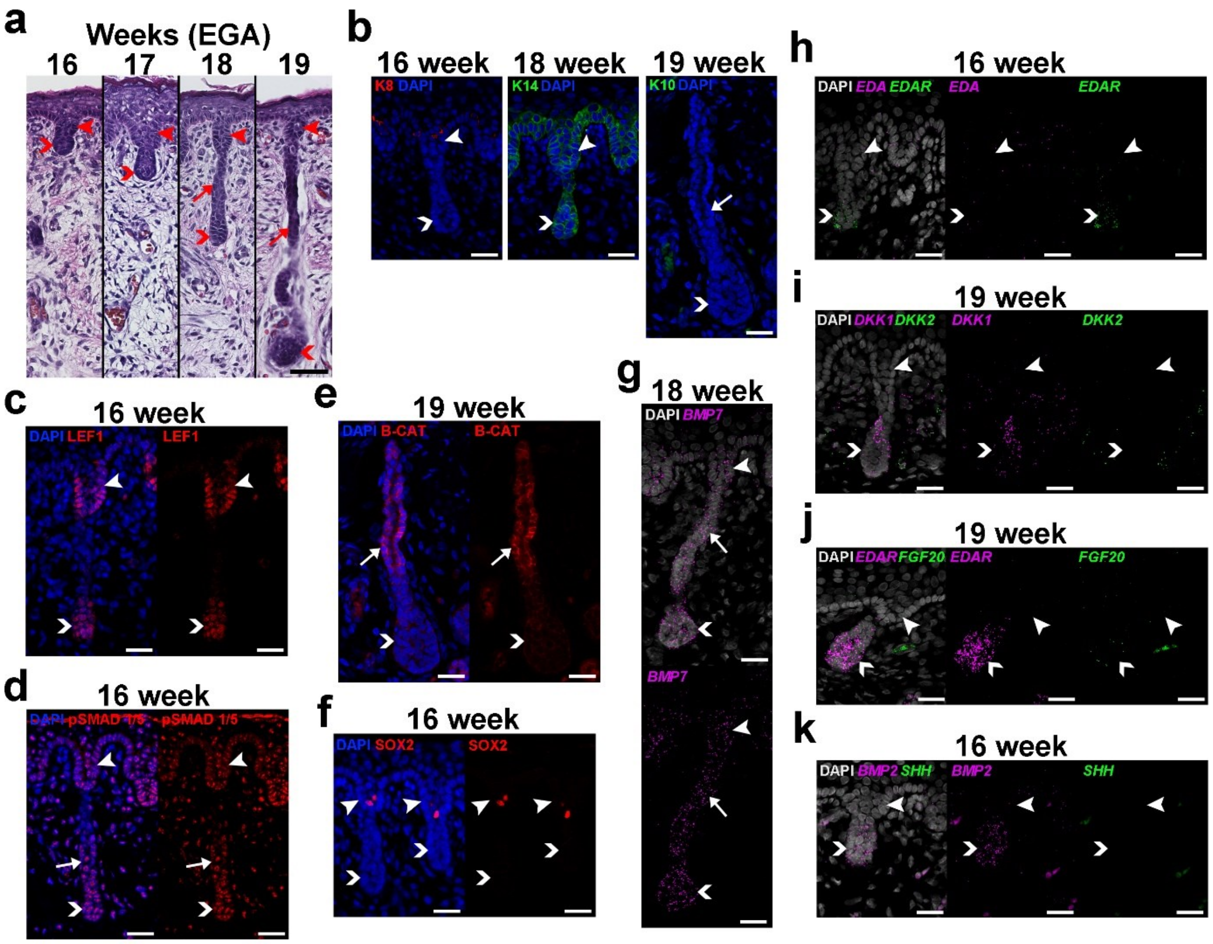

**Supplementary Figure 6: Expression of cell signalling factors during development of volar sweat glands. (a)** H&E stained sections showing sweat gland development on the volar skin from 16 to 19 weeks EGA. Sweat glands bud off from the primary fingerprint/dermatoglyph ridges and grow down as thin epithelial cords with a wider bulb at the leading edge and no overt dermal component. **(b-f)** Immunofluorescent detection of **(b)** Keratins 8, 14, and 10, **(c)** LEF1, **(d)** phospho-SMAD1/5, **(e)** β-catenin, and **(f)** SOX2 in developing sweat glands. **(g-k)** RNA in situ hybridisation to detect expression of **(g)** *BMP7*, **(h)** *EDA* and *EDAR*, **(i)** *DKK1* and *DKK2*, **(j)** *EDAR* and *FGF20*, and **(k)** *BMP2* and *SHH* in sweat glands of the volar skin. Arrowheads indicate primary fingerprint ridges, arrows indicate sweat gland ducts, and chevrons indicate sweat gland bulbs. Scale bars **a** = 50 μm; **b-k** = 25 μm

### Supplementary Tables

**Supplementary Table 1: Shared Placode Enriched Genes. U score = avg_log2FC x pct.1/pct.2. U score > 1.5 for mouse and human.**

| | Human | | | | | | Mouse (Qu et al., 2022) | | | | | | Sennett et al. (2015) | Sulic et al. (2023) |
|---|---|---|---|---|---|---|---|---|---|---|---|---|---|---|
| | p_val | avg_log2FC | pct.1 | pct.2 | p_val_adj | UScore | p_val | avg_log2FC | pct.1 | pct.2 | p_val_adj | UScore | log2FC | log2FC |
| SHH | 3.89E-23 | 1.03 | 0.16 | 0.001 | 1.14E-18 | 164.27 | 1.0251E-60 | 0.38 | 0.18 | 0.001 | 1.8684E-56 | 66.90 | | 1.25 |
| LHX2 | 7.04E-20 | 0.41 | 0.15 | 0.001 | 2.07E-15 | 63.04 | 2.4758E-32 | 0.29 | 0.10 | 0.002 | 4.5127E-28 | 14.63 | | 3.22 |
| PTCH2 | 7.51E-102 | 3.50 | 0.85 | 0.121 | 2.21E-97 | 24.51 | 6.531E-82 | 0.88 | 0.57 | 0.187 | 1.1904E-77 | 2.68 | | 3.09 |
| CSGALNACT1 | 2.01E-32 | 1.33 | 0.33 | 0.022 | 5.89E-28 | 19.72 | 1.2098E-27 | 0.11 | 0.10 | 0.003 | 2.2051E-23 | 3.67 | | 2.86 |
| WIF1 | 1.57E-24 | 0.94 | 0.26 | 0.016 | 4.60E-20 | 15.41 | 6.5644E-78 | 0.58 | 0.30 | 0.020 | 1.1965E-73 | 8.65 | 1.31 | 2.80 |
| NRP2 | 7.27E-37 | 1.01 | 0.39 | 0.035 | 2.13E-32 | 11.25 | 5.4246E-47 | 0.23 | 0.20 | 0.014 | 9.8874E-43 | 3.24 | | 4.69 |
| FGD5 | 5.85E-29 | 0.71 | 0.33 | 0.026 | 1.72E-24 | 9.09 | 1.95E-37 | 0.25 | 0.17 | 0.014 | 3.5543E-33 | 2.94 | 2.11 | 3.13 |
| TSPAN18 | 3.71E-32 | 0.84 | 0.41 | 0.046 | 1.09E-27 | 7.42 | 5.4959E-33 | 0.22 | 0.15 | 0.013 | 1.0017E-28 | 2.51 | | 4.90 |
| TGFB2 | 7.82E-19 | 0.75 | 0.22 | 0.024 | 2.29E-14 | 6.99 | 4.0563E-66 | 0.45 | 0.22 | 0.009 | 7.3935E-62 | 11.04 | | 4.73 |
| FGF20 | 8.11E-11 | 0.38 | 0.13 | 0.012 | 2.38E-06 | 4.27 | 6.9613E-68 | 0.65 | 0.30 | 0.030 | 1.2688E-63 | 6.50 | | 2.85 |
| MYB | 2.38E-05 | 0.40 | 0.06 | 0.007 | 0.700 | 3.22 | 2.6699E-45 | 0.32 | 0.19 | 0.016 | 4.8664E-41 | 3.74 | | 5.22 |
| MICAL2 | 1.02E-14 | 0.65 | 0.36 | 0.103 | 3.00E-10 | 2.31 | 3.0415E-25 | 0.18 | 0.12 | 0.012 | 5.5437E-21 | 1.78 | | 2.44 |
| TNF | 6.48E-11 | 0.41 | 0.20 | 0.037 | 1.90E-06 | 2.26 | 1.0955E-40 | 0.33 | 0.18 | 0.016 | 1.9967E-36 | 3.71 | | 2.61 |
| SH3GL3 | 2.01E-14 | 0.66 | 0.44 | 0.149 | 5.91E-10 | 1.96 | 3.0316E-33 | 0.20 | 0.18 | 0.021 | 5.5258E-29 | 1.65 | 2.46 | 5.76 |
| EDAR | 7.39E-16 | 0.95 | 0.59 | 0.314 | 2.17E-11 | 1.78 | 3.0061E-92 | 0.78 | 0.49 | 0.098 | 5.4793E-88 | 3.90 | | 3.51 |
| DUSP6 | 1.39E-07 | 0.18 | 0.10 | 0.010 | 0.00409 | 1.76 | 4.373E-223 | 1.90 | 0.86 | 0.265 | 7.971E-219 | 6.14 | | 3.71 |
| ALCAM | 7.26E-49 | 1.57 | 0.99 | 0.889 | 2.13E-44 | 1.74 | 8.4782E-77 | 0.90 | 0.50 | 0.167 | 1.5453E-72 | 2.73 | | 4.10 |
| CCDC85A | 1.47E-06 | 0.52 | 0.17 | 0.059 | 0.0431 | 1.51 | 3.1888E-21 | 0.17 | 0.09 | 0.008 | 5.8121E-17 | 2.01 | | 5.39 |

**Supplementary Table 2: Human Specific Placode Enriched Genes. This table was generated through a two-step process. First, U score = avg_log2FC x pct.1/pct.2. U score > 1.5 for human, <0.5 for mouse was calculated using data from Glover et al. (2023) and Qu et al. (2022). Secondly, any genes found to be placode enriched in mouse by either Sennett et al. (2015) or Sulic et al. (2023) were not included in this table.**

| | Human | | | | | | Mouse (Qu et al., 2022) | | | | | |
|---|---|---|---|---|---|---|---|---|---|---|---|---|
| | p_val | avg_log2FC | pct.1 | pct.2 | p_val_adj | UScore | p_val | avg_log2FC | pct.1 | pct.2 | p_val_adj | UScore |
| FAT3 | 3.72E-49 | 2.591366 | 0.414 | 0.018 | 1.09E-44 | 59.60 | 0.48663145 | -0.0003 | 0 | 0.001 | 1 | 0.00 |
| LGR5 | 3.60E-30 | 0.874001 | 0.248 | 0.004 | 1.06E-25 | 54.19 | 0.213983084 | -0.00137 | 0 | 0.002 | 1 | 0.00 |
| RHBDL3 | 1.30E-53 | 1.637244 | 0.471 | 0.025 | 3.81E-49 | 30.85 | 0.104367095 | -0.00699 | 0.06 | 0.08 | 1 | -0.01 |
| GRIP2 | 1.92E-37 | 0.873513 | 0.338 | 0.011 | 5.64E-33 | 26.84 | 0.364577158 | -0.00204 | 0 | 0.001 | 1 | 0.00 |
| DACH1 | 1.35E-28 | 2.10928 | 0.261 | 0.022 | 3.97E-24 | 25.02 | 4.21358E-12 | 0.178778 | 0.127 | 0.052 | 7.68009E-08 | 0.44 |
| VIT | 5.32E-16 | 0.582797 | 0.140 | 0.004 | 1.56E-11 | 20.40 | 0.669847608 | -0.00055 | 0 | 0 | 1 | NE |
| KIRREL3 | 2.55E-28 | 1.839472 | 0.287 | 0.026 | 7.48E-24 | 20.30 | 0.577308617 | 0.007625 | 0.019 | 0.014 | 1 | 0.01 |
| RARB | 3.97E-35 | 1.707979 | 0.363 | 0.033 | 1.16E-30 | 18.79 | 0.000129003 | -0.03028 | 0.002 | 0.026 | 1 | 0.00 |
| KCNMA1 | 3.06E-31 | 2.56447 | 0.338 | 0.056 | 8.97E-27 | 15.48 | 0.151429417 | -0.01409 | 0.01 | 0.02 | 1 | -0.01 |
| CLDN1 | 4.91E-29 | 1.043623 | 0.331 | 0.029 | 1.44E-24 | 11.91 | 0.001750241 | -0.07082 | 0.056 | 0.106 | 1 | -0.04 |
| PPM1H | 9.55E-23 | 0.672834 | 0.242 | 0.016 | 2.80E-18 | 10.18 | 0.038323169 | -0.00316 | 0.002 | 0.003 | 1 | 0.00 |
| DPP4 | 1.48E-08 | 0.261699 | 0.076 | 0.002 | 0.000434 | 9.94 | 0.182795106 | -0.00582 | 0.004 | 0.013 | 1 | 0.00 |
| TBC1D9 | 2.14E-45 | 1.922563 | 0.573 | 0.112 | 6.29E-41 | 9.84 | 0.899749538 | 0.001105 | 0.004 | 0.003 | 1 | 0.00 |
| LDLRAD4 | 8.71E-22 | 1.314958 | 0.274 | 0.037 | 2.56E-17 | 9.74 | 0.213983084 | -6.4E-16 | 0 | 0.002 | 1 | 0.00 |
| TMEM108 | 1.52E-23 | 1.140527 | 0.306 | 0.036 | 4.47E-19 | 9.69 | 0.000255999 | -0.06152 | 0.031 | 0.079 | 1 | -0.02 |
| VWC2 | 1.07E-10 | 0.491112 | 0.096 | 0.005 | 3.15E-06 | 9.43 | 0.48663145 | 0 | 0 | 0.001 | 1 | 0.00 |
| SLC16A10 | 8.90E-40 | 1.601076 | 0.541 | 0.103 | 2.61E-35 | 8.41 | 0.054013496 | -0.01899 | 0.01 | 0.027 | 1 | -0.01 |
| CCDC68 | 3.61E-08 | 0.322953 | 0.076 | 0.003 | 0.001061 | 8.18 | 4.87835E-06 | -0.05796 | 0.027 | 0.082 | 0.088917746 | -0.02 |
| ADAM23 | 1.06E-22 | 0.95117 | 0.293 | 0.036 | 3.12E-18 | 7.74 | 0.364577158 | 0 | 0 | 0.001 | 1 | 0.00 |
| HUNK | 4.39E-49 | 1.513592 | 0.694 | 0.151 | 1.29E-44 | 6.96 | 0.003259411 | -0.08729 | 0.156 | 0.222 | 1 | -0.06 |
| UBASH3B | 1.19E-22 | 1.401086 | 0.299 | 0.062 | 3.49E-18 | 6.76 | 0.575872336 | -1.6E-05 | 0.013 | 0.008 | 1 | 0.00 |
| CSMD2 | 1.06E-29 | 1.577909 | 0.522 | 0.125 | 3.12E-25 | 6.59 | 1 | 0 | 0 | 0 | 1 | NE |
| MME | 2.55E-25 | 1.666149 | 0.350 | 0.091 | 7.50E-21 | 6.41 | 0.056143846 | -0.00441 | 0.019 | 0.039 | 1 | 0.00 |
| TSPAN7 | 1.68E-11 | 0.589849 | 0.121 | 0.012 | 4.95E-07 | 5.95 | 0.286061331 | -0.00755 | 0.071 | 0.084 | 1 | -0.01 |
| LOR | 1.78E-05 | 0.128688 | 0.045 | 0.001 | 0.521661 | 5.79 | 0.012992802 | -0.01799 | 0.056 | 0.086 | 1 | -0.01 |

| | | | | | | | | | | | | |
|---|---|---|---|---|---|---|---|---|---|---|---|---|
| TENM4 | 2.42E-25 | 0.775186 | 0.350 | 0.047 | 7.09E-21 | 5.77 | 1.18643E-06 | 0.13764 | 0.192 | 0.139 | 0.021625054 | 0.19 |
| PIP5K1B | 1.07E-12 | 0.612491 | 0.159 | 0.017 | 3.14E-08 | 5.73 | 0.023051241 | 0.001228 | 0.002 | 0.003 | 1 | 0.00 |
| KCNQ5 | 2.47E-10 | 1.245143 | 0.185 | 0.041 | 7.25E-06 | 5.62 | 0.063116982 | -0.00696 | 0 | 0.005 | 1 | 0.00 |
| PAMR1 | 8.32E-18 | 1.098783 | 0.293 | 0.058 | 2.44E-13 | 5.55 | 0.669847608 | 0 | 0 | 0 | 1 | NE |
| KCNQ3 | 3.39E-19 | 1.630901 | 0.459 | 0.139 | 9.95E-15 | 5.39 | 0.766110338 | 0.000119 | 0.002 | 0.001 | 1 | 0.00 |
| CACNA2D3 | 2.09E-19 | 2.191725 | 0.242 | 0.102 | 6.15E-15 | 5.20 | 1 | 0 | 0 | 0 | 1 | NE |
| TMEM179 | 1.50E-12 | 0.329897 | 0.134 | 0.009 | 4.41E-08 | 4.91 | 0.48663145 | -0.00068 | 0 | 0.001 | 1 | 0.00 |
| DNAJC6 | 2.98E-19 | 0.96788 | 0.363 | 0.074 | 8.75E-15 | 4.75 | 4.86701E-05 | 0.00335 | 0.004 | 0.011 | 0.887109501 | 0.00 |
| PALMD | 2.68E-14 | 0.41486 | 0.178 | 0.016 | 7.87E-10 | 4.62 | 0.713970532 | -0.00034 | 0.029 | 0.032 | 1 | 0.00 |
| PHEX | 8.26E-17 | 0.923988 | 0.261 | 0.053 | 2.43E-12 | 4.55 | 0.48663145 | -0.00123 | 0 | 0.001 | 1 | 0.00 |
| SNED1 | 4.23E-21 | 0.943589 | 0.369 | 0.078 | 1.24E-16 | 4.46 | 0.3329764 | 1.28E-15 | 0.002 | 0 | 1 | NE |
| POF1B | 4.93E-14 | 0.659673 | 0.229 | 0.035 | 1.45E-09 | 4.32 | 1.73073E-09 | 0.129793 | 0.144 | 0.054 | 3.15459E-05 | 0.35 |
| GATA3 | 1.71E-30 | 1.334126 | 0.548 | 0.177 | 5.02E-26 | 4.13 | 1.58619E-07 | 0.246581 | 0.683 | 0.695 | 0.002891145 | 0.24 |
| ENPP6 | 1.39E-08 | 0.373537 | 0.108 | 0.010 | 0.000408 | 4.03 | 1 | 0 | 0 | 0 | 1 | NE |
| PHYHIPL | 1.52E-12 | 0.335137 | 0.153 | 0.013 | 4.46E-08 | 3.94 | 0.765706343 | -0.00364 | 0.006 | 0.009 | 1 | 0.00 |
| TSPAN8 | 3.37E-06 | 0.134219 | 0.057 | 0.002 | 0.099036 | 3.83 | 0.550643616 | 0.000427 | 0.002 | 0.001 | 1 | 0.00 |
| LCP1 | 5.39E-13 | 0.256738 | 0.146 | 0.010 | 1.58E-08 | 3.75 | 0.874442962 | -3.2E-15 | 0.008 | 0.006 | 1 | 0.00 |
| P2RY14 | 4.27E-07 | 0.159148 | 0.070 | 0.003 | 0.012528 | 3.71 | 0.14631683 | 0.00159 | 0.006 | 0.002 | 1 | 0.00 |
| KAT2B | 2.11E-19 | 1.01503 | 0.427 | 0.121 | 6.19E-15 | 3.58 | 0.990889314 | -0.00289 | 0.042 | 0.043 | 1 | 0.00 |
| PAK3 | 3.35E-15 | 0.611266 | 0.261 | 0.045 | 9.85E-11 | 3.55 | 4.17072E-06 | -0.09113 | 0.058 | 0.132 | 0.076019652 | -0.04 |
| ENC1 | 4.25E-09 | 0.181392 | 0.096 | 0.005 | 0.000125 | 3.48 | 6.4244E-05 | 0.084811 | 0.102 | 0.048 | 1 | 0.18 |
| ADGRB1 | 1.82E-15 | 0.339754 | 0.210 | 0.023 | 5.36E-11 | 3.10 | 0.669847608 | -0.00054 | 0 | 0 | 1 | NE |
| ICA1 | 8.19E-26 | 1.212752 | 0.637 | 0.252 | 2.40E-21 | 3.07 | 0.6277553 | -0.00013 | 0.004 | 0.006 | 1 | 0.00 |
| GLCCI1 | 3.13E-28 | 1.409593 | 0.752 | 0.351 | 9.18E-24 | 3.02 | 0.430204236 | -0.00331 | 0.002 | 0.005 | 1 | 0.00 |
| LRP1B | 8.80E-17 | 1.78346 | 0.516 | 0.305 | 2.58E-12 | 3.02 | 0.669847608 | -0.00059 | 0 | 0 | 1 | NE |
| THRB | 5.11E-11 | 0.559118 | 0.204 | 0.039 | 1.50E-06 | 2.92 | 0.063116982 | -0.00187 | 0 | 0.005 | 1 | 0.00 |
| KCND2 | 2.67E-10 | 0.803481 | 0.236 | 0.065 | 7.85E-06 | 2.92 | 0.166093215 | -0.00231 | 0 | 0.003 | 1 | 0.00 |
| KIF26B | 9.64E-11 | 0.895762 | 0.261 | 0.083 | 2.83E-06 | 2.82 | 0.669847608 | -5.4E-13 | 0 | 0 | 1 | NE |
| TMEM132B | 6.19E-09 | 0.258618 | 0.102 | 0.010 | 0.000182 | 2.64 | 0.669847608 | -0.00028 | 0 | 0 | 1 | NE |
| KIF5C | 2.74E-10 | 0.491861 | 0.178 | 0.034 | 8.04E-06 | 2.58 | 0.680938647 | 0.001528 | 0.021 | 0.022 | 1 | 0.00 |
| CASZ1 | 1.87E-15 | 0.788873 | 0.344 | 0.106 | 5.48E-11 | 2.56 | 0.503146083 | -0.00037 | 0.017 | 0.011 | 1 | 0.00 |
| SEZ6L | 0.000148 | 0.067281 | 0.038 | 0.001 | 1 | 2.56 | 1 | 0 | 0 | 0 | 1 | NE |

| | | | | | | | | | | | | |
|---|---|---|---|---|---|---|---|---|---|---|---|---|
| VAV3 | 3.85E-17 | 1.133504 | 0.554 | 0.246 | 1.13E-12 | 2.55 | 3.96681E-08 | -0.22725 | 0.452 | 0.596 | 0.00072303 | -0.17 |
| ESRRG | 2.55E-07 | 0.398621 | 0.108 | 0.017 | 0.007492 | 2.53 | 0.669847608 | -0.00035 | 0 | 0 | 1 | NE |
| SORBS1 | 3.31E-19 | 0.934087 | 0.522 | 0.193 | 9.73E-15 | 2.53 | 9.53378E-05 | 0.016708 | 0.008 | 0.005 | 1 | 0.03 |
| KCNN1 | 8.99E-12 | 0.445256 | 0.191 | 0.034 | 2.64E-07 | 2.50 | 0.00913916 | 0.014916 | 0.017 | 0.008 | 1 | 0.03 |
| CTTNBP2 | 3.33E-13 | 0.412453 | 0.229 | 0.038 | 9.77E-09 | 2.49 | 0.804098129 | -0.00042 | 0.013 | 0.01 | 1 | 0.00 |
| BEST3 | 1.32E-05 | 0.190567 | 0.064 | 0.005 | 0.386848 | 2.44 | 1 | 0 | 0 | 0 | 1 | NE |
| SRGAP2 | 3.09E-21 | 0.783494 | 0.554 | 0.181 | 9.06E-17 | 2.40 | 0.006121101 | 0.088425 | 0.248 | 0.205 | 1 | 0.11 |
| SGPP2 | 1.88E-13 | 0.795474 | 0.382 | 0.127 | 5.53E-09 | 2.39 | 0.48663145 | -0.0007 | 0 | 0.001 | 1 | 0.00 |
| PREX2 | 1.18E-14 | 1.088772 | 0.478 | 0.219 | 3.47E-10 | 2.38 | 0.776686078 | 3.2E-16 | 0.006 | 0.004 | 1 | 0.00 |
| ARHGAP8 | 2.16E-20 | 0.873006 | 0.471 | 0.179 | 6.35E-16 | 2.30 | 0.096164022 | -0.0328 | 0.052 | 0.079 | 1 | -0.02 |
| ERC2 | 1.16E-06 | 0.619284 | 0.121 | 0.033 | 0.034184 | 2.27 | 0.374699764 | -0.00379 | 0.002 | 0.006 | 1 | 0.00 |
| CXADR | 1.03E-21 | 0.90785 | 0.624 | 0.256 | 3.02E-17 | 2.21 | 3.62933E-08 | 0.145771 | 0.383 | 0.363 | 0.000661518 | 0.15 |
| GPR161 | 4.75E-15 | 0.712997 | 0.408 | 0.132 | 1.39E-10 | 2.20 | 0.829625269 | -0.00672 | 0.031 | 0.037 | 1 | -0.01 |
| ACVR2A | 1.44E-60 | 1.671724 | 0.911 | 0.738 | 4.24E-56 | 2.06 | 8.28485E-05 | 0.114547 | 0.238 | 0.18 | 1 | 0.15 |
| ARHGAP42 | 2.12E-15 | 0.934966 | 0.503 | 0.228 | 6.21E-11 | 2.06 | 0.332500105 | -0.02564 | 0.221 | 0.248 | 1 | -0.02 |
| CYSLTR2 | 0.004452 | 0.082115 | 0.025 | 0.001 | 1 | 2.05 | 1 | 0 | 0 | 0 | 1 | NE |
| TENM3 | 1.07E-44 | 1.500704 | 0.917 | 0.672 | 3.14E-40 | 2.05 | 0.008751638 | -0.05397 | 0.171 | 0.228 | 1 | -0.04 |
| DPEP1 | 0.000146 | 0.053723 | 0.038 | 0.001 | 1 | 2.04 | 0.669847608 | -3.6E-14 | 0 | 0 | 1 | NE |
| ADAMTS6 | 2.48E-08 | 0.771803 | 0.229 | 0.088 | 0.000729 | 2.01 | 0.731916537 | 0.006839 | 0.017 | 0.013 | 1 | 0.01 |
| RALGPS1 | 4.97E-16 | 0.902174 | 0.580 | 0.261 | 1.46E-11 | 2.00 | 0.976269267 | -0.00092 | 0.019 | 0.019 | 1 | 0.00 |
| SLC8A1 | 2.01E-06 | 0.702815 | 0.210 | 0.074 | 0.058868 | 1.99 | 0.182884177 | -0.01067 | 0.015 | 0.026 | 1 | -0.01 |
| PACRG | 3.21E-18 | 1.176012 | 0.726 | 0.430 | 9.44E-14 | 1.99 | 0.023341215 | 0.011851 | 0.013 | 0.003 | 1 | 0.05 |
| PAPSS2 | 7.43E-09 | 0.335016 | 0.159 | 0.027 | 0.000218 | 1.97 | 0.703093208 | 2.03E-10 | 0.01 | 0.007 | 1 | 0.00 |
| SHANK2 | 5.37E-07 | 0.405436 | 0.146 | 0.031 | 0.015768 | 1.91 | 0.669847608 | -0.0003 | 0 | 0 | 1 | NE |
| CPA6 | 3.83E-28 | 1.577267 | 0.764 | 0.634 | 1.12E-23 | 1.90 | 0.030545964 | -0.01674 | 0.002 | 0.014 | 1 | 0.00 |
| VCAN | 7.97E-25 | 0.463385 | 0.516 | 0.126 | 2.34E-20 | 1.90 | 6.15877E-51 | -0.90935 | 0.348 | 0.675 | 1.12256E-46 | -0.47 |
| GULP1 | 5.08E-12 | 0.589369 | 0.350 | 0.112 | 1.49E-07 | 1.84 | 0.915813688 | -3.2E-15 | 0.004 | 0.006 | 1 | 0.00 |
| ALDH2 | 9.27E-11 | 0.613278 | 0.312 | 0.105 | 2.72E-06 | 1.82 | 0.745673283 | 0.022008 | 0.717 | 0.713 | 1 | 0.02 |
| ANKRD6 | 7.13E-11 | 1.002484 | 0.459 | 0.254 | 2.09E-06 | 1.81 | 0.176143491 | 0.030013 | 0.096 | 0.084 | 1 | 0.03 |
| IL22RA1 | 4.08E-10 | 0.332123 | 0.185 | 0.034 | 1.20E-05 | 1.81 | 0.669847608 | -0.0005 | 0 | 0 | 1 | NE |
| OSBPL6 | 1.47E-12 | 0.996711 | 0.510 | 0.284 | 4.32E-08 | 1.79 | 0.22679774 | -0.0084 | 0.004 | 0.011 | 1 | 0.00 |
| GLI2 | 1.05E-27 | 1.193791 | 0.796 | 0.549 | 3.07E-23 | 1.73 | 0.000428662 | -0.07746 | 0.092 | 0.158 | 1 | -0.05 |

| | | | | | | | | | | | | |
|---|---|---|---|---|---|---|---|---|---|---|---|---|
| SNTG2 | 1.16E-07 | 0.540693 | 0.172 | 0.054 | 0.003409 | 1.72 | 0.669847608 | -0.00041 | 0 | 0 | 1 | NE |
| LEPR | 8.34E-12 | 0.578048 | 0.389 | 0.135 | 2.45E-07 | 1.67 | 0.022001247 | 0.025652 | 0.021 | 0.008 | 1 | 0.07 |
| PSD3 | 2.44E-54 | 1.31858 | 0.994 | 0.789 | 7.15E-50 | 1.66 | 0.552898691 | 0.018311 | 0.092 | 0.077 | 1 | 0.02 |
| MGAT5 | 3.95E-20 | 1.085004 | 0.854 | 0.573 | 1.16E-15 | 1.62 | 4.52018E-05 | -0.13656 | 0.231 | 0.334 | 0.823893922 | -0.09 |
| IRX2 | 2.99E-15 | 0.706943 | 0.465 | 0.206 | 8.78E-11 | 1.60 | 2.75424E-08 | -0.12562 | 0.635 | 0.769 | 0.000502015 | -0.10 |
| MYRFL | 7.21E-05 | 0.13766 | 0.057 | 0.005 | 1 | 1.57 | 1 | 0 | 0 | 0 | 1 | NE |
| CMTM7 | 4.60E-12 | 0.336743 | 0.255 | 0.056 | 1.35E-07 | 1.53 | 0.293267434 | -0.02393 | 0.202 | 0.23 | 1 | -0.02 |
| SFXN1 | 7.92E-20 | 0.908176 | 0.688 | 0.408 | 2.33E-15 | 1.53 | 1.98463E-08 | -0.19349 | 0.394 | 0.544 | 0.000361738 | -0.14 |
| LAMA1 | 0.00498 | 0.304848 | 0.045 | 0.009 | 1 | 1.52 | 0.619559797 | 0.007514 | 0.025 | 0.018 | 1 | 0.01 |
| OPCML | 1.43E-05 | 0.623959 | 0.083 | 0.034 | 0.421248 | 1.52 | 0.079987398 | -0.00058 | 0 | 0.004 | 1 | 0.00 |
| GNAL | 4.50E-06 | 0.32995 | 0.096 | 0.021 | 0.132192 | 1.51 | 0.400236335 | 0.004997 | 0.013 | 0.007 | 1 | 0.01 |
| PRRG4 | 9.23E-09 | 0.44641 | 0.242 | 0.072 | 0.000271 | 1.50 | 0.092190736 | -0.01229 | 0.019 | 0.033 | 1 | -0.01 |

**Supplementary Table 3: Mouse Specific Placode Enriched Genes. U score = avg_log2FC x pct.1/pct.2. U score > 1.5 for mouse, <0.5 for human.**

| | Human | | | | | | Mouse (Qu et al., 2022) | | | | | | Sennett et al. (2015) | Sulic et al. (2023) |
|---|---|---|---|---|---|---|---|---|---|---|---|---|---|---|
| | p_val | avg_log2FC | pct.1 | pct.2 | p_val_adj | UScore | p_val | avg_log2FC | pct.1 | pct.2 | p_val_adj | UScore | log2FC | log2FC |
| Dkk4 | 0.090272 | 0.012803 | 0.006 | 0.000 | 1 | NE | 8.6E-161 | 2.640725 | 0.492 | 0.018 | 1.6E-156 | 72.18 | 2.20 | 3.17 |
| Ltb | 0.282728 | 0.043207 | 0.032 | 0.013 | 1 | 0.11 | 6.6E-139 | 1.824606 | 0.488 | 0.041 | 1.2E-134 | 21.72 | | 4.53 |
| Ascl4 | 0.044002 | 0.02631 | 0.013 | 0.001 | 1 | 0.34 | 1.81E-98 | 0.849237 | 0.344 | 0.017 | 3.29E-94 | 17.18 | 2.31 | 3.51 |
| Trps1 | 0.005159 | 0.019146 | 0.401 | 0.271 | 1 | 0.03 | 7.1E-132 | 0.971082 | 0.494 | 0.044 | 1.3E-127 | 10.90 | | 3.55 |
| Nkain3 | 0.699602 | -0.03283 | 0.038 | 0.042 | 1 | -0.03 | 1.69E-30 | 0.186021 | 0.098 | 0.002 | 3.09E-26 | 9.12 | | 4.45 |
| Madcam1 | 0.240087 | 0.009309 | 0.019 | 0.007 | 1 | 0.03 | 1.72E-66 | 0.526566 | 0.250 | 0.015 | 3.14E-62 | 8.78 | | 6.55 |
| Slc14a1 | 0.287178 | -0.00756 | 0.000 | 0.005 | 1 | 0.00 | 3.73E-34 | 0.199629 | 0.115 | 0.003 | 6.79E-30 | 7.65 | | 5.84 |
| Rragd | 0.872578 | -0.00594 | 0.038 | 0.035 | 1 | -0.01 | 1.13E-66 | 0.462154 | 0.256 | 0.017 | 2.05E-62 | 6.96 | | 3.71 |
| Il23a | 0.012121 | 0.03597 | 0.045 | 0.010 | 1 | 0.16 | 6.58E-19 | 0.110621 | 0.062 | 0.001 | 1.2E-14 | 6.86 | | 3.61 |
| Cers4 | 0.080426 | 0.192722 | 0.299 | 0.217 | 1 | 0.27 | 9.08E-86 | 0.650291 | 0.356 | 0.034 | 1.65E-81 | 6.81 | | 3.62 |
| Socs2 | 0.538771 | -0.01584 | 0.006 | 0.010 | 1 | -0.01 | 2.5E-155 | 1.310568 | 0.717 | 0.163 | 4.5E-151 | 5.76 | | 2.92 |
| Frem1 | 5.62E-05 | 0.066384 | 0.312 | 0.192 | 1 | 0.11 | 4.36E-90 | 0.579889 | 0.383 | 0.040 | 7.94E-86 | 5.55 | | 4.17 |
| Etv4 | 0.752597 | 0.016976 | 0.019 | 0.017 | 1 | 0.02 | 1.46E-95 | 0.696021 | 0.458 | 0.058 | 2.67E-91 | 5.50 | | 3.07 |
| Cxcr4 | 0.99019 | 0.000751 | 0.006 | 0.007 | 1 | 0.00 | 7.89E-23 | 0.123508 | 0.079 | 0.002 | 1.44E-18 | 4.88 | | 3.48 |
| Ccl20 | 0.010371 | 0.016382 | 0.013 | 0.000 | 1 | NE | 2.79E-20 | 0.320868 | 0.088 | 0.006 | 5.09E-16 | 4.71 | | 2.69 |
| Krt17 | 0.002416 | 0.385511 | 0.739 | 0.609 | 1 | 0.47 | 1.8E-192 | 1.935453 | 0.896 | 0.386 | 3.3E-188 | 4.49 | | 2.29 |
| S100a9 | 0.557621 | -0.00796 | 0.000 | 0.002 | 1 | 0.00 | 2.55E-41 | 0.481566 | 0.183 | 0.021 | 4.65E-37 | 4.20 | | 1.75 |
| Ifitm1 | 1 | 0 | 0.000 | 0.000 | 1 | NE | 1.1E-122 | 1.23485 | 0.662 | 0.212 | 1.9E-118 | 3.86 | | 2.93 |
| Stxbp6 | 0.000115 | -0.36889 | 0.459 | 0.638 | 1 | -0.27 | 1E-114 | 0.944208 | 0.585 | 0.144 | 1.9E-110 | 3.84 | | 2.56 |
| Rnf182 | 0.005504 | 0.066946 | 0.051 | 0.010 | 1 | 0.34 | 1.98E-39 | 0.29183 | 0.169 | 0.013 | 3.6E-35 | 3.79 | | 3.29 |
| Cd74 | 0.070993 | 0.065815 | 0.045 | 0.014 | 1 | 0.21 | 1.35E-86 | 0.884106 | 0.510 | 0.131 | 2.46E-82 | 3.44 | | 3.42 |
| Ctgf | 0.996746 | 0.003405 | 0.013 | 0.013 | 1 | 0.00 | 1.86E-51 | 0.573076 | 0.312 | 0.056 | 3.39E-47 | 3.19 | | 2.16 |
| Gpm6b | 0.00488 | -0.14153 | 0.108 | 0.122 | 1 | -0.13 | 2.56E-29 | 0.142725 | 0.108 | 0.005 | 4.66E-25 | 3.08 | | 2.73 |
| Tspan5 | 0.165012 | 0.117997 | 0.102 | 0.060 | 1 | 0.20 | 5.89E-98 | 0.875122 | 0.540 | 0.162 | 1.07E-93 | 2.92 | | 1.46 |
| Bmp2 | 0.077075 | 0.126681 | 0.102 | 0.053 | 1 | 0.24 | 3.36E-71 | 0.938331 | 0.450 | 0.147 | 6.12E-67 | 2.87 | 1.36 | 1.71 |
| Trib1 | 0.003884 | -0.25592 | 0.127 | 0.210 | 1 | -0.15 | 1.23E-54 | 0.55348 | 0.338 | 0.068 | 2.25E-50 | 2.75 | | 2.08 |

| | | | | | | | | | | | | | | |
|---|---|---|---|---|---|---|---|---|---|---|---|---|---|---|
| Steap4 | 0.443217 | 0.016948 | 0.013 | 0.004 | 1 | 0.06 | 1.48E-20 | 0.171424 | 0.088 | 0.006 | 2.71E-16 | 2.51 | | 4.97 |
| Kcnmb4 | 0.001984 | 0.184467 | 0.210 | 0.106 | 1 | 0.37 | 6.2E-45 | 0.345569 | 0.235 | 0.033 | 1.13E-40 | 2.46 | | 2.82 |
| Cadm1 | 0.39756 | 0.207144 | 0.809 | 0.836 | 1 | 0.20 | 7.4E-127 | 1.161116 | 0.746 | 0.376 | 1.3E-122 | 2.30 | | 1.68 |
| Ncam1 | 0.27858 | -0.00516 | 0.025 | 0.051 | 1 | 0.00 | 5.24E-46 | 0.439003 | 0.304 | 0.058 | 9.55E-42 | 2.30 | | 3.32 |
| Tubb2b | 0.108193 | -0.02099 | 0.000 | 0.011 | 1 | 0.00 | 1.03E-78 | 0.541884 | 0.462 | 0.112 | 1.88E-74 | 2.24 | | 2.20 |
| Trib2 | 0.00221 | 0.005821 | 0.038 | 0.008 | 1 | 0.03 | 5.87E-34 | 0.141825 | 0.140 | 0.009 | 1.07E-29 | 2.21 | | 1.51 |
| Stx11 | 0.039568 | -0.05873 | 0.006 | 0.033 | 1 | -0.01 | 6.82E-30 | 0.257584 | 0.150 | 0.018 | 1.24E-25 | 2.15 | | 1.60 |
| Mybpc1 | 0.557621 | -0.004 | 0.000 | 0.002 | 1 | 0.00 | 2.2E-48 | 0.475269 | 0.310 | 0.070 | 4.01E-44 | 2.10 | | 1.85 |
| Cacna1e | 0.008031 | 0.057645 | 0.038 | 0.005 | 1 | 0.44 | 1.19E-09 | 0.067886 | 0.031 | 0.001 | 2.17E-05 | 2.10 | | 4.95 |
| Fn1 | 0.064524 | -0.02452 | 0.178 | 0.132 | 1 | -0.03 | 6.54E-80 | 0.347093 | 0.448 | 0.074 | 1.19E-75 | 2.10 | | 3.50 |
| Eif4e3 | 0.070437 | 0.058563 | 0.108 | 0.059 | 1 | 0.11 | 2.86E-37 | 0.340477 | 0.221 | 0.036 | 5.21E-33 | 2.09 | | 3.07 |
| Cxcl1 | 0.23434 | -0.01262 | 0.000 | 0.007 | 1 | 0.00 | 4.38E-32 | 0.349037 | 0.206 | 0.035 | 7.99E-28 | 2.05 | | 3.34 |
| Ptchd4 | 6.42E-31 | -1.27211 | 0.452 | 0.866 | 1.88E-26 | -0.66 | 1.94E-13 | 0.085592 | 0.046 | 0.002 | 3.53E-09 | 1.97 | | 5.20 |
| Ifitm3 | 0.258893 | 0.006244 | 0.038 | 0.023 | 1 | 0.01 | 1.6E-268 | 1.735494 | 1.000 | 0.897 | 2.9E-264 | 1.93 | 0.62 | 1.50 |
| Etv5 | 0.847426 | 0.080888 | 0.338 | 0.320 | 1 | 0.09 | 1.15E-51 | 0.499364 | 0.404 | 0.105 | 2.1E-47 | 1.92 | | 2.00 |
| Zdhhc2 | 0.451814 | 0.108229 | 0.223 | 0.181 | 1 | 0.13 | 4.51E-47 | 0.503228 | 0.358 | 0.098 | 8.22E-43 | 1.84 | | 1.50 |
| Scube1 | 0.343786 | 0.033596 | 0.006 | 0.004 | 1 | 0.05 | 2.14E-26 | 0.204778 | 0.133 | 0.015 | 3.9E-22 | 1.82 | | 2.78 |
| Pdlim4 | 0.311577 | -0.09677 | 0.140 | 0.170 | 1 | -0.08 | 4.69E-61 | 0.640115 | 0.448 | 0.159 | 8.56E-57 | 1.80 | | 1.53 |
| Pfn2 | 0.290083 | 0.022555 | 0.108 | 0.079 | 1 | 0.03 | 6.04E-90 | 0.999038 | 0.650 | 0.367 | 1.1E-85 | 1.77 | | 1.22 |
| Cyba | 0.592266 | -0.04755 | 0.051 | 0.059 | 1 | -0.04 | 1.1E-104 | 0.917158 | 0.740 | 0.387 | 2E-100 | 1.75 | | 1.52 |
| Snai3 | 0.401075 | 0.023947 | 0.038 | 0.021 | 1 | 0.04 | 5.08E-16 | 0.118902 | 0.058 | 0.004 | 9.27E-12 | 1.72 | | 3.78 |
| Tll1 | 3.56E-06 | -0.43305 | 0.274 | 0.481 | 0.104506 | -0.25 | 4.89E-11 | 0.04386 | 0.038 | 0.001 | 8.91E-07 | 1.67 | | 4.73 |
| Stap1 | 0.090272 | 0.005627 | 0.006 | 0.000 | 1 | NE | 4.62E-09 | 0.054508 | 0.029 | 0.001 | 8.41E-05 | 1.58 | | 7.89 |
| Sox2 | 0.644171 | 0.019667 | 0.006 | 0.003 | 1 | 0.04 | 1.49E-14 | 0.059782 | 0.052 | 0.002 | 2.71E-10 | 1.55 | | 3.62 |
| Cxcl10 | 1 | 0 | 0.000 | 0.000 | 1 | NE | 4.07E-11 | 0.123401 | 0.050 | 0.004 | 7.41E-07 | 1.54 | | 1.87 |
| Ctxn1 | 0.245799 | 0.009447 | 0.013 | 0.002 | 1 | 0.06 | 9.55E-54 | 0.55667 | 0.481 | 0.174 | 1.74E-49 | 1.54 | | 1.63 |
| Map1b | 0.031663 | -0.15114 | 0.197 | 0.177 | 1 | -0.17 | 1.23E-64 | 0.660864 | 0.569 | 0.249 | 2.23E-60 | 1.51 | | 1.03 |

**Supplementary Table 4: Shared Dermal Condensate Enriched Genes. U score = avg_log2FC x pct.1/pct.2. U score > 1.5 for mouse and human.**

| | Human | | | | | | Mouse (Qu et al., 2022) | | | | | | Sennett et al. (2015) |
|---|---|---|---|---|---|---|---|---|---|---|---|---|---|
| | p_val | avg_log2FC | pct.1 | pct.2 | p_val_adj | UScore | p_val | avg_log2FC | pct.1 | pct.2 | p_val_adj | UScore | log2FC |
| SOX2 | 1.02E-29 | 0.320004 | 0.104 | 0.005 | 3E-25 | 6.66 | 2.3E-149 | 1.451343 | 0.439 | 0.001 | 4.1E-145 | 637.14 | 9.64 |
| BMP3 | 1.69E-12 | 0.180838 | 0.049 | 0.004 | 4.95E-08 | 2.22 | 3.53E-76 | 1.114155 | 0.245 | 0.002 | 6.43E-72 | 136.48 | 5.40 |
| SOX18 | 2.3E-45 | 0.443514 | 0.18 | 0.013 | 6.74E-41 | 6.14 | 6.45E-98 | 0.874396 | 0.316 | 0.003 | 1.18E-93 | 92.10 | 8.25 |
| ADAMTS18 | 2.78E-50 | 1.240805 | 0.488 | 0.235 | 8.15E-46 | 2.58 | 1.41E-62 | 0.436917 | 0.203 | 0.001 | 2.58E-58 | 88.69 | 8.23 |
| FOXD1 | 6.12E-40 | 0.54671 | 0.204 | 0.027 | 1.8E-35 | 4.13 | 1.4E-161 | 1.691708 | 0.516 | 0.01 | 2.6E-157 | 87.29 | 7.81 |
| DLL1 | 1.84E-66 | 0.712489 | 0.288 | 0.028 | 5.41E-62 | 7.33 | 1.94E-60 | 0.516979 | 0.209 | 0.003 | 3.53E-56 | 36.02 | 7.65 |
| SOX11 | 1.5E-126 | 1.19309 | 0.634 | 0.126 | 4.3E-122 | 6.00 | 2E-240 | 2.392083 | 0.824 | 0.131 | 3.6E-236 | 15.05 | 3.59 |
| GLIS1 | 1E-107 | 1.330173 | 0.856 | 0.669 | 3E-103 | 1.70 | 3.27E-92 | 0.930726 | 0.391 | 0.026 | 5.96E-88 | 14.00 | |
| IGF1 | 3.65E-45 | 0.626442 | 0.155 | 0.006 | 1.07E-40 | 16.18 | 3.2E-176 | 2.39949 | 0.693 | 0.124 | 5.8E-172 | 13.41 | 2.88 |
| PLK2 | 4.1E-99 | 0.204855 | 0.397 | 0.037 | 1.22E-94 | 2.20 | 1.2E-113 | 1.150402 | 0.499 | 0.044 | 2.3E-109 | 13.05 | 4.12 |
| NRP2 | 1.3E-161 | 1.620842 | 0.645 | 0.1 | 3.9E-157 | 10.45 | 1.1E-139 | 2.282766 | 0.543 | 0.097 | 2E-135 | 12.78 | 4.12 |
| PDE3A | 4.8E-153 | 1.744093 | 0.927 | 0.635 | 1.4E-148 | 2.55 | 3.25E-93 | 1.105969 | 0.448 | 0.048 | 5.93E-89 | 10.32 | 4.08 |
| PRDM1 | 2.22E-72 | 1.095171 | 0.479 | 0.13 | 6.52E-68 | 4.04 | 1.26E-93 | 1.157208 | 0.46 | 0.053 | 2.3E-89 | 10.04 | 4.96 |
| FGF10 | 2.37E-26 | 0.575365 | 0.235 | 0.063 | 6.95E-22 | 2.15 | 3.4E-130 | 1.481206 | 0.6 | 0.093 | 6.3E-126 | 9.56 | 3.83 |
| SMOC1 | 6.8E-179 | 1.842147 | 0.831 | 0.198 | 2E-174 | 7.73 | 3.65E-87 | 1.119707 | 0.439 | 0.053 | 6.65E-83 | 9.27 | 5.24 |
| CXCR4 | 2.46E-41 | 0.412053 | 0.155 | 0.009 | 7.23E-37 | 7.10 | 7.41E-31 | 0.332555 | 0.125 | 0.005 | 1.35E-26 | 8.31 | 5.32 |
| KIF26B | 0 | 3.129237 | 0.969 | 0.354 | 0 | 8.57 | 1.46E-24 | 0.17492 | 0.093 | 0.002 | 2.67E-20 | 8.13 | |
| HEYL | 9.92E-42 | 0.181995 | 0.146 | 0.006 | 2.91E-37 | 4.43 | 1.79E-32 | 0.310289 | 0.137 | 0.006 | 3.27E-28 | 7.08 | |
| EFCC1 | 1.11E-96 | 1.093614 | 0.332 | 0.017 | 3.25E-92 | 21.36 | 2.78E-14 | 0.137663 | 0.051 | 0.001 | 5.06E-10 | 7.02 | 6.42 |
| BMP7 | 4.5E-251 | 2.352424 | 0.761 | 0.077 | 1.3E-246 | 23.25 | 1.1E-97 | 1.056613 | 0.573 | 0.102 | 2.01E-93 | 5.94 | 4.93 |
| NDNF | 2.16E-35 | 0.887174 | 0.341 | 0.118 | 6.35E-31 | 2.56 | 2.32E-78 | 1.263277 | 0.394 | 0.086 | 4.22E-74 | 5.79 | 4.75 |
| EBF1 | 0 | 2.633083 | 0.996 | 0.845 | 0 | 3.10 | 4.8E-132 | 2.506295 | 0.627 | 0.297 | 8.8E-128 | 5.29 | 2.50 |
| PTCH2 | 2.5E-147 | 1.503054 | 0.9 | 0.362 | 7.3E-143 | 3.74 | 2.6E-62 | 0.482917 | 0.343 | 0.033 | 4.73E-58 | 5.02 | |
| HHIP | 2.2E-155 | 2.013644 | 0.561 | 0.075 | 6.5E-151 | 15.06 | 2.93E-47 | 0.76886 | 0.26 | 0.04 | 5.34E-43 | 5.00 | 3.42 |
| NPTX2 | 5.86E-48 | 0.563054 | 0.171 | 0.01 | 1.72E-43 | 9.63 | 9.08E-51 | 0.75412 | 0.334 | 0.051 | 1.66E-46 | 4.94 | 3.53 |
| TAGLN3 | 6.4E-14 | 0.094927 | 0.04 | 0.001 | 1.88E-09 | 3.80 | 6.07E-15 | 0.096522 | 0.051 | 0.001 | 1.11E-10 | 4.92 | |
| COL23A1 | 2.57E-96 | 1.397689 | 0.86 | 0.55 | 7.53E-92 | 2.19 | 1.5E-120 | 1.447795 | 0.633 | 0.196 | 2.7E-116 | 4.68 | 3.36 |
| HS3ST3B1 | 3.55E-57 | 0.562922 | 0.341 | 0.062 | 1.04E-52 | 3.10 | 1.24E-42 | 0.48958 | 0.239 | 0.026 | 2.26E-38 | 4.50 | 3.92 |

| | | | | | | | | | | | | | |
|---|---|---|---|---|---|---|---|---|---|---|---|---|---|
| CPNE5 | 3.49E-84 | 0.831935 | 0.348 | 0.035 | 1.02E-79 | 8.27 | 3.9E-176 | 1.678114 | 0.857 | 0.364 | 7.1E-172 | 3.95 | 3.72 |
| CADPS2 | 4.94E-323 | 2.888342 | 0.792 | 0.038 | 1.45E-318 | 60.20 | 1.84E-14 | 0.125585 | 0.057 | 0.002 | 3.35E-10 | 3.58 | 4.00 |
| SOX4 | 7.33E-59 | 0.958994 | 0.761 | 0.44 | 2.15E-54 | 1.66 | 4.9E-193 | 1.744069 | 0.901 | 0.488 | 9E-189 | 3.22 | |
| SPON1 | 1.6E-284 | 2.906145 | 0.936 | 0.336 | 4.8E-280 | 8.10 | 5.06E-63 | 0.871753 | 0.445 | 0.122 | 9.23E-59 | 3.18 | |
| LAMC3 | 0 | 3.210807 | 0.923 | 0.157 | 0 | 18.88 | 1.44E-16 | 0.202969 | 0.093 | 0.008 | 2.62E-12 | 2.36 | 4.52 |
| LAMA2 | 6.1E-197 | 2.552021 | 0.811 | 0.253 | 1.8E-192 | 8.18 | 3.46E-59 | 0.865818 | 0.406 | 0.158 | 6.31E-55 | 2.22 | 2.64 |
| TP53INP1 | 2.5E-53 | 0.729789 | 0.33 | 0.081 | 7.35E-49 | 2.97 | 3.19E-17 | 0.146625 | 0.09 | 0.006 | 5.81E-13 | 2.20 | 2.73 |
| NTF3 | 3.57E-40 | 0.266419 | 0.24 | 0.041 | 1.05E-35 | 1.56 | 6.02E-34 | 0.637869 | 0.313 | 0.105 | 1.1E-29 | 1.90 | 3.51 |
| SEMA6A | 4.17E-91 | 1.171855 | 0.791 | 0.414 | 1.22E-86 | 2.24 | 2.43E-57 | 0.929345 | 0.576 | 0.294 | 4.42E-53 | 1.82 | 2.18 |

**Supplementary Table 5: Human Specific Dermal Condensate Enriched Genes. This table was generated by a two-step process. First U score = avg_log2FC x pct.1/pct.2. U score > 1.5 for human, <0.5 for mouse using data from Glover et al. (2023) and Qu et al. (2022). Secondly, any genes found to be dermal condensate enriched in mouse by Sennett et al. (2015) were not included in this table.**

| | Human | | | | | | Mouse (Qu et al., 2022) | | | | | |
|---|---|---|---|---|---|---|---|---|---|---|---|---|
| | p_val | avg_log2FC | pct.1 | pct.2 | p_val_adj | UScore | p_val | avg_log2FC | pct.1 | pct.2 | p_val_adj | UScore |
| RAP1GAP2 | 1.13E-277 | 2.809703 | 0.668 | 0.021 | 3.31E-273 | 89.38 | 0.818519696 | -0.00186 | 0.009 | 0.010 | 1 | 0.00 |
| SUSD4 | 2.73E-93 | 0.58353 | 0.251 | 0.003 | 8.01E-89 | 48.82 | 0.964424531 | -5.7E-09 | 0.006 | 0.007 | 1 | 0.00 |
| CRYM | 9.51E-97 | 1.229708 | 0.291 | 0.010 | 2.79E-92 | 35.78 | 0.006194139 | 0.018733 | 0.012 | 0.001 | 1 | 0.22 |
| SCUBE1 | 1.56E-119 | 1.284403 | 0.372 | 0.014 | 4.58E-115 | 34.13 | 0.062992208 | 0.018671 | 0.006 | 0.002 | 1 | 0.06 |
| ANO2 | 9.00E-171 | 1.619874 | 0.519 | 0.027 | 2.64E-166 | 31.14 | 0.000614336 | 0.020917 | 0.009 | 0.000 | 1 | NE |
| TMEM132E | 1.85E-33 | 0.326131 | 0.093 | 0.001 | 5.44E-29 | 30.33 | 0.023988914 | -0.01827 | 0.000 | 0.009 | 1 | 0.00 |
| KL | 1.30E-34 | 0.297941 | 0.097 | 0.001 | 3.81E-30 | 28.90 | 0.363615574 | 0.005249 | 0.006 | 0.002 | 1 | 0.02 |
| LAMA3 | 2.79E-239 | 1.388358 | 0.692 | 0.037 | 8.18E-235 | 25.97 | 0.366981978 | 0 | 0.006 | 0.002 | 1 | 0.00 |
| ANKRD29 | 4.25E-122 | 1.167022 | 0.383 | 0.019 | 1.25E-117 | 23.52 | 0.000155967 | 0.090895 | 0.054 | 0.021 | 1 | 0.23 |
| CFAP299 | 1.37E-95 | 2.204927 | 0.350 | 0.033 | 4.02E-91 | 23.39 | 0.070755385 | 0.010937 | 0.003 | 0.000 | 1 | NE |
| ATP8A2 | 6.05E-78 | 1.187708 | 0.253 | 0.013 | 1.78E-73 | 23.11 | 0.366981416 | -0.00274 | 0.000 | 0.002 | 1 | 0.00 |
| ALPL | 2.53E-122 | 1.321368 | 0.395 | 0.023 | 7.42E-118 | 22.69 | 0.899830119 | -0.00508 | 0.009 | 0.011 | 1 | 0.00 |
| KCNN2 | 1.05E-274 | 3.646063 | 0.789 | 0.143 | 3.08E-270 | 20.12 | 0.001979855 | -0.09124 | 0.063 | 0.114 | 1 | -0.05 |
| ITGA8 | 3.28E-147 | 1.897399 | 0.559 | 0.065 | 9.64E-143 | 16.32 | 0.002491668 | -0.08782 | 0.182 | 0.251 | 1 | -0.06 |
| BRINP2 | 8.06E-35 | 0.539747 | 0.113 | 0.004 | 2.37E-30 | 15.25 | 0.771710263 | -0.00038 | 0.000 | 0.000 | 1 | NE |
| HEY2 | 4.71E-130 | 1.367419 | 0.474 | 0.043 | 1.38E-125 | 15.07 | 0.005939773 | 0.07106 | 0.078 | 0.066 | 1 | 0.08 |
| CACNA1D | 2.24E-194 | 1.749479 | 0.667 | 0.080 | 6.57E-190 | 14.59 | 0.426191964 | -0.00987 | 0.027 | 0.035 | 1 | -0.01 |
| LINGO2 | 1.57E-56 | 0.739472 | 0.197 | 0.010 | 4.60E-52 | 14.57 | 0.060596835 | -0.02557 | 0.012 | 0.029 | 1 | -0.01 |
| ASTN1 | 2.05E-93 | 1.245411 | 0.359 | 0.031 | 6.01E-89 | 14.42 | 0.976013014 | -0.0013 | 0.003 | 0.003 | 1 | 0.00 |
| MEGF11 | 1.18E-87 | 0.928802 | 0.321 | 0.021 | 3.46E-83 | 14.20 | 0.771710263 | -0.00082 | 0.000 | 0.000 | 1 | NE |
| DCX | 1.05E-19 | 0.227939 | 0.058 | 0.001 | 3.07E-15 | 13.22 | 0.771710263 | -2.9E-12 | 0.000 | 0.000 | 1 | NE |
| PGM5 | 2.39E-78 | 0.569585 | 0.268 | 0.013 | 7.02E-74 | 11.74 | 0.00358881 | -0.11322 | 0.084 | 0.146 | 1 | -0.07 |
| NRXN3 | 1.35E-128 | 1.360597 | 0.525 | 0.064 | 3.96E-124 | 11.16 | 0.006168392 | 0.008048 | 0.003 | 0.002 | 1 | 0.01 |
| CTNND2 | 7.02E-138 | 1.811042 | 0.621 | 0.102 | 2.06E-133 | 11.03 | 0.004494784 | 0.014866 | 0.009 | 0.000 | 1 | NE |
| NOTUM | 6.03E-41 | 0.759151 | 0.162 | 0.013 | 1.77E-36 | 9.46 | 0.229238899 | 0.074877 | 0.170 | 0.142 | 1 | 0.09 |

| | | | | | | | | | | | | |
|---|---|---|---|---|---|---|---|---|---|---|---|---|
| ELMOD1 | 2.11E-73 | 0.978055 | 0.324 | 0.034 | 6.21E-69 | 9.32 | 1 | 0 | 0.000 | 0.000 | 1 | NE |
| KCNH8 | 6.63E-78 | 0.494511 | 0.277 | 0.015 | 1.95E-73 | 9.13 | 1 | 0 | 0.000 | 0.000 | 1 | NE |
| HES5 | 1.49E-25 | 0.224053 | 0.078 | 0.002 | 4.36E-21 | 8.74 | 8.09836E-12 | 0.233769 | 0.039 | 0.000 | 1.48E-07 | NE |
| PCSK2 | 5.30E-91 | 1.583741 | 0.443 | 0.083 | 1.55E-86 | 8.45 | 0.789922331 | -0.00123 | 0.003 | 0.003 | 1 | 0.00 |
| PTPRR | 7.71E-31 | 0.651232 | 0.102 | 0.008 | 2.26E-26 | 8.30 | 1 | 0 | 0.000 | 0.000 | 1 | NE |
| PREX2 | 1.79E-193 | 2.149445 | 0.821 | 0.219 | 5.26E-189 | 8.06 | 0.007984755 | 9.01E-05 | 0.233 | 0.269 | 1 | 0.00 |
| LIX1 | 7.71E-59 | 0.791122 | 0.253 | 0.026 | 2.26E-54 | 7.70 | 8.5149E-10 | -0.41087 | 0.534 | 0.703 | 1.55E-05 | -0.31 |
| IGSF3 | 6.67E-127 | 1.311259 | 0.546 | 0.098 | 1.96E-122 | 7.31 | 0.006427353 | 0.006648 | 0.373 | 0.419 | 1 | 0.01 |
| CYTL1 | 4.25E-53 | 0.367064 | 0.191 | 0.010 | 1.25E-48 | 7.01 | 0.771710263 | -0.00162 | 0.000 | 0.000 | 1 | NE |
| IGSF5 | 9.89E-48 | 0.668212 | 0.206 | 0.020 | 2.90E-43 | 6.88 | 1 | 0 | 0.000 | 0.000 | 1 | NE |
| TPD52 | 3.74E-100 | 0.522144 | 0.381 | 0.029 | 1.10E-95 | 6.86 | 0.016637664 | -0.00394 | 0.048 | 0.090 | 1 | 0.00 |
| TENM4 | 3.09E-228 | 2.38238 | 0.871 | 0.303 | 9.07E-224 | 6.85 | 6.27021E-13 | -0.22494 | 0.030 | 0.155 | 1.14E-08 | -0.04 |
| PPEF1 | 4.69E-50 | 1.077814 | 0.251 | 0.042 | 1.38E-45 | 6.44 | 1 | 0 | 0.000 | 0.000 | 1 | NE |
| TSPAN18 | 3.33E-131 | 1.61101 | 0.681 | 0.184 | 9.76E-127 | 5.96 | 0.000106049 | 0.04079 | 0.021 | 0.006 | 1 | 0.14 |
| CAPN6 | 1.55E-24 | 0.209908 | 0.084 | 0.003 | 4.54E-20 | 5.88 | 0.000648877 | -0.09052 | 0.024 | 0.073 | 1 | -0.03 |
| PDZRN4 | 2.14E-20 | 0.452338 | 0.075 | 0.006 | 6.29E-16 | 5.65 | 0.312054936 | -0.00393 | 0.000 | 0.002 | 1 | 0.00 |
| FRAS1 | 2.76E-96 | 0.810922 | 0.446 | 0.064 | 8.12E-92 | 5.65 | 0.897216002 | -0.0015 | 0.021 | 0.025 | 1 | 0.00 |
| NTM | 1.11E-116 | 0.866587 | 0.561 | 0.088 | 3.26E-112 | 5.52 | 0.003157388 | -0.05146 | 0.039 | 0.075 | 1 | -0.03 |
| FGF13 | 2.01E-60 | 0.546904 | 0.259 | 0.026 | 5.90E-56 | 5.45 | 9.24894E-09 | -0.17434 | 0.134 | 0.260 | 0.000169 | -0.09 |
| JAG1 | 3.21E-140 | 0.905491 | 0.643 | 0.107 | 9.43E-136 | 5.44 | 0.529414484 | -0.0021 | 0.009 | 0.008 | 1 | 0.00 |
| PLCB4 | 9.43E-143 | 0.797765 | 0.630 | 0.094 | 2.77E-138 | 5.35 | 0.464998614 | 0.009368 | 0.012 | 0.006 | 1 | 0.02 |
| ATP8A1 | 1.69E-84 | 1.434752 | 0.485 | 0.135 | 4.97E-80 | 5.15 | 0.03400147 | 0.015235 | 0.018 | 0.004 | 1 | 0.07 |
| ADGRB1 | 7.44E-89 | 1.121406 | 0.536 | 0.117 | 2.18E-84 | 5.14 | 1.56873E-05 | 0.054022 | 0.018 | 0.000 | 0.285933 | NE |
| SEMA3G | 2.86E-28 | 0.258901 | 0.098 | 0.005 | 8.39E-24 | 5.07 | 0.06612548 | 0.001301 | 0.006 | 0.002 | 1 | 0.00 |
| MFAP5 | 4.56E-26 | 0.382326 | 0.106 | 0.008 | 1.34E-21 | 5.07 | 0.027453526 | -0.01996 | 0.000 | 0.009 | 1 | 0.00 |
| EGFLAM | 2.76E-115 | 1.316338 | 0.672 | 0.176 | 8.09E-111 | 5.03 | 0.000307796 | -0.03626 | 0.119 | 0.170 | 1 | -0.03 |
| GPC5 | 6.38E-42 | 1.638976 | 0.275 | 0.091 | 1.87E-37 | 4.95 | 0.026341495 | 0.004443 | 0.006 | 0.000 | 1 | NE |
| KIRREL3 | 7.26E-72 | 0.90925 | 0.375 | 0.069 | 2.13E-67 | 4.94 | 0.026341495 | 0.006503 | 0.006 | 0.000 | 1 | NE |
| SLC27A6 | 2.26E-62 | 1.105772 | 0.381 | 0.086 | 6.64E-58 | 4.90 | 0.004845053 | -0.06112 | 0.015 | 0.050 | 1 | -0.02 |
| GREB1 | 1.63E-110 | 1.531255 | 0.579 | 0.189 | 4.78E-106 | 4.69 | 6.08694E-08 | 0.019502 | 0.036 | 0.002 | 0.001109 | 0.35 |
| GFRA2 | 1.59E-38 | 0.729496 | 0.191 | 0.030 | 4.67E-34 | 4.64 | 5.7132E-06 | -0.11465 | 0.024 | 0.090 | 0.104135 | -0.03 |
| PTPRE | 1.24E-43 | 0.561511 | 0.171 | 0.021 | 3.64E-39 | 4.57 | 0.096255241 | 1.25E-09 | 0.012 | 0.003 | 1 | 0.00 |

| | | | | | | | | | | | | |
|---|---|---|---|---|---|---|---|---|---|---|---|---|
| CSMD1 | 7.78E-192 | 2.155743 | 0.949 | 0.472 | 2.28E-187 | 4.33 | 0.266550775 | -0.0067 | 0.000 | 0.003 | 1 | 0.00 |
| CASZ1 | 7.98E-67 | 0.732666 | 0.384 | 0.068 | 2.34E-62 | 4.14 | 5.21028E-11 | -0.36383 | 0.149 | 0.322 | 9.5E-07 | -0.17 |
| TRIM36 | 2.65E-39 | 0.356219 | 0.168 | 0.015 | 7.79E-35 | 3.99 | 0.001583919 | 0.019701 | 0.015 | 0.010 | 1 | 0.03 |
| SLC44A5 | 1.57E-40 | 0.737013 | 0.259 | 0.048 | 4.60E-36 | 3.98 | 0.625473057 | -0.00212 | 0.000 | 0.001 | 1 | 0.00 |
| SLC24A3 | 9.01E-243 | 2.301829 | 0.920 | 0.551 | 2.64E-238 | 3.84 | 0.43415252 | -0.0019 | 0.000 | 0.002 | 1 | 0.00 |
| CDKL5 | 9.96E-144 | 1.839921 | 0.750 | 0.363 | 2.92E-139 | 3.80 | 0.02050582 | 0.057792 | 0.054 | 0.027 | 1 | 0.12 |
| GRID1 | 4.22E-76 | 1.503732 | 0.608 | 0.246 | 1.24E-71 | 3.72 | 0.741473263 | -0.00521 | 0.003 | 0.006 | 1 | 0.00 |
| B3GAT1 | 1.41E-11 | 0.102579 | 0.036 | 0.001 | 4.15E-07 | 3.69 | 1 | 0 | 0.000 | 0.000 | 1 | NE |
| OCA2 | 1.15E-70 | 0.914732 | 0.488 | 0.121 | 3.37E-66 | 3.69 | 0.771710263 | -0.00045 | 0.000 | 0.000 | 1 | NE |
| BCAR3 | 1.46E-104 | 1.164227 | 0.699 | 0.225 | 4.30E-100 | 3.62 | 0.45001006 | 0.0027 | 0.006 | 0.002 | 1 | 0.01 |
| LDLRAD4 | 1.41E-66 | 0.781143 | 0.444 | 0.100 | 4.13E-62 | 3.47 | 0.006442555 | 0.102553 | 0.107 | 0.069 | 1 | 0.16 |
| MYOCD | 4.90E-26 | 0.604651 | 0.160 | 0.029 | 1.44E-21 | 3.34 | 0.43415252 | -0.00246 | 0.000 | 0.002 | 1 | 0.00 |
| CPE | 1.04E-47 | 0.889877 | 0.368 | 0.101 | 3.06E-43 | 3.24 | 0.0170982 | 0.057174 | 0.045 | 0.029 | 1 | 0.09 |
| GULP1 | 6.25E-115 | 1.365953 | 0.816 | 0.353 | 1.83E-110 | 3.16 | 2.25279E-05 | -0.08251 | 0.125 | 0.202 | 0.410616 | -0.05 |
| RBP4 | 1.85E-10 | 0.095444 | 0.033 | 0.001 | 5.42E-06 | 3.15 | 3.44874E-08 | -0.09596 | 0.000 | 0.049 | 0.000629 | 0.00 |
| ARL9 | 2.11E-28 | 0.515115 | 0.158 | 0.027 | 6.19E-24 | 3.01 | 0.771710263 | -0.00045 | 0.000 | 0.000 | 1 | NE |
| OTUD7A | 6.84E-43 | 1.159348 | 0.399 | 0.156 | 2.01E-38 | 2.97 | 0.806165084 | -0.01006 | 0.009 | 0.013 | 1 | -0.01 |
| UPP2 | 1.46E-29 | 0.647249 | 0.189 | 0.042 | 4.30E-25 | 2.91 | 0.038062691 | 0.043114 | 0.060 | 0.053 | 1 | 0.05 |
| MRAP2 | 4.58E-38 | 0.767953 | 0.297 | 0.079 | 1.34E-33 | 2.89 | 1 | 0 | 0.000 | 0.000 | 1 | NE |
| VSTM4 | 2.41E-102 | 1.402813 | 0.743 | 0.372 | 7.08E-98 | 2.80 | 4.45872E-17 | -0.34166 | 0.239 | 0.474 | 8.13E-13 | -0.17 |
| PKP4 | 4.03E-93 | 1.095411 | 0.778 | 0.308 | 1.18E-88 | 2.77 | 0.404568989 | -0.02935 | 0.063 | 0.078 | 1 | -0.02 |
| NKAIN4 | 1.63E-37 | 0.541267 | 0.228 | 0.045 | 4.78E-33 | 2.74 | 0.081306126 | 0.060668 | 0.069 | 0.043 | 1 | 0.10 |
| ME1 | 8.84E-45 | 0.268853 | 0.202 | 0.020 | 2.60E-40 | 2.72 | 0.002251808 | -0.085 | 0.048 | 0.100 | 1 | -0.04 |
| GASK1A | 2.71E-38 | 0.821723 | 0.311 | 0.096 | 7.96E-34 | 2.66 | 0.625473057 | -0.00103 | 0.000 | 0.001 | 1 | 0.00 |
| AGAP1 | 0 | 2.428347 | 1.000 | 0.913 | 0 | 2.66 | 2.34123E-07 | 0.242063 | 0.364 | 0.301 | 0.004267 | 0.29 |
| EDNRA | 2.49E-259 | 2.060883 | 0.989 | 0.787 | 7.31E-255 | 2.59 | 0.024068092 | 0.097214 | 0.373 | 0.360 | 1 | 0.10 |
| CADPS | 2.20E-33 | 0.711785 | 0.257 | 0.071 | 6.47E-29 | 2.58 | 0.126430983 | -0.00785 | 0.000 | 0.005 | 1 | 0.00 |
| PCYT1B | 1.90E-22 | 0.309345 | 0.107 | 0.013 | 5.58E-18 | 2.55 | 4.20614E-05 | 0.047661 | 0.027 | 0.003 | 0.766652 | 0.43 |
| KCNB2 | 4.64E-18 | 0.266228 | 0.086 | 0.009 | 1.36E-13 | 2.54 | 0.078969229 | 0.025757 | 0.027 | 0.021 | 1 | 0.03 |
| CNKSR2 | 7.68E-50 | 1.006409 | 0.457 | 0.181 | 2.26E-45 | 2.54 | 0.957752273 | -0.0058 | 0.024 | 0.026 | 1 | -0.01 |
| CSMD2 | 1.02E-96 | 1.433766 | 0.750 | 0.427 | 3.00E-92 | 2.52 | 0.194816356 | 0.006695 | 0.003 | 0.000 | 1 | NE |
| MCC | 1.04E-142 | 1.374282 | 0.969 | 0.532 | 3.05E-138 | 2.50 | 0.146560123 | -0.01854 | 0.134 | 0.158 | 1 | -0.02 |

| | | | | | | | | | | | | |
|---|---|---|---|---|---|---|---|---|---|---|---|---|
| ENTPD2 | 5.17E-19 | 0.113503 | 0.066 | 0.003 | 1.52E-14 | 2.50 | 0.094995075 | -0.01054 | 0.000 | 0.005 | 1 | 0.00 |
| FAM189A1 | 1.15E-94 | 1.497131 | 0.791 | 0.477 | 3.39E-90 | 2.48 | 0.000318669 | -0.04221 | 0.000 | 0.022 | 1 | 0.00 |
| RARB | 2.80E-53 | 1.375282 | 0.588 | 0.327 | 8.22E-49 | 2.47 | 0.014840042 | -0.0629 | 0.027 | 0.060 | 1 | -0.03 |
| MBP | 2.39E-32 | 0.297653 | 0.157 | 0.019 | 7.03E-28 | 2.46 | 0.575966755 | -0.00565 | 0.009 | 0.012 | 1 | 0.00 |
| DPF3 | 6.09E-31 | 0.421777 | 0.186 | 0.032 | 1.79E-26 | 2.45 | 0.563997399 | 0.007472 | 0.006 | 0.002 | 1 | 0.02 |
| RGS20 | 1.70E-46 | 0.645614 | 0.355 | 0.095 | 5.00E-42 | 2.41 | 0.625473057 | -0.00109 | 0.000 | 0.001 | 1 | 0.00 |
| IGDCC3 | 2.23E-27 | 0.503069 | 0.180 | 0.038 | 6.55E-23 | 2.38 | 0.694176596 | -0.00738 | 0.003 | 0.006 | 1 | 0.00 |
| PKNOX2 | 2.53E-35 | 0.214516 | 0.155 | 0.014 | 7.43E-31 | 2.37 | 0.771710263 | -0.00063 | 0.000 | 0.000 | 1 | NE |
| PAPPA2 | 2.74E-187 | 1.862584 | 0.962 | 0.764 | 8.05E-183 | 2.35 | 2.17237E-05 | -0.08724 | 0.346 | 0.453 | 0.395958 | -0.07 |
| ZDHHC14 | 4.78E-44 | 0.890955 | 0.497 | 0.193 | 1.40E-39 | 2.29 | 0.020840447 | -0.07734 | 0.042 | 0.082 | 1 | -0.04 |
| MMP11 | 1.90E-25 | 0.39571 | 0.155 | 0.027 | 5.58E-21 | 2.27 | 0.013249662 | -0.02662 | 0.260 | 0.310 | 1 | -0.02 |
| FRMD3 | 5.56E-60 | 1.071753 | 0.665 | 0.314 | 1.63E-55 | 2.27 | 0.264274008 | 0.00507 | 0.006 | 0.001 | 1 | 0.03 |
| TBX3 | 4.70E-29 | 0.23479 | 0.135 | 0.014 | 1.38E-24 | 2.26 | 1.21627E-07 | 0.164366 | 0.066 | 0.022 | 0.002217 | 0.49 |
| GRK5 | 2.78E-50 | 0.948871 | 0.390 | 0.165 | 8.17E-46 | 2.24 | 0.01951964 | -0.03635 | 0.033 | 0.062 | 1 | -0.02 |
| HS6ST1 | 2.06E-91 | 1.186731 | 0.689 | 0.365 | 6.04E-87 | 2.24 | 0.015245921 | 0.066831 | 0.096 | 0.084 | 1 | 0.08 |
| PLCL2 | 3.63E-56 | 1.108495 | 0.497 | 0.250 | 1.07E-51 | 2.20 | 0.114331077 | -0.00465 | 0.015 | 0.023 | 1 | 0.00 |
| SPATA13 | 8.03E-38 | 0.464906 | 0.250 | 0.053 | 2.36E-33 | 2.19 | 0.58109231 | -0.0008 | 0.006 | 0.004 | 1 | 0.00 |
| FAM53B | 1.50E-55 | 0.790293 | 0.548 | 0.198 | 4.39E-51 | 2.19 | 0.013164698 | 0.066279 | 0.104 | 0.060 | 1 | 0.11 |
| HIST1H2AC | 2.16E-28 | 0.577302 | 0.233 | 0.063 | 6.33E-24 | 2.14 | 0.228493632 | -0.00476 | 0.000 | 0.003 | 1 | 0.00 |
| LMO4 | 2.09E-74 | 1.079595 | 0.709 | 0.360 | 6.14E-70 | 2.13 | 6.1474E-09 | -0.30477 | 0.266 | 0.434 | 0.000112 | -0.19 |
| RAP1GAP | 1.48E-23 | 0.211368 | 0.106 | 0.011 | 4.34E-19 | 2.04 | 0.524506088 | -0.0117 | 0.006 | 0.010 | 1 | -0.01 |
| KCNK4 | 2.80E-09 | 0.168047 | 0.036 | 0.003 | 8.22E-05 | 2.02 | 1 | 0 | 0.000 | 0.000 | 1 | NE |
| SYNPO2 | 3.75E-66 | 0.285398 | 0.335 | 0.048 | 1.10E-61 | 1.99 | 0.082489948 | -0.0107 | 0.000 | 0.006 | 1 | 0.00 |
| CTNNA3 | 3.46E-65 | 0.306765 | 0.355 | 0.056 | 1.02E-60 | 1.94 | 0.229869237 | 0.010261 | 0.006 | 0.001 | 1 | 0.06 |
| PSTPIP2 | 1.32E-56 | 1.03598 | 0.528 | 0.292 | 3.86E-52 | 1.87 | 0.43415252 | -0.00199 | 0.000 | 0.002 | 1 | 0.00 |
| PDK3 | 5.16E-39 | 0.928928 | 0.384 | 0.193 | 1.51E-34 | 1.85 | 0.008248243 | 0.041452 | 0.284 | 0.309 | 1 | 0.04 |
| PHKB | 2.25E-95 | 1.202597 | 0.769 | 0.508 | 6.62E-91 | 1.82 | 0.093203335 | 0.0223 | 0.101 | 0.102 | 1 | 0.02 |
| REEP3 | 7.94E-45 | 0.675097 | 0.488 | 0.181 | 2.33E-40 | 1.82 | 0.003092397 | 0.11468 | 0.418 | 0.401 | 1 | 0.12 |
| NOTCH3 | 8.96E-58 | 0.525059 | 0.485 | 0.142 | 2.63E-53 | 1.79 | 0.001612116 | 0.098868 | 0.107 | 0.060 | 1 | 0.18 |
| STK33 | 2.86E-20 | 0.620251 | 0.182 | 0.063 | 8.41E-16 | 1.79 | 0.059005003 | 0.016171 | 0.006 | 0.001 | 1 | 0.10 |
| AGRN | 6.12E-48 | 0.178773 | 0.217 | 0.022 | 1.80E-43 | 1.76 | 0.809886318 | -0.0147 | 0.128 | 0.139 | 1 | -0.01 |
| ARHGAP44 | 6.48E-53 | 0.761602 | 0.636 | 0.276 | 1.90E-48 | 1.75 | 0.620539261 | 0.00688 | 0.015 | 0.015 | 1 | 0.01 |

| | | | | | | | | | | | | |
|---|---|---|---|---|---|---|---|---|---|---|---|---|
| FGD3 | 1.94E-27 | 0.662047 | 0.319 | 0.121 | 5.69E-23 | 1.75 | 8.75448E-10 | -0.00085 | 0.012 | 0.000 | 1.6E-05 | NE |
| SORCS1 | 6.71E-23 | 0.756534 | 0.346 | 0.150 | 1.97E-18 | 1.75 | 0.771710263 | -0.00042 | 0.000 | 0.000 | 1 | NE |
| CDK19 | 3.69E-138 | 1.372108 | 0.960 | 0.762 | 1.08E-133 | 1.73 | 0.001408535 | 0.108223 | 0.125 | 0.074 | 1 | 0.18 |
| DACH2 | 3.58E-13 | 0.657966 | 0.115 | 0.044 | 1.05E-08 | 1.72 | 0.572892989 | -0.01272 | 0.009 | 0.015 | 1 | -0.01 |
| JAKMIP2 | 4.59E-43 | 0.154598 | 0.188 | 0.017 | 1.35E-38 | 1.71 | 0.011535113 | 0.014409 | 0.006 | 0.002 | 1 | 0.04 |
| RPS6KA2 | 1.66E-45 | 0.711797 | 0.548 | 0.230 | 4.86E-41 | 1.70 | 0.669156519 | -0.00228 | 0.009 | 0.012 | 1 | 0.00 |
| TMEM171 | 8.78E-33 | 0.671874 | 0.388 | 0.154 | 2.58E-28 | 1.69 | 0.194816356 | 0.001761 | 0.003 | 0.000 | 1 | NE |
| ADAMTS6 | 6.03E-40 | 1.102678 | 0.477 | 0.316 | 1.77E-35 | 1.66 | 0.527545138 | 0.012506 | 0.012 | 0.006 | 1 | 0.03 |
| NCAM1 | 1.21E-68 | 1.172112 | 0.776 | 0.553 | 3.56E-64 | 1.64 | 2.31486E-10 | -0.33 | 0.334 | 0.526 | 4.22E-06 | -0.21 |
| PMEPA1 | 6.18E-37 | 0.587929 | 0.424 | 0.155 | 1.81E-32 | 1.61 | 0.001429706 | 0.048615 | 0.179 | 0.197 | 1 | 0.04 |
| MTURN | 2.43E-35 | 0.58374 | 0.333 | 0.121 | 7.15E-31 | 1.61 | 0.00047677 | 0.085172 | 0.063 | 0.040 | 1 | 0.13 |
| LUZP2 | 1.75E-15 | 0.659685 | 0.128 | 0.053 | 5.13E-11 | 1.59 | 0.266550775 | -6.1E-09 | 0.000 | 0.003 | 1 | 0.00 |
| LRFN2 | 2.24E-07 | 0.122288 | 0.026 | 0.002 | 0.006582 | 1.59 | 1 | 0 | 0.000 | 0.000 | 1 | NE |
| GPR173 | 1.13E-49 | 0.842996 | 0.546 | 0.290 | 3.33E-45 | 1.59 | 0.346990166 | 0.017881 | 0.042 | 0.039 | 1 | 0.02 |
| RNF24 | 6.53E-90 | 1.165229 | 0.863 | 0.637 | 1.92E-85 | 1.58 | 1.42527E-05 | 0.168781 | 0.149 | 0.102 | 0.259783 | 0.25 |
| COL5A3 | 8.53E-26 | 0.3853 | 0.175 | 0.043 | 2.51E-21 | 1.57 | 0.494627442 | 0.009895 | 0.009 | 0.004 | 1 | 0.02 |
| DMGDH | 9.92E-22 | 0.495455 | 0.208 | 0.066 | 2.91E-17 | 1.56 | 0.771710263 | -0.0006 | 0.000 | 0.000 | 1 | NE |
| FCHSD2 | 8.88E-54 | 0.979433 | 0.791 | 0.497 | 2.61E-49 | 1.56 | 0.037137382 | 0.032788 | 0.131 | 0.139 | 1 | 0.03 |
| CDH8 | 1.78E-14 | 0.677326 | 0.191 | 0.083 | 5.23E-10 | 1.56 | 0.061034917 | -0.02184 | 0.003 | 0.015 | 1 | 0.00 |
| KLRG2 | 2.15E-10 | 0.046999 | 0.033 | 0.001 | 6.30E-06 | 1.55 | 0.183144742 | -0.00162 | 0.003 | 0.003 | 1 | 0.00 |
| CDH11 | 1.43E-179 | 1.455461 | 0.989 | 0.937 | 4.21E-175 | 1.54 | 3.58543E-16 | 0.461621 | 0.830 | 0.772 | 6.54E-12 | 0.50 |
| NEO1 | 1.17E-58 | 0.827027 | 0.732 | 0.395 | 3.43E-54 | 1.53 | 0.002849553 | 0.089354 | 0.206 | 0.192 | 1 | 0.10 |
| PTN | 3.02E-31 | 0.12493 | 0.133 | 0.011 | 8.88E-27 | 1.51 | 6.36347E-48 | -1.14397 | 0.161 | 0.562 | 1.16E-43 | -0.33 |
| MSANTD4 | 1.04E-26 | 0.475643 | 0.206 | 0.065 | 3.06E-22 | 1.51 | 0.000790946 | 0.10663 | 0.203 | 0.178 | 1 | 0.12 |
| RAP1GAP2 | 1.13E-277 | 2.809703 | 0.668 | 0.021 | 3.31E-273 | 89.38 | 0.818519696 | -0.00186 | 0.009 | 0.010 | 1 | 0.00 |

**Supplementary Table 6: Mouse Specific Dermal Condensate Enriched Genes. U score = avg_log2FC x pct.1/pct.2. U score > 1.5 for mouse, <0.5 for human.**

| | Human | | | | | | Mouse (Qu et al., 2022) | | | | | | Sennett et al. (2015) |
|---|---|---|---|---|---|---|---|---|---|---|---|---|---|
| | p_val | avg_log2FC | pct.1 | pct.2 | p_val_adj | UScore | p_val | avg_log2FC | pct.1 | pct.2 | p_val_adj | UScore | log2FC |
| Grin2a | 0.005347 | 0.068643 | 0.018 | 0.004 | 1 | 0.31 | 3.29E-78 | 0.887006 | 0.242 | 0.001 | 6E-74 | 214.66 | 9.75 |
| Gal | 0.136196 | 1.6E-15 | 0.002 | 0.000 | 1 | NE | 1.23E-84 | 1.57334 | 0.278 | 0.003 | 2.24E-80 | 145.80 | 9.52 |
| Vat1l | 3.14E-10 | -0.20421 | 0.007 | 0.068 | 9.21E-06 | -0.02 | 9.79E-86 | 0.47617 | 0.272 | 0.002 | 1.79E-81 | 64.76 | |
| Pdlim3 | 0.745741 | 0.00644 | 0.026 | 0.020 | 1 | 0.01 | 1.93E-39 | 0.525368 | 0.134 | 0.002 | 3.52E-35 | 35.20 | 7.37 |
| Rgs16 | 0.568393 | -0.00715 | 0.004 | 0.006 | 1 | 0.00 | 5.09E-94 | 0.862124 | 0.349 | 0.010 | 9.27E-90 | 30.09 | |
| Sgip1 | 3.71E-06 | 0.121492 | 0.086 | 0.033 | 0.108785 | 0.32 | 2.6E-145 | 1.561533 | 0.588 | 0.049 | 4.8E-141 | 18.74 | 5.73 |
| Sox9 | 0.609752 | 0.002838 | 0.002 | 0.003 | 1 | 0.00 | 2.44E-69 | 0.856768 | 0.290 | 0.015 | 4.44E-65 | 16.56 | 3.89 |
| Hapln3 | 3.79E-07 | -0.14132 | 0.027 | 0.090 | 0.011142 | -0.04 | 1.7E-104 | 1.229724 | 0.457 | 0.034 | 3E-100 | 16.53 | |
| Cdkn1a | 0.876035 | 0.000442 | 0.005 | 0.007 | 1 | 0.00 | 3.9E-197 | 2.474326 | 0.776 | 0.127 | 7.2E-193 | 15.12 | 5.32 |
| Mycbpap | 0.658147 | 0.005184 | 0.013 | 0.014 | 1 | 0.00 | 5.75E-22 | 0.189184 | 0.072 | 0.001 | 1.05E-17 | 13.62 | 5.77 |
| Ceacam1 | 0.100375 | 0.076512 | 0.071 | 0.061 | 1 | 0.09 | 1.92E-27 | 0.244328 | 0.104 | 0.002 | 3.5E-23 | 12.71 | 5.77 |
| Cyp26b1 | 3.39E-22 | -0.37556 | 0.026 | 0.169 | 9.94E-18 | -0.06 | 4.3E-144 | 1.724228 | 0.612 | 0.087 | 7.8E-140 | 12.13 | 3.99 |
| Sorcs3 | 7.28E-08 | 0.310255 | 0.148 | 0.139 | 0.002136 | 0.33 | 6.04E-33 | 0.290417 | 0.125 | 0.003 | 1.1E-28 | 12.10 | 5.23 |
| Apod | 0.605794 | -0.00349 | 0.002 | 0.005 | 1 | 0.00 | 2.32E-63 | 1.16949 | 0.301 | 0.030 | 4.24E-59 | 11.73 | 2.94 |
| Igfbp3 | 0.00039 | -0.1361 | 0.270 | 0.356 | 1 | -0.10 | 7.53E-88 | 1.356801 | 0.412 | 0.048 | 1.37E-83 | 11.65 | 3.97 |
| Avpr1a | 0.193967 | 0.006401 | 0.007 | 0.002 | 1 | 0.02 | 3.18E-65 | 0.665404 | 0.290 | 0.017 | 5.8E-61 | 11.35 | 4.42 |
| Ndp | 3.93E-14 | 0.086568 | 0.035 | 0.000 | 1.15E-09 | NE | 5.94E-22 | 0.237521 | 0.084 | 0.002 | 1.08E-17 | 9.98 | 7.28 |
| Igfbp4 | 4.42E-09 | 0.058036 | 0.073 | 0.017 | 0.00013 | 0.25 | 9.5E-179 | 2.338188 | 0.728 | 0.210 | 1.7E-174 | 8.11 | 2.25 |
| Col19a1 | 0.354539 | 0 | 0.022 | 0.013 | 1 | 0.00 | 1.12E-25 | 0.165361 | 0.096 | 0.002 | 2.04E-21 | 7.94 | 5.93 |
| Klf4 | 0.0053 | -0.04747 | 0.133 | 0.125 | 1 | -0.05 | 3.5E-118 | 1.591731 | 0.582 | 0.120 | 6.3E-114 | 7.72 | 2.70 |
| Syne1 | 3.23E-19 | -0.58307 | 0.561 | 0.737 | 9.48E-15 | -0.44 | 4.04E-48 | 0.47739 | 0.233 | 0.015 | 7.36E-44 | 7.42 | 3.64 |
| Tmod2 | 0.000912 | 0.1365 | 0.080 | 0.045 | 1 | 0.24 | 6.65E-59 | 0.798381 | 0.301 | 0.034 | 1.21E-54 | 7.07 | |
| Edn3 | 0.336735 | 0.039159 | 0.040 | 0.030 | 1 | 0.05 | 1.3E-125 | 1.57263 | 0.621 | 0.150 | 2.4E-121 | 6.51 | 4.23 |
| Frzb | 0.058367 | 0.000286 | 0.011 | 0.005 | 1 | 0.00 | 2.38E-38 | 0.520399 | 0.191 | 0.016 | 4.35E-34 | 6.21 | 5.95 |
| Plppr4 | 0.483693 | 0.002117 | 0.002 | 0.003 | 1 | 0.00 | 2.38E-17 | 0.179737 | 0.069 | 0.002 | 4.33E-13 | 6.20 | |
| Lynx1 | 0.709863 | -0.00255 | 0.002 | 0.002 | 1 | 0.00 | 8.11E-39 | 0.589205 | 0.203 | 0.021 | 1.48E-34 | 5.70 | 5.69 |
| Scnn1b | 0.688659 | -0.00211 | 0.004 | 0.003 | 1 | 0.00 | 2.39E-51 | 0.679618 | 0.293 | 0.036 | 4.36E-47 | 5.53 | 4.87 |

| | | | | | | | | | | | | | |
|---|---|---|---|---|---|---|---|---|---|---|---|---|---|
| Fam150b | 0.396671 | -0.00628 | 0.002 | 0.006 | 1 | 0.00 | 3.49E-17 | 0.260478 | 0.063 | 0.003 | 6.37E-13 | 5.47 | 6.13 |
| Ccdc109b | 1.48E-06 | -0.26398 | 0.175 | 0.279 | 0.043467 | -0.17 | 3.37E-41 | 0.527653 | 0.224 | 0.022 | 6.15E-37 | 5.37 | 4.43 |
| Dio2 | 0.917951 | -0.0125 | 0.007 | 0.009 | 1 | -0.01 | 5.26E-14 | 0.195293 | 0.054 | 0.002 | 9.59E-10 | 5.27 | 7.01 |
| Lepr | 0.017375 | -0.06984 | 0.164 | 0.204 | 1 | -0.06 | 8.66E-52 | 0.833596 | 0.257 | 0.042 | 1.58E-47 | 5.10 | 3.77 |
| Lrrtm2 | 0.833729 | -0.00084 | 0.049 | 0.046 | 1 | 0.00 | 2.86E-25 | 0.231426 | 0.104 | 0.005 | 5.22E-21 | 4.81 | |
| Dclk1 | 7.23E-46 | -0.60116 | 0.880 | 0.978 | 2.12E-41 | -0.54 | 5.5E-191 | 1.948605 | 0.872 | 0.355 | 1E-186 | 4.79 | 3.73 |
| Hck | 0.010105 | 0.007498 | 0.015 | 0.003 | 1 | 0.04 | 2.61E-56 | 0.754257 | 0.361 | 0.058 | 4.77E-52 | 4.69 | 4.00 |
| Rasd1 | 0.908078 | 0.003272 | 0.007 | 0.006 | 1 | 0.00 | 5.55E-28 | 0.440396 | 0.158 | 0.015 | 1.01E-23 | 4.64 | |
| Ntng1 | 5.38E-16 | -0.77918 | 0.514 | 0.662 | 1.58E-11 | -0.60 | 7.33E-42 | 0.628757 | 0.236 | 0.032 | 1.34E-37 | 4.64 | 2.95 |
| Epn3 | 0.630919 | -0.00033 | 0.000 | 0.001 | 1 | 0.00 | 1.22E-20 | 0.19359 | 0.093 | 0.004 | 2.23E-16 | 4.50 | 6.62 |
| Nmnat2 | 0.136595 | -0.02599 | 0.005 | 0.015 | 1 | -0.01 | 1.19E-11 | 0.095621 | 0.045 | 0.001 | 2.17E-07 | 4.30 | 5.44 |
| Hs3st6 | 2.7E-06 | 0.204453 | 0.100 | 0.052 | 0.079133 | 0.39 | 2.6E-66 | 1.061339 | 0.445 | 0.111 | 4.74E-62 | 4.25 | 4.55 |
| Itih5 | 0.675694 | -0.00163 | 0.011 | 0.009 | 1 | 0.00 | 2.4E-84 | 1.695683 | 0.543 | 0.217 | 4.37E-80 | 4.24 | 2.90 |
| Galnt9 | 3.95E-05 | 0.148038 | 0.082 | 0.034 | 1 | 0.36 | 5.36E-10 | 0.102831 | 0.039 | 0.001 | 9.78E-06 | 4.01 | |
| Serinc2 | 0.250027 | 0.008032 | 0.053 | 0.060 | 1 | 0.01 | 2.54E-30 | 0.267349 | 0.149 | 0.010 | 4.64E-26 | 3.98 | 5.18 |
| Necab1 | 0.016797 | -0.09839 | 0.058 | 0.095 | 1 | -0.06 | 5.09E-19 | 0.35339 | 0.090 | 0.008 | 9.28E-15 | 3.98 | 3.36 |
| Grid2 | 0.005607 | -0.00542 | 0.077 | 0.045 | 1 | -0.01 | 2.34E-14 | 0.139831 | 0.054 | 0.002 | 4.27E-10 | 3.78 | |
| Flt1 | 0.010028 | 0.007254 | 0.009 | 0.022 | 1 | 0.00 | 5.72E-19 | 0.215414 | 0.087 | 0.005 | 1.04E-14 | 3.75 | |
| B4galnt1 | 0.625622 | 0.002768 | 0.004 | 0.002 | 1 | 0.01 | 3.77E-60 | 0.814635 | 0.454 | 0.099 | 6.88E-56 | 3.74 | 3.34 |
| Stc2 | 0.630919 | -9.9E-15 | 0.000 | 0.001 | 1 | 0.00 | 9.18E-21 | 0.219447 | 0.099 | 0.006 | 1.67E-16 | 3.62 | |
| Filip1l | 4.84E-07 | -0.2371 | 0.541 | 0.667 | 0.014219 | -0.19 | 6.95E-53 | 0.950842 | 0.358 | 0.095 | 1.27E-48 | 3.58 | |
| Arhgap15 | 0.00066 | -0.04258 | 0.137 | 0.115 | 1 | -0.05 | 2.14E-50 | 0.855011 | 0.397 | 0.096 | 3.91E-46 | 3.54 | 3.74 |
| Nrtn | 0.014352 | 0.066469 | 0.029 | 0.016 | 1 | 0.12 | 8.6E-128 | 1.528704 | 0.794 | 0.346 | 1.6E-123 | 3.51 | |
| Qpct | 0.134639 | -0.0224 | 0.007 | 0.018 | 1 | -0.01 | 4.07E-39 | 0.689967 | 0.236 | 0.050 | 7.42E-35 | 3.26 | |
| Pxdc1 | 4.79E-15 | -0.28324 | 0.069 | 0.207 | 1.41E-10 | -0.09 | 1.12E-92 | 1.327145 | 0.594 | 0.243 | 2.04E-88 | 3.24 | 2.87 |
| Aplp1 | 0.061745 | 0.028481 | 0.013 | 0.004 | 1 | 0.09 | 3.63E-47 | 0.689418 | 0.322 | 0.069 | 6.62E-43 | 3.22 | 2.82 |
| Dnajc6 | 0.000758 | -0.03499 | 0.020 | 0.033 | 1 | -0.02 | 1.46E-11 | 0.081658 | 0.039 | 0.001 | 2.66E-07 | 3.18 | 6.05 |
| Aqp1 | 0.059464 | -0.06437 | 0.040 | 0.065 | 1 | -0.04 | 4.19E-25 | 0.433843 | 0.167 | 0.023 | 7.64E-21 | 3.15 | |
| Trim9 | 0.527008 | 0.012628 | 0.011 | 0.009 | 1 | 0.02 | 2.88E-29 | 0.417742 | 0.173 | 0.023 | 5.25E-25 | 3.14 | 5.42 |
| Whrn | 0.406772 | -0.04181 | 0.122 | 0.142 | 1 | -0.04 | 3.47E-17 | 0.123698 | 0.075 | 0.003 | 6.32E-13 | 3.09 | 4.94 |
| 1810041L15Rik | 0.647003 | -0.00036 | 0.002 | 0.001 | 1 | 0.00 | 4.36E-21 | 0.154069 | 0.096 | 0.005 | 7.95E-17 | 2.96 | 4.37 |
| Smad7 | 0.35574 | 0.025609 | 0.062 | 0.047 | 1 | 0.03 | 1.75E-67 | 0.893947 | 0.475 | 0.144 | 3.19E-63 | 2.95 | |

| | | | | | | | | | | | | | |
|---|---|---|---|---|---|---|---|---|---|---|---|---|---|
| Gpm6b | 0.000208 | -0.19887 | 0.315 | 0.324 | 1 | -0.19 | 3.09E-32 | 0.469674 | 0.212 | 0.034 | 5.64E-28 | 2.93 | |
| Cd24a | 8.3E-11 | 0.033169 | 0.064 | 0.010 | 2.44E-06 | 0.21 | 3.4E-218 | 1.852985 | 0.988 | 0.634 | 6.1E-214 | 2.89 | 2.91 |
| Mtus1 | 0.003488 | -0.02183 | 0.029 | 0.029 | 1 | -0.02 | 1.28E-45 | 0.735841 | 0.400 | 0.105 | 2.33E-41 | 2.80 | |
| Sdc1 | 1.21E-10 | 0.070216 | 0.160 | 0.070 | 3.54E-06 | 0.16 | 2.8E-134 | 1.444066 | 0.788 | 0.407 | 5E-130 | 2.80 | 2.58 |
| Ryr2 | 0.010983 | -0.02899 | 0.031 | 0.035 | 1 | -0.03 | 2.46E-14 | 0.132935 | 0.063 | 0.003 | 4.49E-10 | 2.79 | 4.38 |
| Pde4d | 2.79E-20 | 0.250543 | 0.944 | 0.801 | 8.18E-16 | 0.30 | 2.79E-82 | 1.070242 | 0.594 | 0.233 | 5.09E-78 | 2.73 | |
| Lrrn3 | 0.516478 | -0.00364 | 0.098 | 0.090 | 1 | 0.00 | 3.76E-31 | 0.482423 | 0.203 | 0.036 | 6.85E-27 | 2.72 | 2.60 |
| Ism1 | 5.72E-09 | -0.30763 | 0.080 | 0.182 | 0.000168 | -0.14 | 3.8E-44 | 0.761202 | 0.415 | 0.121 | 6.92E-40 | 2.61 | 3.10 |
| Prokr2 | 1 | 0 | 0.000 | 0.000 | 1 | NE | 1.99E-07 | 0.082473 | 0.030 | 0.001 | 0.003633 | 2.47 | 4.62 |
| Soga3 | 3.41E-08 | 0.221307 | 0.089 | 0.050 | 0.001002 | 0.39 | 1.86E-26 | 0.486813 | 0.203 | 0.040 | 3.39E-22 | 2.47 | 4.03 |
| Capg | 0.108225 | 0.020575 | 0.024 | 0.011 | 1 | 0.04 | 4.83E-55 | 0.664032 | 0.493 | 0.133 | 8.81E-51 | 2.46 | |
| Cbr3 | 0.090131 | -0.01117 | 0.007 | 0.017 | 1 | 0.00 | 2.16E-62 | 0.991214 | 0.633 | 0.255 | 3.94E-58 | 2.46 | |
| Lxn | 4.17E-05 | -0.07002 | 0.053 | 0.102 | 1 | -0.04 | 3.1E-119 | 1.400518 | 0.740 | 0.423 | 5.7E-115 | 2.45 | 2.74 |
| Lef1 | 2.37E-12 | 0.243637 | 0.883 | 0.746 | 6.95E-08 | 0.29 | 2.03E-89 | 1.093115 | 0.824 | 0.396 | 3.71E-85 | 2.27 | |
| 6030419C18Rik | 0.684497 | 0.00979 | 0.009 | 0.006 | 1 | 0.01 | 3.33E-26 | 0.390529 | 0.185 | 0.032 | 6.08E-22 | 2.26 | |
| Wisp1 | 1.96E-06 | -0.0936 | 0.007 | 0.047 | 0.057603 | -0.01 | 7.31E-84 | 1.208371 | 0.791 | 0.439 | 1.33E-79 | 2.18 | 2.03 |
| Prkar1b | 0.374088 | 0.049262 | 0.158 | 0.137 | 1 | 0.06 | 5.33E-31 | 0.458692 | 0.224 | 0.048 | 9.71E-27 | 2.14 | 3.14 |
| Fam181b | 0.100705 | 0.006133 | 0.004 | 0.001 | 1 | 0.02 | 2.18E-39 | 0.704964 | 0.334 | 0.110 | 3.97E-35 | 2.14 | 3.00 |
| Tspan7 | 0.703624 | -0.00011 | 0.005 | 0.003 | 1 | 0.00 | 1.92E-26 | 0.455416 | 0.206 | 0.044 | 3.49E-22 | 2.13 | |
| Sphkap | 1.6E-138 | -2.42746 | 0.046 | 0.586 | 4.7E-134 | -0.19 | 1.16E-35 | 0.740929 | 0.343 | 0.120 | 2.12E-31 | 2.12 | |
| Peli1 | 2E-20 | -0.5685 | 0.472 | 0.624 | 5.88E-16 | -0.43 | 1.5E-111 | 1.21623 | 0.857 | 0.506 | 2.7E-107 | 2.06 | |
| Spon2 | 2.59E-10 | -0.25359 | 0.060 | 0.161 | 7.59E-06 | -0.09 | 4.9E-16 | 0.243951 | 0.099 | 0.012 | 8.93E-12 | 2.01 | 2.62 |
| Zmiz1 | 0.487225 | -0.05586 | 0.222 | 0.247 | 1 | -0.05 | 3.27E-52 | 0.746232 | 0.504 | 0.187 | 5.97E-48 | 2.01 | |
| Vldlr | 1.85E-09 | 0.045956 | 0.055 | 0.009 | 5.43E-05 | 0.28 | 2.38E-28 | 0.518188 | 0.218 | 0.057 | 4.33E-24 | 1.98 | 1.63 |
| Cacna1g | 2.64E-11 | 0.295439 | 0.231 | 0.180 | 7.76E-07 | 0.38 | 6.16E-49 | 0.756831 | 0.436 | 0.168 | 1.12E-44 | 1.96 | 2.25 |
| Abhd2 | 0.001909 | -0.1514 | 0.169 | 0.229 | 1 | -0.11 | 3.32E-21 | 0.783255 | 0.334 | 0.135 | 6.04E-17 | 1.94 | 2.68 |
| Rtn1 | 0.895501 | 0.000322 | 0.007 | 0.006 | 1 | 0.00 | 9.43E-26 | 0.44405 | 0.152 | 0.035 | 1.72E-21 | 1.93 | 2.01 |
| Epha4 | 1.06E-12 | -0.38955 | 0.086 | 0.212 | 3.11E-08 | -0.16 | 1.67E-25 | 0.431699 | 0.209 | 0.047 | 3.04E-21 | 1.92 | |
| Col25a1 | 0.003101 | -0.30507 | 0.149 | 0.199 | 1 | -0.23 | 5.44E-11 | 0.191718 | 0.060 | 0.006 | 9.92E-07 | 1.92 | 3.04 |
| Cdkn1c | 0.226406 | -0.05258 | 0.060 | 0.081 | 1 | -0.04 | 9.39E-63 | 1.939491 | 0.910 | 0.938 | 1.71E-58 | 1.88 | |
| Arhgef28 | 0.183275 | -5.4E-15 | 0.022 | 0.011 | 1 | 0.00 | 6.16E-17 | 0.189021 | 0.099 | 0.010 | 1.12E-12 | 1.87 | 4.52 |
| Fam222a | 0.027255 | 0.066819 | 0.069 | 0.041 | 1 | 0.11 | 1.36E-15 | 0.182807 | 0.090 | 0.009 | 2.48E-11 | 1.83 | |

| | | | | | | | | | | | | | |
|---|---|---|---|---|---|---|---|---|---|---|---|---|---|
| Lancl3 | 0.702975 | -0.00149 | 0.004 | 0.003 | 1 | 0.00 | 1.01E-10 | 0.113116 | 0.048 | 0.003 | 1.84E-06 | 1.81 | 4.19 |
| Flrt2 | 5.6E-17 | -0.53972 | 0.497 | 0.697 | 1.64E-12 | -0.38 | 7.17E-46 | 0.791895 | 0.463 | 0.204 | 1.31E-41 | 1.80 | |
| Unc5c | 0.096793 | -0.01133 | 0.009 | 0.017 | 1 | -0.01 | 1.04E-08 | 0.098643 | 0.036 | 0.002 | 0.00019 | 1.78 | 1.62 |
| Zfand5 | 1.58E-08 | 0.207266 | 0.350 | 0.346 | 0.000465 | 0.21 | 3E-146 | 1.411027 | 0.928 | 0.740 | 5.4E-142 | 1.77 | |
| Gml | 0.313547 | -0.00407 | 0.000 | 0.002 | 1 | 0.00 | 4.55E-09 | 0.097985 | 0.036 | 0.002 | 8.29E-05 | 1.76 | |
| Mex3b | 8.01E-05 | 0.12824 | 0.077 | 0.044 | 1 | 0.22 | 1.81E-41 | 0.695329 | 0.412 | 0.163 | 3.29E-37 | 1.76 | 1.57 |
| Otof | 0.630919 | -0.00075 | 0.000 | 0.001 | 1 | 0.00 | 1.01E-07 | 0.064745 | 0.027 | 0.001 | 0.001845 | 1.75 | 5.02 |
| Il17rd | 2.19E-05 | 0.217466 | 0.179 | 0.108 | 0.643827 | 0.36 | 1.94E-29 | 0.446584 | 0.233 | 0.060 | 3.54E-25 | 1.73 | 1.47 |
| Tmod3 | 0.307931 | 0.048892 | 0.523 | 0.486 | 1 | 0.05 | 5E-94 | 1.209574 | 0.597 | 0.430 | 9.11E-90 | 1.68 | 3.09 |
| Col7a1 | 1.49E-11 | -0.13899 | 0.246 | 0.242 | 4.39E-07 | -0.14 | 2.82E-67 | 0.983017 | 0.701 | 0.412 | 5.15E-63 | 1.67 | 2.31 |
| Fsd1 | 0.261505 | 0.018554 | 0.011 | 0.006 | 1 | 0.03 | 2.14E-10 | 0.130911 | 0.051 | 0.004 | 3.9E-06 | 1.67 | |
| Alcam | 9.38E-18 | -0.20409 | 0.138 | 0.249 | 2.75E-13 | -0.11 | 2.39E-27 | 0.530252 | 0.319 | 0.103 | 4.35E-23 | 1.64 | |
| Tmsb4x | 0.041739 | 0.051592 | 0.251 | 0.204 | 1 | 0.06 | 4.1E-190 | 1.610085 | 1.000 | 0.982 | 7.5E-186 | 1.64 | |
| Leprot | 0.003764 | 0.07243 | 0.193 | 0.202 | 1 | 0.07 | 4.95E-98 | 1.0964 | 0.728 | 0.489 | 9.02E-94 | 1.63 | 1.91 |
| Epha7 | 0.543606 | 0.015083 | 0.031 | 0.023 | 1 | 0.02 | 6.17E-22 | 0.351284 | 0.161 | 0.035 | 1.12E-17 | 1.62 | 2.34 |
| Gdpd1 | 0.333978 | 0.061565 | 0.091 | 0.074 | 1 | 0.08 | 5.83E-15 | 0.106729 | 0.075 | 0.005 | 1.06E-10 | 1.60 | |
| Fbln2 | 1.04E-05 | 0.171286 | 0.106 | 0.075 | 0.304642 | 0.24 | 1.13E-54 | 0.906576 | 0.573 | 0.325 | 2.07E-50 | 1.60 | 2.13 |
| Ypel1 | 0.00218 | -0.08909 | 0.146 | 0.203 | 1 | -0.06 | 5.19E-23 | 0.371547 | 0.212 | 0.050 | 9.45E-19 | 1.58 | |
| Slitrk6 | 0.000601 | -0.08346 | 0.046 | 0.075 | 1 | -0.05 | 7.91E-37 | 0.837148 | 0.570 | 0.303 | 1.44E-32 | 1.57 | |
| Cd44 | 3.16E-08 | -0.45066 | 0.501 | 0.622 | 0.000928 | -0.36 | 8.25E-53 | 0.866379 | 0.555 | 0.307 | 1.5E-48 | 1.57 | |
| Cdh22 | 0.326068 | 0.007791 | 0.007 | 0.008 | 1 | 0.01 | 6.05E-08 | 0.117838 | 0.039 | 0.003 | 0.001103 | 1.53 | |
| Mef2c | 3.1E-137 | -1.56383 | 0.357 | 0.864 | 9E-133 | -0.65 | 4.52E-38 | 0.841608 | 0.707 | 0.389 | 8.24E-34 | 1.53 | |

**Supplementary Table 7: Shared IFE Enriched Genes. U score = avg_log2FC x pct.2/pct.1. U score < -1.5 for mouse and human.**

| | Human | | | | | | Mouse (Qu et al., 2022) | | | | | |
|---|---|---|---|---|---|---|---|---|---|---|---|---|
| | p_val | avg_log2FC | pct.1 | pct.2 | p_val_adj | UScore | p_val | avg_log2FC | pct.1 | pct.2 | p_val_adj | UScore |
| KRT15 | 3.59E-28 | -1.34049 | 0.541 | 0.875 | 1.05E-23 | -2.17 | 5.3E-190 | -1.92 | 0.7 | 0.965 | 9.7E-186 | -2.65 |
| P3H2 | 2.36E-31 | -1.31119 | 0.586 | 0.888 | 6.92E-27 | -1.99 | 3.87E-22 | -0.17151 | 0.01 | 0.144 | 7.05E-18 | -2.47 |

**Supplementary Table 8: Human Specific IFE Enriched Genes. U score = avg_log2FC x pct.2/pct.1. U score < -1.5 for human, > -0.5 for mouse.**

| | Human | | | | | | Mouse (Qu et al., 2022) | | | | | |
|---|---|---|---|---|---|---|---|---|---|---|---|---|
| | p_val | avg_log2FC | pct.1 | pct.2 | p_val_adj | UScore | p_val | avg_log2FC | pct.1 | pct.2 | p_val_adj | UScore |
| GRID2 | 3.71E-47 | -3.08023 | 0.083 | 0.646 | 1.09E-42 | -23.97 | 0.332976 | 0.000126 | 0.002 | 0 | 1 | 0.00 |
| FSTL5 | 2.90E-17 | -1.26856 | 0.032 | 0.305 | 8.52E-13 | -12.09 | 1 | 0 | 0 | 0 | 1 | NE |
| CSMD1 | 1.40E-58 | -3.03258 | 0.325 | 0.839 | 4.10E-54 | -7.83 | 0.669848 | -0.00253 | 0 | 0 | 1 | NE |
| ERG | 1.18E-17 | -1.01256 | 0.045 | 0.331 | 3.46E-13 | -7.45 | 0.364577 | 0 | 0 | 0.001 | 1 | NE |
| PCDH7 | 4.07E-68 | -2.43484 | 0.293 | 0.884 | 1.19E-63 | -7.35 | 0.037357 | -0.05099 | 0.181 | 0.23 | 1 | -0.06 |
| NRG3 | 3.56E-20 | -1.73779 | 0.115 | 0.478 | 1.04E-15 | -7.22 | 0.486631 | -0.00082 | 0 | 0.001 | 1 | NE |
| KRT14 | 9.30E-25 | -1.47589 | 0.127 | 0.511 | 2.73E-20 | -5.94 | 7.21E-06 | -0.22501 | 0.981 | 0.986 | 0.131394 | -0.23 |
| RALYL | 3.39E-50 | -2.26745 | 0.369 | 0.866 | 9.95E-46 | -5.32 | 1 | 0 | 0 | 0 | 1 | NE |
| SLC4A4 | 5.33E-40 | -2.0283 | 0.306 | 0.798 | 1.56E-35 | -5.29 | 0.065157 | -0.02559 | 0.015 | 0.033 | 1 | -0.06 |
| IGFBP6 | 9.22E-11 | -0.57417 | 0.025 | 0.22 | 2.71E-06 | -5.05 | 0.277795 | -3.2E-16 | 0 | 0.002 | 1 | NE |
| DOCK5 | 4.25E-18 | -0.93296 | 0.07 | 0.378 | 1.25E-13 | -5.04 | 0.277795 | -0.00108 | 0 | 0.002 | 1 | NE |
| ENPP1 | 5.09E-26 | -1.39379 | 0.197 | 0.63 | 1.49E-21 | -4.46 | 0.361478 | -0.00338 | 0.002 | 0.006 | 1 | -0.01 |
| FOS | 7.38E-27 | -1.63047 | 0.229 | 0.603 | 2.17E-22 | -4.29 | 0.000357 | -0.28181 | 0.365 | 0.419 | 1 | -0.32 |
| KHDRBS2 | 2.78E-40 | -1.70705 | 0.376 | 0.852 | 8.15E-36 | -3.87 | 0.332976 | -7.8E-10 | 0.002 | 0 | 1 | 0.00 |
| KRT19 | 6.67E-25 | -1.33728 | 0.229 | 0.642 | 1.96E-20 | -3.75 | 0.013718 | -0.00867 | 0.017 | 0.031 | 1 | -0.02 |
| NPR3 | 9.87E-24 | -1.26324 | 0.21 | 0.618 | 2.90E-19 | -3.72 | 0.002412 | -0.07277 | 0.102 | 0.162 | 1 | -0.12 |
| FAM155A | 1.78E-19 | -1.60242 | 0.28 | 0.644 | 5.22E-15 | -3.69 | 1 | 0 | 0 | 0 | 1 | NE |
| FOSB | 1.18E-17 | -1.18658 | 0.146 | 0.448 | 3.48E-13 | -3.64 | 0.050997 | -0.01972 | 0.048 | 0.073 | 1 | -0.03 |
| SYBU | 1.15E-13 | -0.67672 | 0.064 | 0.319 | 3.39E-09 | -3.37 | 0.364577 | -0.00079 | 0 | 0.001 | 1 | NE |
| PLA2R1 | 1.13E-49 | -1.70565 | 0.452 | 0.879 | 3.33E-45 | -3.32 | 0.189255 | -0.00072 | 0.002 | 0.004 | 1 | 0.00 |
| EML6 | 2.17E-32 | -1.38703 | 0.338 | 0.782 | 6.38E-28 | -3.21 | 0.990105 | -0.00132 | 0.008 | 0.009 | 1 | 0.00 |
| DNER | 5.11E-18 | -1.17209 | 0.21 | 0.556 | 1.50E-13 | -3.10 | 1 | 0 | 0 | 0 | 1 | NE |
| STAT4 | 1.81E-17 | -1.0453 | 0.166 | 0.49 | 5.32E-13 | -3.09 | 0.486631 | -0.00083 | 0 | 0.001 | 1 | NE |
| ADGRB3 | 1.22E-11 | -1.2389 | 0.153 | 0.371 | 3.58E-07 | -3.00 | 0.116 | 0 | 0.002 | 0 | 1 | 0.00 |
| RASSF6 | 5.47E-12 | -0.58249 | 0.051 | 0.258 | 1.61E-07 | -2.95 | 0.766699 | -0.00856 | 0.021 | 0.026 | 1 | -0.01 |
| PDLIM1 | 2.97E-28 | -1.23697 | 0.293 | 0.693 | 8.73E-24 | -2.93 | 1.19E-07 | -0.22525 | 0.785 | 0.847 | 0.002161 | -0.24 |
| PID1 | 3.10E-19 | -1.12519 | 0.217 | 0.542 | 9.11E-15 | -2.81 | 0.032077 | -0.00077 | 0.031 | 0.016 | 1 | 0.00 |
| EDIL3 | 8.43E-12 | -0.98784 | 0.134 | 0.375 | 2.47E-07 | -2.76 | 0.951391 | -1.6E-09 | 0.002 | 0.001 | 1 | 0.00 |

| | | | | | | | | | | | | |
|---|---|---|---|---|---|---|---|---|---|---|---|---|
| IFFO2 | 3.50E-10 | -0.75171 | 0.076 | 0.275 | 1.03E-05 | -2.72 | 0.000275 | -0.08172 | 0.058 | 0.117 | 1 | -0.16 |
| ADCY2 | 2.13E-21 | -1.19606 | 0.306 | 0.679 | 6.27E-17 | -2.65 | 0.550644 | 0 | 0.002 | 0.001 | 1 | 0.00 |
| MEF2C | 3.39E-15 | -0.95518 | 0.153 | 0.423 | 9.94E-11 | -2.64 | 0.998845 | 0 | 0.013 | 0.013 | 1 | 0.00 |
| EXPH5 | 4.60E-38 | -1.59169 | 0.535 | 0.865 | 1.35E-33 | -2.57 | 0.010331 | -0.03867 | 0.019 | 0.045 | 1 | -0.09 |
| FGF14 | 1.21E-14 | -0.85364 | 0.166 | 0.49 | 3.55E-10 | -2.52 | 1 | 0 | 0 | 0 | 1 | NE |
| FAT2 | 2.75E-19 | -0.86549 | 0.197 | 0.562 | 8.09E-15 | -2.47 | 1.3E-11 | -0.12486 | 0.05 | 0.159 | 2.37E-07 | -0.40 |
| PTCHD4 | 6.42E-31 | -1.27211 | 0.452 | 0.866 | 1.88E-26 | -2.44 | 1.94E-13 | 0.085592 | 0.046 | 0.002 | 3.53E-09 | 0.00 |
| NCAM2 | 3.53E-26 | -1.34243 | 0.459 | 0.833 | 1.04E-21 | -2.44 | 0.486631 | -1.3E-15 | 0 | 0.001 | 1 | NE |
| ARHGAP44 | 1.33E-26 | -1.17901 | 0.395 | 0.803 | 3.89E-22 | -2.40 | 0.046099 | 0.023322 | 0.021 | 0.008 | 1 | 0.01 |
| SOX6 | 1.84E-73 | -1.55062 | 0.662 | 0.997 | 5.39E-69 | -2.34 | 1.39E-10 | -0.25193 | 0.294 | 0.459 | 2.52E-06 | -0.39 |
| PKIB | 3.62E-10 | -0.67032 | 0.102 | 0.343 | 1.06E-05 | -2.25 | 0.669848 | 0 | 0 | 0 | 1 | NE |
| FAM189A2 | 5.89E-17 | -0.98318 | 0.255 | 0.581 | 1.73E-12 | -2.24 | 0.024979 | -0.00789 | 0 | 0.007 | 1 | NE |
| PPP2R2C | 1.01E-09 | -0.65297 | 0.089 | 0.3 | 2.96E-05 | -2.20 | 1 | 0 | 0 | 0 | 1 | NE |
| CRYBG3 | 1.53E-19 | -0.92198 | 0.268 | 0.632 | 4.51E-15 | -2.17 | 0.033307 | -0.03531 | 0.133 | 0.174 | 1 | -0.05 |
| CDCP1 | 8.40E-11 | -0.63649 | 0.102 | 0.347 | 2.47E-06 | -2.17 | 1.03E-10 | 0.248757 | 0.371 | 0.258 | 1.88E-06 | 0.17 |
| CNTNAP3 | 3.04E-06 | -0.29042 | 0.019 | 0.139 | 0.089154 | -2.12 | 1 | 0 | 0 | 0 | 1 | NE |
| LGALS3 | 5.43E-17 | -0.83143 | 0.197 | 0.481 | 1.59E-12 | -2.03 | 0.88718 | -0.00615 | 0.04 | 0.04 | 1 | -0.01 |
| PCSK6 | 7.10E-22 | -0.87894 | 0.306 | 0.706 | 2.08E-17 | -2.03 | 0.121871 | -0.0186 | 0.021 | 0.039 | 1 | -0.03 |
| POSTN | 4.56E-32 | -1.18866 | 0.529 | 0.89 | 1.34E-27 | -2.00 | 1.3E-17 | 0.403313 | 0.727 | 0.719 | 2.36E-13 | 0.40 |
| NGF | 7.89E-20 | -0.96554 | 0.357 | 0.717 | 2.32E-15 | -1.94 | 3.14E-07 | -0.08665 | 0.219 | 0.325 | 0.005714 | -0.13 |
| TMEM40 | 2.63E-09 | -0.53688 | 0.076 | 0.273 | 7.71E-05 | -1.93 | 0.46476 | -0.00887 | 0.01 | 0.018 | 1 | -0.02 |
| LSAMP | 2.71E-09 | -0.99052 | 0.217 | 0.415 | 7.95E-05 | -1.89 | 0.957822 | 0 | 0.002 | 0.003 | 1 | 0.00 |
| LRRC7 | 1.66E-55 | -1.36526 | 0.732 | 0.977 | 4.86E-51 | -1.82 | 0.129656 | -0.00462 | 0 | 0.003 | 1 | NE |
| JPH1 | 1.62E-08 | -0.47674 | 0.057 | 0.217 | 0.000476 | -1.81 | 0.255269 | -0.00778 | 0.004 | 0.012 | 1 | -0.02 |
| CCDC3 | 3.31E-15 | -1.05568 | 0.389 | 0.666 | 9.73E-11 | -1.81 | 5.65E-06 | 0.129904 | 0.194 | 0.114 | 0.103067 | 0.08 |
| NACC2 | 4.79E-06 | -0.30233 | 0.025 | 0.149 | 0.140732 | -1.80 | 0.18766 | -0.01235 | 0.008 | 0.019 | 1 | -0.03 |
| MYOF | 1.01E-36 | -1.14498 | 0.58 | 0.912 | 2.97E-32 | -1.80 | 0.219926 | -0.01211 | 0.01 | 0.02 | 1 | -0.02 |
| PRKG1 | 1.33E-35 | -1.36869 | 0.72 | 0.945 | 3.91E-31 | -1.80 | 0.669848 | 0 | 0 | 0 | 1 | NE |
| UNC5C | 2.86E-05 | -0.44949 | 0.038 | 0.151 | 0.838364 | -1.79 | 0.718621 | -0.00119 | 0.004 | 0.006 | 1 | 0.00 |
| BMPR1B | 2.73E-06 | -0.54069 | 0.07 | 0.226 | 0.080255 | -1.75 | 1.03E-05 | -0.0953 | 0.171 | 0.263 | 0.186928 | -0.15 |
| CELSR1 | 6.91E-22 | -0.92783 | 0.414 | 0.773 | 2.03E-17 | -1.73 | 3.03E-06 | -0.12515 | 0.36 | 0.482 | 0.055188 | -0.17 |
| GRIK1 | 4.56E-35 | -1.2426 | 0.688 | 0.958 | 1.34E-30 | -1.73 | 0.116 | 0.001423 | 0.002 | 0 | 1 | 0.00 |

| | | | | | | | | | | | | |
|---|---|---|---|---|---|---|---|---|---|---|---|---|
| UNC80 | 0.000126 | -0.23161 | 0.013 | 0.096 | 1 | -1.71 | 0.486631 | -0.00238 | 0 | 0.001 | 1 | NE |
| TRPC6 | 3.59E-16 | -0.95242 | 0.389 | 0.682 | 1.06E-11 | -1.67 | 1 | 0 | 0 | 0 | 1 | NE |
| RIC3 | 1.14E-12 | -0.76133 | 0.236 | 0.517 | 3.36E-08 | -1.67 | 0.189911 | -0.02087 | 0.021 | 0.036 | 1 | -0.04 |
| CPNE8 | 2.74E-12 | -0.81827 | 0.287 | 0.583 | 8.05E-08 | -1.66 | 0.04557 | -0.01434 | 0.054 | 0.077 | 1 | -0.02 |
| NTN4 | 1.07E-07 | -0.53276 | 0.083 | 0.256 | 0.003143 | -1.64 | 8.96E-05 | -0.07196 | 0.025 | 0.073 | 1 | -0.21 |
| ADGRG6 | 1.04E-10 | -0.64637 | 0.185 | 0.456 | 3.06E-06 | -1.59 | 0.272518 | -0.00664 | 0.008 | 0.018 | 1 | -0.01 |
| PLXNA2 | 1.65E-08 | -0.59841 | 0.121 | 0.322 | 0.000485 | -1.59 | 0.000719 | -0.0789 | 0.088 | 0.151 | 1 | -0.14 |
| SLC24A3 | 2.43E-10 | -0.80502 | 0.242 | 0.465 | 7.14E-06 | -1.55 | 0.923938 | 0.000298 | 0.013 | 0.013 | 1 | 0.00 |
| ROBO2 | 1.89E-38 | -1.31157 | 0.809 | 0.951 | 5.56E-34 | -1.54 | 0.011093 | 0.056791 | 0.117 | 0.076 | 1 | 0.04 |
| TBC1D4 | 1.15E-10 | -0.75082 | 0.248 | 0.508 | 3.37E-06 | -1.54 | 0.089542 | -0.01456 | 0.008 | 0.022 | 1 | -0.04 |
| EFNA5 | 2.52E-28 | -1.12098 | 0.688 | 0.935 | 7.41E-24 | -1.52 | 0.000107 | 0.073754 | 0.331 | 0.357 | 1 | 0.08 |
| SLITRK5 | 2.42E-07 | -0.39006 | 0.057 | 0.221 | 0.007102 | -1.51 | 0.332976 | 0.002492 | 0.002 | 0 | 1 | 0.00 |

**Supplementary Table 9: Mouse Specific IFE Enriched Genes. U score = avg_log2FC x pct.2/pct.1. U score < -1.5 for mouse, > -0.5 for human.**

| | Human | | | | | | Mouse (Qu et al., 2022) | | | | | |
|---|---|---|---|---|---|---|---|---|---|---|---|---|
| | p_val | avg_log2FC | pct.1 | pct.2 | p_val_adj | UScore | p_val | avg_log2FC | pct.1 | pct.2 | p_val_adj | UScore |
| Defb1 | 0.72313 | -0.00105 | 0 | 0.001 | 1 | NE | 8.78E-23 | -0.23245 | 0.01 | 0.146 | 1.60E-18 | -3.39 |
| Edn1 | 0.153251 | 0.104611 | 0.057 | 0.027 | 1 | 0.05 | 1.02E-15 | -0.13245 | 0.004 | 0.094 | 1.86E-11 | -3.11 |
| Mgll | 0.148937 | 0.170967 | 0.102 | 0.063 | 1 | 0.11 | 4.37E-33 | -0.26394 | 0.019 | 0.217 | 7.97E-29 | -3.01 |
| Gas1 | 0.326599 | -0.01164 | 0.051 | 0.039 | 1 | -0.01 | 1.12E-112 | -1.25782 | 0.373 | 0.849 | 2.03E-108 | -2.86 |
| Dlk2 | 6.86E-05 | -0.1367 | 0 | 0.056 | 1 | NE | 1.38E-34 | -0.33756 | 0.033 | 0.258 | 2.52E-30 | -2.64 |
| Ndrg1 | 0.203185 | 0.140154 | 0.178 | 0.189 | 1 | 0.15 | 6.65E-33 | -0.31111 | 0.033 | 0.25 | 1.21E-28 | -2.36 |
| Igfbp5 | 0.015797 | -0.17203 | 0.083 | 0.114 | 1 | -0.24 | 3.59E-92 | -1.16081 | 0.417 | 0.829 | 6.55E-88 | -2.31 |
| Dst | 0.032437 | -0.27546 | 0.943 | 0.96 | 1 | -0.28 | 1.19E-74 | -1.04149 | 0.354 | 0.747 | 2.17E-70 | -2.20 |
| Col12a1 | 0.000136 | -0.37085 | 0.446 | 0.28 | 1 | -0.23 | 1.30E-35 | -0.3656 | 0.058 | 0.307 | 2.38E-31 | -1.94 |
| Wnt4 | 0.081596 | -0.14138 | 0.096 | 0.156 | 1 | -0.23 | 1.56E-66 | -0.81907 | 0.327 | 0.717 | 2.84E-62 | -1.80 |
| Chchd10 | 0.590731 | 0.008547 | 0.013 | 0.005 | 1 | 0.00 | 3.72E-64 | -0.715 | 0.277 | 0.685 | 6.79E-60 | -1.77 |
| Vcan | 7.97E-25 | 0.463385 | 0.516 | 0.126 | 2.34E-20 | 0.11 | 6.16E-51 | -0.90935 | 0.348 | 0.675 | 1.12E-46 | -1.76 |
| Fgfbp1 | 0.258632 | 0.030691 | 0.013 | 0.003 | 1 | 0.01 | 1.31E-76 | -0.98953 | 0.481 | 0.839 | 2.39E-72 | -1.73 |
| Hotairm1 | 0.19983 | -0.09758 | 0.172 | 0.235 | 1 | -0.13 | 2.20E-50 | -0.75584 | 0.296 | 0.656 | 4.00E-46 | -1.68 |
| Wnt16 | 0.090484 | 0.053986 | 0.096 | 0.052 | 1 | 0.03 | 6.07E-33 | -0.48306 | 0.112 | 0.381 | 1.11E-28 | -1.64 |

**Supplementary Table 10: Shared Dermis Enriched Genes. U score = avg_log2FC x pct.2/pct.1. U score < -1.5 for mouse and human.**

| | Human | | | | | | Mouse (Qu et al., 2022) | | | | | |
|---|---|---|---|---|---|---|---|---|---|---|---|---|
| | p_val | avg_log2FC | pct.1 | pct.2 | p_val_adj | UScore | p_val | avg_log2FC | pct.1 | pct.2 | p_val_adj | UScore |
| KCNK2 | 8.23E-103 | -1.6996 | 0.018 | 0.449 | 2.42E-98 | -42.40 | 7.85E-28 | -0.71124 | 0.03 | 0.26 | 1.43E-23 | -6.16 |
| GREM2 | 4.38E-112 | -1.73808 | 0.151 | 0.675 | 1.29E-107 | -7.77 | 2.37E-57 | -0.82919 | 0.039 | 0.441 | 4.31E-53 | -9.38 |
| NFIA | 1.54E-213 | -2.34429 | 0.434 | 0.932 | 4.53E-209 | -5.03 | 7E-65 | -1.23615 | 0.122 | 0.591 | 1.28E-60 | -5.99 |
| FAP | 2.24E-38 | -0.63475 | 0.04 | 0.259 | 6.57E-34 | -4.11 | 2.21E-14 | -0.18439 | 0.003 | 0.103 | 4.03E-10 | -6.33 |
| ABI3BP | 2.65E-202 | -1.87159 | 0.557 | 0.974 | 7.79E-198 | -3.27 | 6.29E-12 | -0.2156 | 0.012 | 0.115 | 1.15E-07 | -2.07 |
| ARHGAP6 | 6.90E-52 | -0.93312 | 0.213 | 0.573 | 2.02E-47 | -2.51 | 8.48E-12 | -0.2024 | 0.012 | 0.115 | 1.55E-07 | -1.94 |
| EFNA5 | 3.00E-206 | -1.86875 | 0.75 | 0.983 | 8.82E-202 | -2.45 | 3.47E-39 | -0.70342 | 0.299 | 0.678 | 6.32E-35 | -1.60 |
| ADAMTSL1 | 4.00E-73 | -1.15649 | 0.388 | 0.765 | 1.18E-68 | -2.28 | 8.72E-23 | -0.37503 | 0.03 | 0.231 | 1.59E-18 | -2.89 |
| VGLL3 | 3.26E-28 | -0.65847 | 0.107 | 0.325 | 9.57E-24 | -2.00 | 5.4E-13 | -0.21789 | 0.012 | 0.124 | 9.84E-09 | -2.25 |
| CREB3L1 | 5.70E-27 | -0.40008 | 0.047 | 0.231 | 1.67E-22 | -1.97 | 5.57E-53 | -0.97306 | 0.149 | 0.578 | 1.02E-48 | -3.77 |
| ITGA4 | 7.52E-18 | -0.34098 | 0.031 | 0.155 | 2.21E-13 | -1.70 | 1.2E-16 | -0.27184 | 0.018 | 0.163 | 2.18E-12 | -2.46 |

**Supplementary Table 11: Human Specific Dermis Enriched Genes. U score = avg_log2FC x pct.2/pct.1. U score < -1.5 for human, > -0.5 for mouse.**

| | Human | | | | | | Mouse (Qu et al., 2022) | | | | | |
|---|---|---|---|---|---|---|---|---|---|---|---|---|
| | p_val | avg_log2FC | pct.1 | pct.2 | p_val_adj | UScore | p_val | avg_log2FC | pct.1 | pct.2 | p_val_adj | UScore |
| FSTL5 | 1.00E-123 | -2.45655 | 0.038 | 0.537 | 2.94E-119 | -34.71 | 0.326373758 | 0.006738 | 0.003 | 0.001 | 1 | 0.00 |
| SPHKAP | 1.59E-138 | -2.42746 | 0.046 | 0.586 | 4.67E-134 | -30.92 | 1.16317E-35 | 0.740929 | 0.343 | 0.12 | 2.12012E-31 | 0.26 |
| CPNE4 | 5.40E-61 | -1.61688 | 0.02 | 0.31 | 1.58E-56 | -25.06 | 0.326373758 | -0.00058 | 0.003 | 0.001 | 1 | 0.00 |
| SEMA3A | 5.77E-197 | -2.62606 | 0.104 | 0.753 | 1.69E-192 | -19.01 | 0.006701553 | -0.05272 | 0.116 | 0.163 | 1 | -0.07 |
| PLXNA4 | 4.60E-79 | -1.16992 | 0.026 | 0.383 | 1.35E-74 | -17.23 | 2.14676E-12 | 0.332481 | 0.293 | 0.237 | 3.91291E-08 | 0.27 |
| DOK5 | 4.49E-108 | -1.76145 | 0.071 | 0.549 | 1.32E-103 | -13.62 | 0.034051368 | 0.082012 | 0.113 | 0.097 | 1 | 0.07 |
| ATP10A | 5.82E-101 | -1.39923 | 0.053 | 0.507 | 1.71E-96 | -13.39 | 0.245354298 | -0.00803 | 0.003 | 0.009 | 1 | -0.02 |
| WSCD2 | 2.27E-18 | -0.26361 | 0.002 | 0.09 | 6.66E-14 | -11.86 | 0.038279119 | -0.01594 | 0.116 | 0.144 | 1 | -0.02 |
| PLEKHG1 | 3.49E-175 | -2.33164 | 0.151 | 0.754 | 1.02E-170 | -11.64 | 4.24689E-06 | 0.138563 | 0.116 | 0.079 | 0.077407989 | 0.09 |
| AKAP6 | 2.91E-99 | -2.00229 | 0.104 | 0.574 | 8.54E-95 | -11.05 | 0.657070963 | -0.00357 | 0.003 | 0.005 | 1 | -0.01 |
| ETV1 | 7.28E-81 | -1.12247 | 0.046 | 0.432 | 2.14E-76 | -10.54 | 0.000354153 | 0.139666 | 0.125 | 0.085 | 1 | 0.09 |
| PID1 | 1.80E-112 | -2.33772 | 0.135 | 0.599 | 5.29E-108 | -10.37 | 0.000264943 | -0.22612 | 0.212 | 0.317 | 1 | -0.34 |
| PVT1 | 1.85E-123 | -1.51337 | 0.098 | 0.628 | 5.43E-119 | -9.70 | 0.435114391 | 0.019952 | 0.051 | 0.045 | 1 | 0.02 |
| DCHS2 | 4.84E-151 | -1.91661 | 0.151 | 0.737 | 1.42E-146 | -9.35 | 0.027549034 | -0.03346 | 0.003 | 0.02 | 1 | -0.22 |
| GPC6 | 2.67E-252 | -3.21604 | 0.326 | 0.888 | 7.84E-248 | -8.76 | 0.02524238 | -0.07319 | 0.087 | 0.129 | 1 | -0.11 |
| STXBP6 | 7.80E-51 | -0.91876 | 0.033 | 0.283 | 2.29E-46 | -7.88 | 0.175632543 | 0.06056 | 0.128 | 0.098 | 1 | 0.05 |
| PDGFD | 1.05E-111 | -2.11253 | 0.186 | 0.662 | 3.10E-107 | -7.52 | 0.332171339 | -0.00528 | 0.003 | 0.007 | 1 | -0.01 |
| GXYLT2 | 5.27E-93 | -1.27211 | 0.093 | 0.542 | 1.55E-88 | -7.41 | 2.40261E-06 | -0.17734 | 0.275 | 0.407 | 0.043792311 | -0.26 |
| PDE2A | 1.10E-22 | -0.286 | 0.005 | 0.118 | 3.23E-18 | -6.75 | 0.001120463 | 7.93E-08 | 0.024 | 0.003 | 1 | 0.00 |
| MTSS1 | 8.98E-123 | -1.78022 | 0.186 | 0.704 | 2.64E-118 | -6.74 | 0.274410136 | 0.047422 | 0.152 | 0.121 | 1 | 0.04 |
| GLIS3 | 5.80E-75 | -1.63116 | 0.118 | 0.487 | 1.70E-70 | -6.73 | 0.008397736 | 0.016935 | 0.006 | 0.002 | 1 | 0.01 |
| ARHGAP26 | 4.02E-108 | -1.91803 | 0.191 | 0.651 | 1.18E-103 | -6.54 | 0.000156071 | 0.112279 | 0.084 | 0.046 | 1 | 0.06 |
| SYNDIG1 | 4.09E-105 | -1.34532 | 0.137 | 0.642 | 1.20E-100 | -6.30 | 7.54198E-06 | 0.080494 | 0.045 | 0.008 | 0.137467643 | 0.01 |
| DPP10 | 4.74E-35 | -0.99561 | 0.036 | 0.222 | 1.39E-30 | -6.14 | 0.070755385 | 0.006863 | 0.003 | 0 | 1 | 0.00 |
| HECW1 | 5.78E-76 | -1.25312 | 0.107 | 0.52 | 1.70E-71 | -6.09 | 0.091181007 | 0.037204 | 0.015 | 0.007 | 1 | 0.02 |
| ANGPT1 | 5.55E-35 | -0.76068 | 0.029 | 0.232 | 1.63E-30 | -6.09 | 0.041025993 | 0.007635 | 0.009 | 0.01 | 1 | 0.01 |
| NRXN1 | 1.54E-137 | -1.89274 | 0.244 | 0.757 | 4.53E-133 | -5.87 | 0.004818933 | 0.071912 | 0.033 | 0.01 | 1 | 0.02 |

| | | | | | | | | | | | | |
|---|---|---|---|---|---|---|---|---|---|---|---|---|
| ETV5 | 1.08E-71 | -1.01152 | 0.087 | 0.478 | 3.18E-67 | -5.56 | 0.67635861 | -0.01595 | 0.167 | 0.181 | 1 | -0.02 |
| MITF | 2.74E-75 | -1.32331 | 0.122 | 0.506 | 8.03E-71 | -5.49 | 0.962146777 | 0.003385 | 0.024 | 0.022 | 1 | 0.00 |
| PRUNE2 | 8.36E-31 | -0.65046 | 0.024 | 0.201 | 2.45E-26 | -5.45 | 0.070755385 | 6.41E-16 | 0.003 | 0 | 1 | 0.00 |
| RHOJ | 6.78E-111 | -1.6432 | 0.213 | 0.693 | 1.99E-106 | -5.35 | 1.06778E-07 | -0.07733 | 0.373 | 0.502 | 0.001946237 | -0.10 |
| GUCY1A1 | 5.13E-28 | -0.53524 | 0.018 | 0.178 | 1.51E-23 | -5.29 | 0.326635958 | 0.007167 | 0.009 | 0.008 | 1 | 0.01 |
| RGMB | 9.39E-103 | -1.43768 | 0.191 | 0.669 | 2.76E-98 | -5.04 | 0.03598336 | -0.05614 | 0.287 | 0.346 | 1 | -0.07 |
| MPP7 | 1.38E-56 | -1.11638 | 0.093 | 0.418 | 4.04E-52 | -5.02 | 0.813136616 | -0.00614 | 0.021 | 0.024 | 1 | -0.01 |
| PDE7B | 3.42E-100 | -2.0486 | 0.293 | 0.707 | 1.01E-95 | -4.94 | 1.04832E-09 | 0.348772 | 0.379 | 0.259 | 1.91077E-05 | 0.24 |
| IL1RAPL1 | 1.52E-102 | -1.78268 | 0.266 | 0.737 | 4.46E-98 | -4.94 | 0.00220736 | 0.03194 | 0.018 | 0.002 | 1 | 0.00 |
| EFNB2 | 6.64E-81 | -1.26122 | 0.146 | 0.568 | 1.95E-76 | -4.91 | 0.001282495 | 0.128735 | 0.137 | 0.086 | 1 | 0.08 |
| SASH1 | 1.63E-136 | -1.72394 | 0.271 | 0.763 | 4.78E-132 | -4.85 | 0.0284212 | -0.08205 | 0.224 | 0.287 | 1 | -0.11 |
| CPED1 | 5.25E-221 | -2.23208 | 0.43 | 0.929 | 1.54E-216 | -4.82 | 5.95507E-11 | 0.299103 | 0.504 | 0.471 | 1.08543E-06 | 0.28 |
| L3MBTL4 | 3.06E-58 | -1.10303 | 0.117 | 0.465 | 8.99E-54 | -4.38 | 0.755729821 | 0.004025 | 0.003 | 0.001 | 1 | 0.00 |
| ZNF536 | 1.62E-137 | -1.65126 | 0.313 | 0.829 | 4.75E-133 | -4.37 | 0.000471562 | 0.076899 | 0.263 | 0.275 | 1 | 0.08 |
| HUNK | 4.16E-95 | -1.32088 | 0.206 | 0.682 | 1.22E-90 | -4.37 | 0.019244996 | -0.04668 | 0.209 | 0.258 | 1 | -0.06 |
| PHACTR1 | 1.19E-52 | -1.21544 | 0.122 | 0.433 | 3.50E-48 | -4.31 | 0.017701462 | -0.02224 | 0.024 | 0.045 | 1 | -0.04 |
| LIMCH1 | 6.40E-67 | -1.35341 | 0.169 | 0.537 | 1.88E-62 | -4.30 | 0.016687687 | 0.002661 | 0.012 | 0.02 | 1 | 0.00 |
| ARL4C | 9.52E-29 | -0.37436 | 0.016 | 0.177 | 2.79E-24 | -4.14 | 3.20243E-14 | 0.186457 | 0.352 | 0.423 | 5.83707E-10 | 0.22 |
| NEGR1 | 9.77E-63 | -1.73764 | 0.277 | 0.655 | 2.87E-58 | -4.11 | 0.023669927 | 0.045286 | 0.024 | 0.008 | 1 | 0.02 |
| ADGRA2 | 3.51E-62 | -0.80165 | 0.089 | 0.45 | 1.03E-57 | -4.05 | 3.48564E-06 | -0.13488 | 0.161 | 0.263 | 0.063532847 | -0.22 |
| ANKRD44 | 3.27E-97 | -1.26012 | 0.222 | 0.703 | 9.61E-93 | -3.99 | 5.34122E-11 | 0.276317 | 0.26 | 0.116 | 9.73544E-07 | 0.12 |
| RGL1 | 3.50E-65 | -0.99836 | 0.129 | 0.505 | 1.03E-60 | -3.91 | 5.74415E-07 | 0.18041 | 0.164 | 0.123 | 0.010469856 | 0.14 |
| MYOF | 1.23E-99 | -1.34993 | 0.242 | 0.697 | 3.60E-95 | -3.89 | 0.10396726 | -0.03336 | 0.027 | 0.048 | 1 | -0.06 |
| PELI2 | 1.65E-103 | -1.58929 | 0.317 | 0.759 | 4.84E-99 | -3.81 | 0.01833482 | 0.006705 | 0.054 | 0.07 | 1 | 0.01 |
| MEF2C | 3.08E-137 | -1.56383 | 0.357 | 0.864 | 9.05E-133 | -3.78 | 4.5231E-38 | 0.841608 | 0.707 | 0.389 | 8.24426E-34 | 0.46 |
| DOCK10 | 2.09E-132 | -1.58822 | 0.355 | 0.841 | 6.13E-128 | -3.76 | 0.432762986 | 0.046523 | 0.143 | 0.124 | 1 | 0.04 |
| COL13A1 | 1.11E-37 | -0.62234 | 0.046 | 0.278 | 3.26E-33 | -3.76 | 0.156572157 | 0.013819 | 0.009 | 0.002 | 1 | 0.00 |
| DENND2A | 5.70E-110 | -1.80691 | 0.372 | 0.762 | 1.67E-105 | -3.70 | 1.60577E-13 | 0.299924 | 0.224 | 0.132 | 2.92683E-09 | 0.18 |
| COL4A6 | 9.12E-27 | -0.45783 | 0.024 | 0.184 | 2.68E-22 | -3.51 | 0.11931498 | -0.01607 | 0.003 | 0.015 | 1 | -0.08 |
| SLC24A4 | 2.79E-09 | -0.14398 | 0.002 | 0.048 | 8.18E-05 | -3.46 | 0.733042545 | 0.003543 | 0.003 | 0.001 | 1 | 0.00 |
| FMN2 | 1.66E-28 | -0.56662 | 0.038 | 0.224 | 4.87E-24 | -3.34 | 0.993292363 | -0.00075 | 0.042 | 0.041 | 1 | 0.00 |
| F13A1 | 3.64E-31 | -0.7356 | 0.06 | 0.272 | 1.07E-26 | -3.33 | 0.422861724 | 0 | 0.003 | 0.009 | 1 | 0.00 |

| | | | | | | | | | | | | |
|---|---|---|---|---|---|---|---|---|---|---|---|---|
| KANK1 | 1.24E-71 | -1.0908 | 0.168 | 0.513 | 3.65E-67 | -3.33 | 0.069782263 | -0.02728 | 0.006 | 0.022 | 1 | -0.10 |
| ADAM12 | 2.15E-89 | -1.63335 | 0.386 | 0.783 | 6.33E-85 | -3.31 | 0.843761538 | -0.00883 | 0.009 | 0.012 | 1 | -0.01 |
| CACNA1C | 4.56E-78 | -1.14744 | 0.239 | 0.68 | 1.34E-73 | -3.26 | 0.002638199 | 0.036872 | 0.039 | 0.044 | 1 | 0.04 |
| APBB2 | 1.45E-200 | -1.82983 | 0.539 | 0.946 | 4.27E-196 | -3.21 | 1.27458E-05 | -0.15584 | 0.152 | 0.261 | 0.232317646 | -0.27 |
| SYNJ2 | 3.20E-44 | -0.6935 | 0.078 | 0.361 | 9.41E-40 | -3.21 | 0.005784672 | 0.041113 | 0.036 | 0.011 | 1 | 0.01 |
| TOX | 5.34E-56 | -1.19164 | 0.184 | 0.487 | 1.57E-51 | -3.15 | 0.09894147 | -0.01853 | 0.024 | 0.039 | 1 | -0.03 |
| RIPOR2 | 4.13E-104 | -1.22751 | 0.311 | 0.797 | 1.21E-99 | -3.15 | 6.04266E-05 | 0.11387 | 0.096 | 0.066 | 1 | 0.08 |
| KCNB1 | 1.28E-27 | -0.52049 | 0.036 | 0.216 | 3.76E-23 | -3.12 | 0.143361625 | -0.02188 | 0.006 | 0.018 | 1 | -0.07 |
| OSBP2 | 1.01E-61 | -1.07506 | 0.206 | 0.593 | 2.96E-57 | -3.09 | 0.326373758 | 0.002275 | 0.003 | 0.001 | 1 | 0.00 |
| ANXA11 | 3.05E-58 | -0.85894 | 0.138 | 0.497 | 8.95E-54 | -3.09 | 0.343821258 | 0.006489 | 0.087 | 0.091 | 1 | 0.01 |
| SYBU | 2.63E-35 | -0.55978 | 0.051 | 0.279 | 7.72E-31 | -3.06 | 0.005881707 | -0.03393 | 0.021 | 0.046 | 1 | -0.07 |
| EEPD1 | 5.17E-66 | -1.07059 | 0.219 | 0.618 | 1.52E-61 | -3.02 | 0.000240533 | -0.12306 | 0.051 | 0.118 | 1 | -0.28 |
| SCN8A | 2.93E-63 | -0.96896 | 0.189 | 0.586 | 8.62E-59 | -3.00 | 0.310673498 | 0.029883 | 0.033 | 0.021 | 1 | 0.02 |
| ZFP36L1 | 8.38E-75 | -1.13922 | 0.251 | 0.657 | 2.46E-70 | -2.98 | 6.38437E-09 | -0.23504 | 0.77 | 0.89 | 0.000116368 | -0.27 |
| DKK2 | 2.98E-29 | -0.7561 | 0.071 | 0.28 | 8.74E-25 | -2.98 | 0.137671894 | 0.03675 | 0.021 | 0.01 | 1 | 0.02 |
| NFATC2 | 7.26E-28 | -0.76441 | 0.066 | 0.255 | 2.13E-23 | -2.95 | 0.02587202 | 0.003872 | 0.003 | 0.002 | 1 | 0.00 |
| NAV3 | 1.40E-33 | -1.13509 | 0.128 | 0.33 | 4.11E-29 | -2.93 | 0.000175023 | -0.11677 | 0.024 | 0.079 | 1 | -0.38 |
| CMTM8 | 5.38E-18 | -0.4906 | 0.022 | 0.131 | 1.58E-13 | -2.92 | 0.000717564 | 0.047304 | 0.045 | 0.012 | 1 | 0.01 |
| SHROOM3 | 1.85E-55 | -0.94356 | 0.173 | 0.533 | 5.44E-51 | -2.91 | 0.031950726 | 0.036011 | 0.03 | 0.015 | 1 | 0.02 |
| CMKLR1 | 1.19E-11 | -0.17847 | 0.004 | 0.065 | 3.49E-07 | -2.90 | 0.900248149 | 0.008228 | 0.027 | 0.023 | 1 | 0.01 |
| NTRK3 | 1.40E-23 | -0.54086 | 0.031 | 0.165 | 4.12E-19 | -2.88 | 0.152595708 | 0.006814 | 0.009 | 0.013 | 1 | 0.01 |
| SLC16A7 | 4.99E-45 | -0.83569 | 0.129 | 0.442 | 1.47E-40 | -2.86 | 0.916351045 | -0.00182 | 0.012 | 0.012 | 1 | 0.00 |
| RBPMS | 1.85E-51 | -1.06705 | 0.202 | 0.538 | 5.42E-47 | -2.84 | 0.496574705 | -0.01893 | 0.048 | 0.061 | 1 | -0.02 |
| RIN3 | 3.41E-37 | -0.60831 | 0.067 | 0.312 | 1.00E-32 | -2.83 | 0.626355573 | -0.00495 | 0.009 | 0.013 | 1 | -0.01 |
| PLXNA2 | 8.34E-23 | -0.30038 | 0.016 | 0.15 | 2.45E-18 | -2.82 | 0.152065178 | -0.00614 | 0.012 | 0.028 | 1 | -0.01 |
| TTC39B | 6.49E-45 | -0.87351 | 0.144 | 0.461 | 1.90E-40 | -2.80 | 0.016353631 | -0.03357 | 0.003 | 0.022 | 1 | -0.25 |
| SRGAP1 | 1.09E-81 | -1.28834 | 0.352 | 0.76 | 3.20E-77 | -2.78 | 0.009371026 | 0.008056 | 0.116 | 0.138 | 1 | 0.01 |
| MASP1 | 1.71E-18 | -0.29572 | 0.013 | 0.122 | 5.02E-14 | -2.78 | 0.802410383 | -0.00261 | 0.003 | 0.005 | 1 | 0.00 |
| TES | 2.28E-42 | -0.82906 | 0.126 | 0.409 | 6.69E-38 | -2.69 | 0.001057583 | 0.053083 | 0.134 | 0.157 | 1 | 0.06 |
| PEAK1 | 4.10E-89 | -1.23307 | 0.359 | 0.783 | 1.20E-84 | -2.69 | 2.83938E-06 | -0.13609 | 0.185 | 0.296 | 0.051753434 | -0.22 |
| CADM2 | 5.79E-16 | -0.89707 | 0.067 | 0.2 | 1.70E-11 | -2.68 | 0.49717574 | 0.002803 | 0.003 | 0.001 | 1 | 0.00 |
| SFMBT2 | 1.47E-33 | -0.73425 | 0.087 | 0.317 | 4.31E-29 | -2.68 | 0.552283544 | 0.020024 | 0.048 | 0.043 | 1 | 0.02 |

| | | | | | | | | | | | | |
|---|---|---|---|---|---|---|---|---|---|---|---|---|
| ARHGAP28 | 1.47E-90 | -1.36708 | 0.423 | 0.824 | 4.31E-86 | -2.66 | 2.08325E-05 | -0.07701 | 0.281 | 0.376 | 0.379713783 | -0.10 |
| PLPPR1 | 6.20E-22 | -0.5645 | 0.042 | 0.198 | 1.82E-17 | -2.66 | 0.744903961 | -8.6E-05 | 0.009 | 0.007 | 1 | 0.00 |
| SHC4 | 4.35E-22 | -0.34966 | 0.02 | 0.15 | 1.28E-17 | -2.62 | 0.615291595 | 0.000903 | 0.012 | 0.009 | 1 | 0.00 |
| DAPK1 | 3.98E-93 | -1.35399 | 0.43 | 0.83 | 1.17E-88 | -2.61 | 0.07782179 | 0.077745 | 0.182 | 0.153 | 1 | 0.07 |
| LIN7A | 1.92E-49 | -0.78306 | 0.144 | 0.48 | 5.65E-45 | -2.61 | 1.02789E-23 | 0.612768 | 0.337 | 0.177 | 1.87353E-19 | 0.32 |
| ADCY1 | 2.22E-38 | -0.73578 | 0.102 | 0.359 | 6.53E-34 | -2.59 | 5.08872E-07 | 0.080904 | 0.036 | 0.014 | 0.00927521 | 0.03 |
| MGLL | 1.86E-45 | -0.79674 | 0.144 | 0.464 | 5.45E-41 | -2.57 | 4.63164E-19 | 0.208065 | 0.155 | 0.027 | 8.44208E-15 | 0.04 |
| NAV1 | 1.02E-71 | -1.24394 | 0.362 | 0.736 | 3.00E-67 | -2.53 | 2.96977E-05 | -0.12972 | 0.039 | 0.111 | 0.541300482 | -0.37 |
| LIMA1 | 2.51E-65 | -1.0371 | 0.27 | 0.657 | 7.38E-61 | -2.52 | 0.091176578 | -0.04476 | 0.549 | 0.604 | 1 | -0.05 |
| TGFBR3 | 2.58E-84 | -1.17827 | 0.372 | 0.79 | 7.56E-80 | -2.50 | 2.43422E-05 | 0.131397 | 0.391 | 0.395 | 0.44368475 | 0.13 |
| DPY19L1 | 8.23E-61 | -0.96606 | 0.244 | 0.627 | 2.42E-56 | -2.48 | 5.02768E-05 | -0.09044 | 0.045 | 0.106 | 0.91639444 | -0.21 |
| PALM2-AKAP2 | 4.10E-101 | -1.5105 | 0.525 | 0.859 | 1.20E-96 | -2.47 | 0.943421068 | -0.00225 | 0.003 | 0.004 | 1 | 0.00 |
| DLC1 | 6.04E-161 | -1.61704 | 0.627 | 0.952 | 1.77E-156 | -2.46 | 8.98377E-12 | -0.2723 | 0.316 | 0.515 | 1.63747E-07 | -0.44 |
| CYP26B1 | 3.39E-22 | -0.37556 | 0.026 | 0.169 | 9.94E-18 | -2.44 | 4.2871E-144 | 1.724228 | 0.612 | 0.087 | 7.8141E-140 | 0.25 |
| SLC7A11 | 2.53E-11 | -0.17561 | 0.005 | 0.069 | 7.42E-07 | -2.42 | 0.493855787 | -2.2E-14 | 0.003 | 0.008 | 1 | 0.00 |
| PLSCR1 | 2.79E-32 | -0.61946 | 0.078 | 0.305 | 8.18E-28 | -2.42 | 0.000141195 | -0.04667 | 0.11 | 0.169 | 1 | -0.07 |
| LHFPL2 | 1.19E-58 | -1.16624 | 0.328 | 0.681 | 3.49E-54 | -2.42 | 9.08262E-08 | -0.17012 | 0.146 | 0.269 | 0.001655489 | -0.31 |
| BCL11B | 5.40E-80 | -1.11209 | 0.355 | 0.771 | 1.58E-75 | -2.42 | 4.71139E-05 | -0.02395 | 0.707 | 0.808 | 0.858744946 | -0.03 |
| NCKAP5 | 1.01E-48 | -1.30533 | 0.304 | 0.562 | 2.96E-44 | -2.41 | 0.704074782 | -0.009 | 0.009 | 0.012 | 1 | -0.01 |
| SEMA3C | 2.48E-32 | -0.73873 | 0.095 | 0.309 | 7.29E-28 | -2.40 | 0.001439846 | 0.090076 | 0.036 | 0.014 | 1 | 0.04 |
| WIF1 | 5.91E-44 | -0.99841 | 0.235 | 0.565 | 1.74E-39 | -2.40 | 1.25139E-35 | 0.799541 | 0.701 | 0.559 | 2.2809E-31 | 0.64 |
| CNTN4 | 2.29E-100 | -1.61124 | 0.599 | 0.887 | 6.74E-96 | -2.39 | 2.08726E-06 | -0.12127 | 0.03 | 0.098 | 0.038044424 | -0.40 |
| PHACTR2 | 9.55E-33 | -0.77402 | 0.12 | 0.366 | 2.80E-28 | -2.36 | 0.00365529 | 0.007776 | 0.167 | 0.195 | 1 | 0.01 |
| ANK3 | 1.88E-38 | -0.66724 | 0.077 | 0.27 | 5.51E-34 | -2.34 | 0.045133613 | -0.08219 | 0.149 | 0.202 | 1 | -0.11 |
| SLC39A8 | 2.43E-25 | -0.47635 | 0.046 | 0.222 | 7.15E-21 | -2.30 | 0.778178722 | 0.004854 | 0.018 | 0.013 | 1 | 0.00 |
| UNC5D | 5.09E-11 | -0.72653 | 0.044 | 0.139 | 1.49E-06 | -2.30 | 0.742691138 | -0.00859 | 0.024 | 0.028 | 1 | -0.01 |
| IRF1 | 4.27E-15 | -0.55597 | 0.036 | 0.148 | 1.25E-10 | -2.29 | 0.233452457 | -0.05385 | 0.075 | 0.102 | 1 | -0.07 |
| ADGRL2 | 9.24E-152 | -1.50904 | 0.632 | 0.951 | 2.71E-147 | -2.27 | 2.68429E-06 | 0.27013 | 0.57 | 0.532 | 0.048926516 | 0.25 |
| SLIT2 | 3.96E-50 | -1.31493 | 0.395 | 0.679 | 1.16E-45 | -2.26 | 0.180422728 | 0.011884 | 0.134 | 0.142 | 1 | 0.01 |
| FAM78B | 9.57E-45 | -0.90003 | 0.222 | 0.551 | 2.81E-40 | -2.23 | 0.119539621 | 0.001858 | 0.009 | 0.005 | 1 | 0.00 |
| CAMK1D | 1.84E-101 | -1.37488 | 0.559 | 0.906 | 5.39E-97 | -2.23 | 0.142882925 | -0.03143 | 0.057 | 0.079 | 1 | -0.04 |
| PLEKHH2 | 1.05E-101 | -1.16339 | 0.454 | 0.869 | 3.07E-97 | -2.23 | 1.47707E-05 | 0.120351 | 0.185 | 0.167 | 0.269225414 | 0.11 |

| | | | | | | | | | | | | |
|---|---|---|---|---|---|---|---|---|---|---|---|---|
| EVA1A | 5.80E-31 | -0.67748 | 0.106 | 0.344 | 1.70E-26 | -2.20 | 1.37511E-19 | 0.472734 | 0.242 | 0.123 | 2.50641E-15 | 0.24 |
| CDK6 | 6.88E-75 | -1.1502 | 0.41 | 0.778 | 2.02E-70 | -2.18 | 1.35676E-05 | -0.22529 | 0.14 | 0.251 | 0.247296065 | -0.40 |
| LRRC4C | 6.88E-39 | -1.35437 | 0.355 | 0.567 | 2.02E-34 | -2.16 | 0.685947768 | -0.00504 | 0.003 | 0.005 | 1 | -0.01 |
| UST | 1.93E-70 | -1.13015 | 0.41 | 0.775 | 5.66E-66 | -2.14 | 0.147366156 | 0.041886 | 0.101 | 0.094 | 1 | 0.04 |
| SLC2A13 | 3.08E-48 | -0.96481 | 0.282 | 0.623 | 9.04E-44 | -2.13 | 0.043911244 | 0.045218 | 0.042 | 0.024 | 1 | 0.03 |
| SATB2 | 6.97E-55 | -0.95506 | 0.304 | 0.677 | 2.04E-50 | -2.13 | 0.162467232 | -0.02345 | 0.009 | 0.022 | 1 | -0.06 |
| XAF1 | 1.68E-11 | -0.13282 | 0.004 | 0.063 | 4.93E-07 | -2.09 | 0.11583889 | 0.011948 | 0.006 | 0.004 | 1 | 0.01 |
| FIGN | 9.52E-32 | -0.75941 | 0.135 | 0.363 | 2.80E-27 | -2.04 | 0.209019083 | -0.01018 | 0.003 | 0.012 | 1 | -0.04 |
| TMTC2 | 5.41E-48 | -1.28475 | 0.49 | 0.771 | 1.59E-43 | -2.02 | 8.2245E-13 | 0.20044 | 0.096 | 0.039 | 1.49908E-08 | 0.08 |
| PDE1B | 9.92E-17 | -0.23674 | 0.013 | 0.11 | 2.91E-12 | -2.00 | 0.811084624 | 0.003817 | 0.006 | 0.005 | 1 | 0.00 |
| VAT1L | 3.14E-10 | -0.20421 | 0.007 | 0.068 | 9.21E-06 | -1.98 | 9.79325E-86 | 0.47617 | 0.272 | 0.002 | 1.78502E-81 | 0.00 |
| EMX2OS | 5.10E-24 | -0.50403 | 0.06 | 0.23 | 1.50E-19 | -1.93 | 9.50556E-07 | -0.14838 | 0.063 | 0.155 | 0.017325788 | -0.37 |
| NAALADL2 | 1.62E-123 | -1.37037 | 0.681 | 0.958 | 4.76E-119 | -1.93 | 0.704188161 | 0.002479 | 0.006 | 0.005 | 1 | 0.00 |
| NOS1AP | 5.71E-21 | -0.50321 | 0.058 | 0.222 | 1.68E-16 | -1.93 | 0.48236013 | 0.009246 | 0.03 | 0.029 | 1 | 0.01 |
| SH3D19 | 1.09E-85 | -1.2318 | 0.543 | 0.846 | 3.21E-81 | -1.92 | 0.116757829 | -0.02978 | 0.137 | 0.169 | 1 | -0.04 |
| TCEA3 | 3.92E-22 | -0.31828 | 0.029 | 0.174 | 1.15E-17 | -1.91 | 0.046199623 | -0.03431 | 0.009 | 0.028 | 1 | -0.11 |
| EGFR | 3.36E-25 | -0.68807 | 0.093 | 0.258 | 9.87E-21 | -1.91 | 0.032281474 | -0.06233 | 0.036 | 0.069 | 1 | -0.12 |
| RASA3 | 1.59E-26 | -0.43528 | 0.055 | 0.241 | 4.66E-22 | -1.91 | 0.028183838 | -0.04132 | 0.018 | 0.044 | 1 | -0.10 |
| PPARGC1B | 1.06E-12 | -0.16864 | 0.007 | 0.079 | 3.10E-08 | -1.90 | 0.134799839 | -0.02919 | 0.018 | 0.032 | 1 | -0.05 |
| TRABD2B | 4.71E-20 | -0.55651 | 0.069 | 0.234 | 1.38E-15 | -1.89 | 0.039115769 | 0.080193 | 0.158 | 0.142 | 1 | 0.07 |
| SULF1 | 1.76E-40 | -0.8339 | 0.253 | 0.571 | 5.17E-36 | -1.88 | 0.000401397 | 0.105034 | 0.352 | 0.368 | 1 | 0.11 |
| COL4A5 | 1.83E-87 | -1.18328 | 0.554 | 0.881 | 5.39E-83 | -1.88 | 0.085457654 | -0.05117 | 0.045 | 0.076 | 1 | -0.09 |
| GEM | 2.36E-23 | -0.61147 | 0.095 | 0.292 | 6.92E-19 | -1.88 | 4.34126E-15 | 0.31386 | 0.143 | 0.043 | 7.91281E-11 | 0.09 |
| AKAP12 | 1.94E-34 | -0.86935 | 0.235 | 0.507 | 5.69E-30 | -1.88 | 0.013575175 | -0.12259 | 0.036 | 0.073 | 1 | -0.25 |
| ST6GALNAC5 | 6.48E-13 | -0.38257 | 0.02 | 0.098 | 1.90E-08 | -1.87 | 0.859320123 | 0.002307 | 0.003 | 0.002 | 1 | 0.00 |
| HOXD3 | 9.93E-36 | -0.64779 | 0.144 | 0.416 | 2.92E-31 | -1.87 | 0.414779242 | -0.01954 | 0.027 | 0.039 | 1 | -0.03 |
| RUNX1 | 3.04E-39 | -0.80198 | 0.217 | 0.504 | 8.94E-35 | -1.86 | 0.00102879 | 0.179582 | 0.236 | 0.202 | 1 | 0.15 |
| BACE2 | 7.23E-29 | -0.57215 | 0.102 | 0.329 | 2.12E-24 | -1.85 | 0.19350864 | -0.0272 | 0.012 | 0.024 | 1 | -0.05 |
| STAT1 | 8.21E-17 | -0.733 | 0.089 | 0.22 | 2.41E-12 | -1.81 | 0.773887338 | -0.00073 | 0.039 | 0.042 | 1 | 0.00 |
| TEKT1 | 1.86E-09 | -0.13128 | 0.004 | 0.055 | 5.47E-05 | -1.81 | 0.000277305 | 6.79E-06 | 0.015 | 0 | 1 | 0.00 |
| TAX1BP1 | 4.07E-96 | -1.30113 | 0.67 | 0.927 | 1.19E-91 | -1.80 | 0.466629534 | -0.07504 | 0.642 | 0.646 | 1 | -0.08 |
| TSHZ2 | 5.38E-44 | -1.05492 | 0.432 | 0.736 | 1.58E-39 | -1.80 | 0.039462963 | 0.0525 | 0.072 | 0.067 | 1 | 0.05 |

| | | | | | | | | | | | | |
|---|---|---|---|---|---|---|---|---|---|---|---|---|
| MTHFD1L | 2.02E-23 | -0.58365 | 0.086 | 0.261 | 5.93E-19 | -1.77 | 0.202120936 | -0.0379 | 0.03 | 0.048 | 1 | -0.06 |
| PDZRN3 | 3.80E-147 | -1.29715 | 0.727 | 0.976 | 1.12E-142 | -1.74 | 4.33836E-08 | -0.19193 | 0.093 | 0.212 | 0.000790752 | -0.44 |
| NDUFB9 | 1.00E-24 | -0.46321 | 0.071 | 0.261 | 2.95E-20 | -1.70 | 0.423662062 | -0.06 | 0.869 | 0.884 | 1 | -0.06 |
| CHN1 | 4.51E-33 | -0.76228 | 0.228 | 0.507 | 1.32E-28 | -1.70 | 1.98701E-10 | 0.245745 | 0.179 | 0.099 | 3.62172E-06 | 0.14 |
| LAMB1 | 3.03E-69 | -1.02264 | 0.497 | 0.813 | 8.88E-65 | -1.67 | 0.000146355 | 0.182917 | 0.296 | 0.227 | 1 | 0.14 |
| LMO7 | 9.79E-45 | -0.8655 | 0.342 | 0.653 | 2.88E-40 | -1.65 | 0.005827958 | 0.120674 | 0.185 | 0.123 | 1 | 0.08 |
| DISP1 | 2.74E-33 | -0.68818 | 0.2 | 0.48 | 8.05E-29 | -1.65 | 0.008068388 | -0.00015 | 0.054 | 0.069 | 1 | 0.00 |
| DTNB | 3.41E-44 | -0.91567 | 0.373 | 0.671 | 1.00E-39 | -1.65 | 0.345942417 | -0.01895 | 0.119 | 0.138 | 1 | -0.02 |
| TTC39C | 2.82E-31 | -0.66059 | 0.18 | 0.447 | 8.29E-27 | -1.64 | 7.37895E-05 | 0.074869 | 0.069 | 0.021 | 1 | 0.02 |
| PLXDC2 | 4.40E-46 | -0.97994 | 0.441 | 0.738 | 1.29E-41 | -1.64 | 0.088564141 | -0.05316 | 0.146 | 0.187 | 1 | -0.07 |
| TRAF3 | 6.31E-29 | -0.65736 | 0.148 | 0.365 | 1.85E-24 | -1.62 | 0.256144565 | -0.0315 | 0.066 | 0.089 | 1 | -0.04 |
| PBX1 | 5.59E-54 | -0.96289 | 0.474 | 0.788 | 1.64E-49 | -1.60 | 7.74355E-11 | 0.268492 | 0.567 | 0.55 | 1.41142E-06 | 0.26 |
| ADAM22 | 5.06E-74 | -0.99128 | 0.546 | 0.878 | 1.49E-69 | -1.59 | 4.55134E-08 | 0.223786 | 0.146 | 0.092 | 0.000829573 | 0.14 |
| ZFPM2 | 1.34E-21 | -0.79232 | 0.182 | 0.366 | 3.94E-17 | -1.59 | 0.505523379 | -0.00385 | 0.003 | 0.006 | 1 | -0.01 |
| SNX8 | 2.89E-28 | -0.52401 | 0.113 | 0.343 | 8.48E-24 | -1.59 | 0.00087526 | -0.05396 | 0.036 | 0.076 | 1 | -0.11 |
| LRFN5 | 1.13E-23 | -0.67966 | 0.166 | 0.388 | 3.31E-19 | -1.59 | 0.280556787 | -0.0122 | 0.003 | 0.011 | 1 | -0.04 |
| RASSF8 | 2.70E-54 | -0.95496 | 0.481 | 0.79 | 7.94E-50 | -1.57 | 0.008907442 | 0.11058 | 0.209 | 0.172 | 1 | 0.09 |
| MRPS22 | 2.18E-23 | -0.52284 | 0.1 | 0.298 | 6.41E-19 | -1.56 | 0.008764623 | -0.03756 | 0.167 | 0.215 | 1 | -0.05 |
| TSC22D2 | 5.24E-48 | -0.99506 | 0.47 | 0.729 | 1.54E-43 | -1.54 | 4.35818E-09 | -0.27713 | 0.254 | 0.423 | 7.94365E-05 | -0.46 |
| EPS8 | 1.01E-41 | -0.86271 | 0.35 | 0.623 | 2.97E-37 | -1.54 | 1.56125E-05 | 0.089383 | 0.051 | 0.032 | 0.284569476 | 0.06 |
| IGDCC4 | 2.78E-31 | -0.56304 | 0.151 | 0.411 | 8.16E-27 | -1.53 | 0.000110816 | -0.13174 | 0.104 | 0.188 | 1 | -0.24 |
| PPFIBP1 | 6.78E-50 | -0.89437 | 0.362 | 0.619 | 1.99E-45 | -1.53 | 5.33382E-05 | -0.1962 | 0.149 | 0.245 | 0.97219534 | -0.32 |
| IFI35 | 2.04E-09 | -0.11453 | 0.004 | 0.053 | 6.00E-05 | -1.52 | 0.00859574 | -0.05238 | 0.036 | 0.073 | 1 | -0.11 |
| NCAM2 | 1.27E-17 | -1.02799 | 0.383 | 0.562 | 3.73E-13 | -1.51 | 0.001502199 | 0.070596 | 0.021 | 0.005 | 1 | 0.02 |
| FAM189A2 | 2.63E-17 | -0.36472 | 0.04 | 0.165 | 7.71E-13 | -1.50 | 0.003766627 | 0.017081 | 0.015 | 0.002 | 1 | 0.00 |
| FSTL5 | 1.00E-123 | -2.45655 | 0.038 | 0.537 | 2.94E-119 | -34.71 | 0.326373758 | 0.006738 | 0.003 | 0.001 | 1 | 0.00 |

**Supplementary Table 12: Mouse Specific Dermis Enriched Genes. U score = avg_log2FC x pct.2/pct.1. U score < -1.5 for mouse, > -0.5 for human.**

| | Human | | | | | | Mouse (Qu et al., 2022) | | | | | |
|---|---|---|---|---|---|---|---|---|---|---|---|---|
| | p_val | avg_log2FC | pct.1 | pct.2 | p_val_adj | UScore | p_val | avg_log2FC | pct.1 | pct.2 | p_val_adj | UScore |
| Cthrc1 | 0.381502 | 0.041375 | 0.056 | 0.043 | 1 | 0.03 | 1.18E-72 | -1.16185 | 0.054 | 0.533 | 2.14E-68 | -11.47 |
| Erg | 0.706292 | 0.004069 | 0.031 | 0.03 | 1 | 0.00 | 6.10E-18 | -0.25906 | 0.003 | 0.132 | 1.11E-13 | -11.40 |
| Lox | 0.001358 | 0.096364 | 0.111 | 0.109 | 1 | 0.09 | 7.09E-62 | -1.18174 | 0.054 | 0.478 | 1.29E-57 | -10.46 |
| Ch25h | 0.231494 | -0.02395 | 0.005 | 0.014 | 1 | -0.07 | 1.30E-33 | -0.52378 | 0.015 | 0.262 | 2.37E-29 | -9.15 |
| Cldn10 | 0.824119 | -0.00237 | 0.002 | 0.003 | 1 | 0.00 | 7.68E-17 | -0.31067 | 0.006 | 0.134 | 1.40E-12 | -6.94 |
| Fibin | 0.022857 | -0.02622 | 0.002 | 0.014 | 1 | -0.18 | 2.01E-22 | -0.52541 | 0.015 | 0.195 | 3.66E-18 | -6.83 |
| Nfib | 8.54E-16 | -0.29531 | 0.206 | 0.258 | 2.51E-11 | -0.37 | 1.16E-90 | -1.55621 | 0.161 | 0.697 | 2.12E-86 | -6.74 |
| Ogn | 0.00033 | -0.10176 | 0.089 | 0.15 | 1 | -0.17 | 3.57E-28 | -0.57413 | 0.021 | 0.245 | 6.50E-24 | -6.70 |
| Rspo1 | 0.095817 | 0.051678 | 0.071 | 0.067 | 1 | 0.05 | 6.53E-75 | -1.18282 | 0.125 | 0.637 | 1.19E-70 | -6.03 |
| Nnmt | 0.052317 | -0.02248 | 0.007 | 0.022 | 1 | -0.07 | 2.73E-12 | -0.18886 | 0.003 | 0.094 | 4.98E-08 | -5.92 |
| Cxcl12 | 1.27E-06 | -0.06623 | 0.046 | 0.102 | 0.037322 | -0.15 | 3.42E-54 | -1.01982 | 0.087 | 0.497 | 6.24E-50 | -5.83 |
| Kitl | 0.086129 | -0.02582 | 0.026 | 0.036 | 1 | -0.04 | 1.97E-39 | -0.9992 | 0.072 | 0.399 | 3.59E-35 | -5.54 |
| Islr | 0.096531 | 0.015312 | 0.033 | 0.039 | 1 | 0.02 | 2.61E-33 | -0.55072 | 0.03 | 0.296 | 4.76E-29 | -5.43 |
| Efemp1 | 0.684624 | 0.01613 | 0.026 | 0.02 | 1 | 0.01 | 7.05E-17 | -0.29135 | 0.009 | 0.142 | 1.29E-12 | -4.60 |
| Fkbp11 | 0.37768 | 0.013434 | 0.049 | 0.054 | 1 | 0.01 | 2.84E-49 | -0.93215 | 0.107 | 0.509 | 5.18E-45 | -4.43 |
| Dpt | 0.007413 | 0.077519 | 0.137 | 0.137 | 1 | 0.08 | 2.42E-21 | -0.49629 | 0.024 | 0.209 | 4.40E-17 | -4.32 |
| Tmem150c | 0.106817 | -0.05659 | 0.035 | 0.056 | 1 | -0.09 | 3.41E-28 | -0.5013 | 0.033 | 0.272 | 6.21E-24 | -4.13 |
| Irx1 | 8.62E-11 | -0.20772 | 0.08 | 0.19 | 2.53E-06 | -0.49 | 6.19E-76 | -1.23002 | 0.218 | 0.721 | 1.13E-71 | -4.07 |
| Ptn | 3.02E-31 | 0.12493 | 0.133 | 0.011 | 8.88E-27 | 0.01 | 6.36E-48 | -1.14397 | 0.161 | 0.562 | 1.16E-43 | -3.99 |
| Vcam1 | 0.000761 | -0.07275 | 0.015 | 0.047 | 1 | -0.23 | 2.23E-107 | -1.45203 | 0.322 | 0.854 | 4.07E-103 | -3.85 |
| Enho | 0.00048 | 0.051247 | 0.018 | 0.003 | 1 | 0.01 | 3.13E-15 | -0.24549 | 0.009 | 0.132 | 5.70E-11 | -3.60 |
| Tubb6 | 0.774617 | 0.022392 | 0.29 | 0.274 | 1 | 0.02 | 2.59E-40 | -0.94712 | 0.134 | 0.497 | 4.73E-36 | -3.51 |
| Nrn1 | 9.37E-05 | -0.0791 | 0.046 | 0.093 | 1 | -0.16 | 1.28E-35 | -0.8249 | 0.101 | 0.43 | 2.34E-31 | -3.51 |
| 2810417H13Rik | 0.250405 | -0.03293 | 0.04 | 0.057 | 1 | -0.05 | 4.65E-52 | -1.13152 | 0.209 | 0.638 | 8.47E-48 | -3.45 |
| Pcolce | 0.651555 | 0.013428 | 0.064 | 0.064 | 1 | 0.01 | 1.16E-57 | -1.02835 | 0.197 | 0.641 | 2.11E-53 | -3.35 |
| Sdpr | 0.051587 | -0.01664 | 0.002 | 0.012 | 1 | -0.10 | 2.08E-09 | -0.13461 | 0.003 | 0.073 | 3.79E-05 | -3.28 |
| Gins2 | 0.248399 | -0.01295 | 0.044 | 0.058 | 1 | -0.02 | 9.76E-38 | -0.73192 | 0.099 | 0.438 | 1.78E-33 | -3.24 |

| | | | | | | | | | | | | |
|---|---|---|---|---|---|---|---|---|---|---|---|---|
| Mest | 1.84E-17 | 0.3929 | 0.712 | 0.743 | 5.4E-13 | 0.41 | 3.97E-104 | -1.70787 | 0.469 | 0.887 | 7.24E-100 | -3.23 |
| Lpl | 0.000304 | 0.023433 | 0.026 | 0.006 | 1 | 0.01 | 5.95E-19 | -0.31803 | 0.018 | 0.178 | 1.08E-14 | -3.14 |
| Sp5 | 0.052647 | 0.055299 | 0.047 | 0.037 | 1 | 0.04 | 1.35E-35 | -0.67048 | 0.09 | 0.414 | 2.47E-31 | -3.08 |
| Birc5 | 0.257452 | -0.03246 | 0.024 | 0.034 | 1 | -0.05 | 3.88E-26 | -1.09356 | 0.158 | 0.439 | 7.07E-22 | -3.04 |
| Cdkn3 | 0.316463 | -0.02742 | 0.027 | 0.04 | 1 | -0.04 | 1.51E-18 | -0.41694 | 0.027 | 0.196 | 2.74E-14 | -3.03 |
| Htra3 | 0.043627 | -0.02294 | 0.005 | 0.018 | 1 | -0.08 | 4.14E-11 | -0.18603 | 0.006 | 0.095 | 7.55E-07 | -2.95 |
| Cenpa | 0.016653 | -0.06324 | 0.022 | 0.048 | 1 | -0.14 | 9.57E-22 | -1.01433 | 0.128 | 0.371 | 1.74E-17 | -2.94 |
| Eif4ebp1 | 0.120934 | 0.04562 | 0.042 | 0.033 | 1 | 0.04 | 9.70E-82 | -1.14778 | 0.331 | 0.824 | 1.77E-77 | -2.86 |
| Idi1 | 0.119173 | 0.006244 | 0.089 | 0.101 | 1 | 0.01 | 3.13E-30 | -0.7079 | 0.101 | 0.397 | 5.70E-26 | -2.78 |
| Zfp385b | 0.238101 | -0.00832 | 0.002 | 0.008 | 1 | -0.03 | 3.79E-26 | -0.44449 | 0.045 | 0.28 | 6.91E-22 | -2.77 |
| P4ha2 | 1.12E-05 | 0.198145 | 0.189 | 0.118 | 0.329369 | 0.12 | 4.51E-46 | -0.85188 | 0.179 | 0.579 | 8.22E-42 | -2.76 |
| Sfrp5 | 0.613431 | 0.000511 | 0.004 | 0.002 | 1 | 0.00 | 3.39E-27 | -0.83443 | 0.128 | 0.412 | 6.18E-23 | -2.69 |
| Tmem97 | 0.02336 | 0.061751 | 0.04 | 0.029 | 1 | 0.04 | 6.02E-36 | -0.7053 | 0.119 | 0.452 | 1.10E-31 | -2.68 |
| Mki67 | 0.144057 | -0.05453 | 0.055 | 0.065 | 1 | -0.06 | 3.89E-18 | -0.80238 | 0.09 | 0.3 | 7.09E-14 | -2.67 |
| Irx3 | 0.084706 | -0.02801 | 0.011 | 0.026 | 1 | -0.07 | 7.07E-44 | -0.85465 | 0.194 | 0.586 | 1.29E-39 | -2.58 |
| Crip1 | 0.063538 | 0.028878 | 0.06 | 0.038 | 1 | 0.02 | 7.75E-254 | -1.73756 | 0.722 | 0.996 | 1.41E-249 | -2.40 |
| Adam33 | 3.49E-06 | 0.168485 | 0.095 | 0.068 | 0.102337 | 0.12 | 4.76E-27 | -0.62944 | 0.099 | 0.373 | 8.68E-23 | -2.37 |
| Hmgn3 | 1.87E-07 | 0.234242 | 0.69 | 0.685 | 0.005485 | 0.23 | 6.06E-114 | -1.45043 | 0.57 | 0.928 | 1.11E-109 | -2.36 |
| Cdca3 | 0.125848 | -0.02295 | 0.02 | 0.034 | 1 | -0.04 | 4.85E-20 | -0.70106 | 0.09 | 0.297 | 8.83E-16 | -2.31 |
| Osr1 | 0.032375 | -0.05405 | 0.015 | 0.034 | 1 | -0.12 | 4.22E-60 | -0.87751 | 0.284 | 0.746 | 7.69E-56 | -2.31 |
| Fam114a1 | 3.46E-11 | 0.225977 | 0.386 | 0.382 | 1.02E-06 | 0.22 | 2.23E-33 | -0.66721 | 0.134 | 0.462 | 4.07E-29 | -2.30 |
| Ccnb2 | 0.027655 | -0.04349 | 0.015 | 0.035 | 1 | -0.10 | 3.79E-20 | -0.6794 | 0.096 | 0.325 | 6.91E-16 | -2.30 |
| Sulf2 | 1.32E-07 | 0.267122 | 0.168 | 0.095 | 0.00388 | 0.15 | 2.39E-32 | -0.61137 | 0.116 | 0.434 | 4.35E-28 | -2.29 |
| Gm22 | 0.509142 | 0.005685 | 0.005 | 0.006 | 1 | 0.01 | 3.02E-14 | -0.24281 | 0.015 | 0.14 | 5.51E-10 | -2.27 |
| Cks2 | 0.609013 | 5.03E-05 | 0.015 | 0.018 | 1 | 0.00 | 4.22E-26 | -1.05599 | 0.251 | 0.536 | 7.69E-22 | -2.26 |
| Comt | 8.96E-09 | -0.19952 | 0.239 | 0.367 | 0.000263 | -0.31 | 8.44E-38 | -0.68861 | 0.167 | 0.53 | 1.54E-33 | -2.19 |
| Maf | 0.011347 | -0.08426 | 0.118 | 0.167 | 1 | -0.12 | 8.15E-26 | -0.46408 | 0.069 | 0.321 | 1.49E-21 | -2.16 |
| Medag | 0.136196 | 1.28E-15 | 0.002 | 0 | 1 | 0.00 | 1.42E-07 | -0.1069 | 0.003 | 0.06 | 0.002586 | -2.14 |
| Tacc3 | 0.010177 | -0.11282 | 0.036 | 0.069 | 1 | -0.22 | 4.58E-18 | -0.61207 | 0.078 | 0.271 | 8.36E-14 | -2.13 |
| Fkbp10 | 0.004931 | 0.022187 | 0.239 | 0.265 | 1 | 0.02 | 1.57E-31 | -0.64156 | 0.14 | 0.461 | 2.86E-27 | -2.11 |
| Grem1 | 0.002228 | -0.04527 | 0.022 | 0.048 | 1 | -0.10 | 5.56E-13 | -0.36849 | 0.027 | 0.153 | 1.01E-08 | -2.09 |
| Copz2 | 0.039826 | -0.01769 | 0.027 | 0.044 | 1 | -0.03 | 2.99E-42 | -0.83084 | 0.257 | 0.641 | 5.45E-38 | -2.07 |

| | | | | | | | | | | | | |
|---|---|---|---|---|---|---|---|---|---|---|---|---|
| Gas1 | 0.013795 | 0.050553 | 0.117 | 0.131 | 1 | 0.06 | 6.20E-111 | -1.18453 | 0.546 | 0.955 | 1.13E-106 | -2.07 |
| Cenpf | 0.00072 | -0.18853 | 0.062 | 0.115 | 1 | -0.35 | 5.96E-10 | -0.76821 | 0.081 | 0.216 | 1.09E-05 | -2.05 |
| Selm | 0.422765 | -0.00661 | 0.016 | 0.023 | 1 | -0.01 | 1.77E-56 | -1.00312 | 0.397 | 0.796 | 3.22E-52 | -2.01 |
| Pcsk6 | 8.99E-08 | -0.18353 | 0.046 | 0.112 | 0.002641 | -0.45 | 3.15E-21 | -0.40645 | 0.054 | 0.266 | 5.73E-17 | -2.00 |
| Ccna2 | 0.09408 | -0.04653 | 0.035 | 0.057 | 1 | -0.08 | 3.79E-16 | -0.63442 | 0.093 | 0.288 | 6.91E-12 | -1.96 |
| En1 | 0.029047 | -0.04038 | 0.029 | 0.053 | 1 | -0.07 | 3.82E-26 | -0.47913 | 0.087 | 0.349 | 6.97E-22 | -1.92 |
| Tpx2 | 0.002051 | -0.16809 | 0.046 | 0.083 | 1 | -0.30 | 2.62E-13 | -0.60293 | 0.075 | 0.239 | 4.78E-09 | -1.92 |
| Ifitm3 | 0.736595 | -0.00966 | 0.027 | 0.033 | 1 | -0.01 | 7.75E-33 | -0.61037 | 0.137 | 0.431 | 1.41E-28 | -1.92 |
| Dusp10 | 0.379044 | -0.03034 | 0.035 | 0.048 | 1 | -0.04 | 5.98E-22 | -0.47205 | 0.075 | 0.305 | 1.09E-17 | -1.92 |
| Dnajc15 | 0.0002 | -0.15562 | 0.302 | 0.396 | 1 | -0.20 | 1.78E-44 | -0.87784 | 0.313 | 0.679 | 3.24E-40 | -1.90 |
| Hmmr | 0.319033 | -0.02574 | 0.007 | 0.014 | 1 | -0.05 | 4.94E-11 | -0.63918 | 0.069 | 0.205 | 9.00E-07 | -1.90 |
| Fxyd1 | 0.427844 | 0.00583 | 0.013 | 0.015 | 1 | 0.01 | 1.32E-18 | -0.38759 | 0.048 | 0.234 | 2.40E-14 | -1.89 |
| Rcn3 | 6.88E-06 | -0.05626 | 0.077 | 0.131 | 0.201934 | -0.10 | 8.57E-68 | -1.15635 | 0.528 | 0.855 | 1.56E-63 | -1.87 |
| Dcn | 9.77E-07 | -0.26212 | 0.397 | 0.523 | 0.028682 | -0.35 | 2.13E-238 | -1.56224 | 0.839 | 0.995 | 3.88E-234 | -1.85 |
| Pcsk9 | 1.23E-05 | -0.09134 | 0.027 | 0.074 | 0.361138 | -0.25 | 1.39E-12 | -0.21855 | 0.015 | 0.127 | 2.53E-08 | -1.85 |
| Gap43 | 0.498744 | 0.027605 | 0.038 | 0.037 | 1 | 0.03 | 9.23E-27 | -0.6938 | 0.176 | 0.469 | 1.68E-22 | -1.85 |
| Tubb4b | 0.575717 | -0.00073 | 0.056 | 0.049 | 1 | 0.00 | 5.12E-30 | -1.11566 | 0.409 | 0.669 | 9.34E-26 | -1.82 |
| Fdps | 0.002889 | -0.0446 | 0.189 | 0.24 | 1 | -0.06 | 1.11E-26 | -0.71955 | 0.194 | 0.489 | 2.02E-22 | -1.81 |
| Adk | 0.000628 | -0.17734 | 0.792 | 0.837 | 1 | -0.19 | 2.74E-34 | -0.58537 | 0.164 | 0.508 | 4.99E-30 | -1.81 |
| Igf2 | 0.056968 | 0.008575 | 0.02 | 0.009 | 1 | 0.00 | 5.58E-25 | -1.25536 | 0.513 | 0.739 | 1.02E-20 | -1.81 |
| Cdc20 | 0.512382 | -0.00672 | 0.005 | 0.011 | 1 | -0.01 | 1.89E-11 | -0.7036 | 0.096 | 0.246 | 3.44E-07 | -1.80 |
| Cd34 | 0.002439 | 0.03026 | 0.299 | 0.337 | 1 | 0.03 | 4.01E-17 | -0.3822 | 0.048 | 0.226 | 7.30E-13 | -1.80 |
| Kdelr2 | 0.003118 | -0.0219 | 0.082 | 0.114 | 1 | -0.03 | 9.85E-47 | -1.05666 | 0.484 | 0.822 | 1.80E-42 | -1.79 |
| Hic1 | 0.006065 | -0.04009 | 0.004 | 0.021 | 1 | -0.21 | 5.03E-28 | -0.56487 | 0.14 | 0.44 | 9.17E-24 | -1.78 |
| Tram1 | 0.045633 | -0.03612 | 0.231 | 0.271 | 1 | -0.04 | 3.07E-45 | -0.81333 | 0.331 | 0.717 | 5.60E-41 | -1.76 |
| Il34 | 0.103868 | -0.05591 | 0.038 | 0.06 | 1 | -0.09 | 4.74E-06 | -0.10434 | 0.003 | 0.05 | 0.086448 | -1.74 |
| Srm | 0.494193 | 0.019046 | 0.055 | 0.043 | 1 | 0.01 | 1.41E-42 | -0.90718 | 0.382 | 0.726 | 2.57E-38 | -1.72 |
| Insig1 | 0.313819 | -0.02509 | 0.04 | 0.056 | 1 | -0.04 | 4.10E-21 | -0.50911 | 0.101 | 0.34 | 7.47E-17 | -1.71 |
| Emx2 | 3.48E-06 | -0.10681 | 0.022 | 0.072 | 0.102067 | -0.35 | 5.14E-19 | -0.33001 | 0.045 | 0.229 | 9.36E-15 | -1.68 |
| Tgfbi | 2.23E-06 | 0.118417 | 0.129 | 0.15 | 0.065398 | 0.14 | 1.34E-13 | -0.25961 | 0.024 | 0.155 | 2.43E-09 | -1.68 |
| Kdelr3 | 0.006325 | -0.00586 | 0.044 | 0.063 | 1 | -0.01 | 3.27E-41 | -0.82902 | 0.349 | 0.704 | 5.96E-37 | -1.67 |
| Cdca8 | 0.053267 | -0.06303 | 0.022 | 0.043 | 1 | -0.12 | 9.67E-14 | -0.59464 | 0.099 | 0.275 | 1.76E-09 | -1.65 |

| | | | | | | | | | | | | |
|---|---|---|---|---|---|---|---|---|---|---|---|---|
| Ckap2l | 0.244296 | -0.03568 | 0.026 | 0.04 | 1 | -0.05 | 1.25E-11 | -0.36242 | 0.036 | 0.163 | 2.28E-07 | -1.64 |
| Ttc39a | 0.217855 | -0.01659 | 0.009 | 0.018 | 1 | -0.03 | 3.03E-06 | -0.09361 | 0.003 | 0.051 | 0.055255 | -1.59 |
| Tuba1b | 0.011803 | -0.09067 | 0.082 | 0.126 | 1 | -0.14 | 2.59E-144 | -1.34074 | 0.836 | 0.984 | 4.72E-140 | -1.58 |
| Cpz | 2.73E-05 | 0.121042 | 0.055 | 0.017 | 0.80163 | 0.04 | 3.54E-12 | -0.23669 | 0.021 | 0.138 | 6.45E-08 | -1.56 |
| Col1a1 | 1.62E-15 | -0.2836 | 0.761 | 0.9 | 4.77E-11 | -0.34 | 4.82E-203 | -1.485 | 0.97 | 0.998 | 8.79E-199 | -1.53 |
| Ung | 0.43519 | 0.023273 | 0.056 | 0.043 | 1 | 0.02 | 3.06E-16 | -0.34504 | 0.051 | 0.224 | 5.58E-12 | -1.52 |
| Nucb2 | 0.05946 | -0.038 | 0.128 | 0.157 | 1 | -0.05 | 8.71E-19 | -0.35177 | 0.06 | 0.257 | 1.59E-14 | -1.51 |
| Msmo1 | 0.009338 | 0.117408 | 0.142 | 0.113 | 1 | 0.09 | 1.53E-17 | -0.50773 | 0.11 | 0.325 | 2.79E-13 | -1.50 |

**Supplementary Table 13: Placode and IFE enriched genes identified in either the mouse (Qu et al., 2022) or human where no orthologue was found in the other species.**

| | Human | | | | | | Mouse (Qu et al., 2022) | | | | | |
|---|---|---|---|---|---|---|---|---|---|---|---|---|
| | p_val | avg_log2FC | pct.1 | pct.2 | p_val_adj | UScore | p_val | avg_log2FC | pct.1 | pct.2 | p_val_adj | UScore |
| **Placode Enriched** | | | | | | | | | | | | |
| AC092422.1 | 2.28E-27 | 1.45 | 0.25 | 0.012 | 6.70E-23 | 30.05 | #N/A | #N/A | #N/A | #N/A | #N/A | #N/A |
| AL392172.2 | 3.26E-12 | 0.30 | 0.10 | 0.001 | 9.57E-08 | 28.89 | #N/A | #N/A | #N/A | #N/A | #N/A | #N/A |
| DEC01 | 4.18E-54 | 2.13 | 0.54 | 0.048 | 1.23E-49 | 23.98 | #N/A | #N/A | #N/A | #N/A | #N/A | #N/A |
| AC055854.1 | 1.31E-31 | 0.95 | 0.31 | 0.014 | 3.85E-27 | 20.80 | #N/A | #N/A | #N/A | #N/A | #N/A | #N/A |
| KIF12 | 2.91E-27 | 0.51 | 0.24 | 0.007 | 8.54E-23 | 17.56 | #N/A | #N/A | #N/A | #N/A | #N/A | #N/A |
| KCNMA1-AS1 | 5.84E-13 | 0.32 | 0.11 | 0.002 | 1.71E-08 | 17.05 | #N/A | #N/A | #N/A | #N/A | #N/A | #N/A |
| AC078923.1 | 1.40E-26 | 1.33 | 0.32 | 0.029 | 4.12E-22 | 14.58 | #N/A | #N/A | #N/A | #N/A | #N/A | #N/A |
| PEBP4 | 2.97E-30 | 1.37 | 0.34 | 0.033 | 8.72E-26 | 14.25 | #N/A | #N/A | #N/A | #N/A | #N/A | #N/A |
| FAM177B | 8.68E-08 | 0.21 | 0.06 | 0.001 | 0.002549 | 13.48 | #N/A | #N/A | #N/A | #N/A | #N/A | #N/A |
| AC079382.2 | 8.68E-08 | 0.19 | 0.06 | 0.001 | 0.002548 | 11.86 | #N/A | #N/A | #N/A | #N/A | #N/A | #N/A |
| AP002762.2 | 1.77E-15 | 0.54 | 0.15 | 0.007 | 5.20E-11 | 11.80 | #N/A | #N/A | #N/A | #N/A | #N/A | #N/A |
| AC022915.2 | 2.92E-20 | 0.57 | 0.21 | 0.012 | 8.56E-16 | 10.05 | #N/A | #N/A | #N/A | #N/A | #N/A | #N/A |
| LINC00378 | 1.85E-13 | 1.07 | 0.19 | 0.023 | 5.44E-09 | 8.59 | #N/A | #N/A | #N/A | #N/A | #N/A | #N/A |
| LINC01239 | 6.79E-05 | 0.20 | 0.04 | 0.001 | 1 | 7.61 | #N/A | #N/A | #N/A | #N/A | #N/A | #N/A |
| AL583785.1 | 1.16E-14 | 1.31 | 0.24 | 0.041 | 3.42E-10 | 7.54 | #N/A | #N/A | #N/A | #N/A | #N/A | #N/A |
| MME-AS1 | 6.02E-13 | 0.53 | 0.15 | 0.013 | 1.77E-08 | 6.22 | #N/A | #N/A | #N/A | #N/A | #N/A | #N/A |
| MUC19 | 9.96E-10 | 0.28 | 0.10 | 0.005 | 2.92E-05 | 5.62 | #N/A | #N/A | #N/A | #N/A | #N/A | #N/A |
| PTPRN2 | 2.51E-22 | 1.25 | 0.37 | 0.089 | 7.35E-18 | 5.18 | #N/A | #N/A | #N/A | #N/A | #N/A | #N/A |
| LINC00882 | 1.66E-17 | 1.98 | 0.24 | 0.096 | 4.87E-13 | 4.99 | #N/A | #N/A | #N/A | #N/A | #N/A | #N/A |
| AC105094.2 | 2.96E-09 | 0.33 | 0.10 | 0.007 | 8.68E-05 | 4.85 | #N/A | #N/A | #N/A | #N/A | #N/A | #N/A |
| AC011287.1 | 8.91E-07 | 1.26 | 0.15 | 0.043 | 0.026168 | 4.29 | #N/A | #N/A | #N/A | #N/A | #N/A | #N/A |
| AL449403.2 | 5.23E-07 | 0.13 | 0.06 | 0.002 | 0.015357 | 4.06 | #N/A | #N/A | #N/A | #N/A | #N/A | #N/A |
| MYCNOS | 0.000137 | 0.10 | 0.04 | 0.001 | 1 | 3.88 | #N/A | #N/A | #N/A | #N/A | #N/A | #N/A |
| AC009313.1 | 1.86E-09 | 0.28 | 0.11 | 0.008 | 5.45E-05 | 3.79 | #N/A | #N/A | #N/A | #N/A | #N/A | #N/A |
| ATP2B2 | 3.56E-11 | 0.52 | 0.17 | 0.024 | 1.05E-06 | 3.76 | #N/A | #N/A | #N/A | #N/A | #N/A | #N/A |
| AC068305.2 | 1.00E-15 | 0.96 | 0.37 | 0.096 | 2.95E-11 | 3.69 | #N/A | #N/A | #N/A | #N/A | #N/A | #N/A |

| | | | | | | | | | | | | |
|---|---|---|---|---|---|---|---|---|---|---|---|---|
| KIAA1549L | 1.29E-13 | 0.62 | 0.24 | 0.041 | 3.78E-09 | 3.67 | #N/A | #N/A | #N/A | #N/A | #N/A | #N/A |
| LINC01919 | 0.00072 | 0.11 | 0.03 | 0.001 | 1 | 3.50 | #N/A | #N/A | #N/A | #N/A | #N/A | #N/A |
| AC092110.1 | 0.000883 | 0.11 | 0.03 | 0.001 | 1 | 3.44 | #N/A | #N/A | #N/A | #N/A | #N/A | #N/A |
| AC034111.2 | 0.000553 | 0.18 | 0.04 | 0.002 | 1 | 3.41 | #N/A | #N/A | #N/A | #N/A | #N/A | #N/A |
| AC005237.1 | 1.65E-13 | 0.58 | 0.25 | 0.050 | 4.84E-09 | 2.87 | #N/A | #N/A | #N/A | #N/A | #N/A | #N/A |
| LINC00926 | 5.88E-12 | 0.26 | 0.15 | 0.014 | 1.73E-07 | 2.85 | #N/A | #N/A | #N/A | #N/A | #N/A | #N/A |
| NBAT1 | 1.32E-29 | 1.12 | 0.67 | 0.264 | 3.87E-25 | 2.83 | #N/A | #N/A | #N/A | #N/A | #N/A | #N/A |
| AL449983.1 | 2.19E-14 | 0.53 | 0.27 | 0.053 | 6.42E-10 | 2.69 | #N/A | #N/A | #N/A | #N/A | #N/A | #N/A |
| CASC15 | 4.93E-118 | 2.41 | 0.99 | 0.897 | 1.45E-113 | 2.67 | #N/A | #N/A | #N/A | #N/A | #N/A | #N/A |
| AC090004.2 | 0.000878 | 0.08 | 0.03 | 0.001 | 1 | 2.65 | #N/A | #N/A | #N/A | #N/A | #N/A | #N/A |
| HTN1 | 1.12E-05 | 0.14 | 0.06 | 0.003 | 0.327819 | 2.61 | #N/A | #N/A | #N/A | #N/A | #N/A | #N/A |
| AL161729.2 | 3.30E-06 | 0.09 | 0.06 | 0.002 | 0.096888 | 2.57 | #N/A | #N/A | #N/A | #N/A | #N/A | #N/A |
| AC078820.2 | 7.03E-08 | 0.30 | 0.10 | 0.012 | 0.002064 | 2.57 | #N/A | #N/A | #N/A | #N/A | #N/A | #N/A |
| AL096865.1 | 0.00012 | 0.07 | 0.04 | 0.001 | 1 | 2.57 | #N/A | #N/A | #N/A | #N/A | #N/A | #N/A |
| AC009063.2 | 5.81E-05 | 0.11 | 0.05 | 0.002 | 1 | 2.50 | #N/A | #N/A | #N/A | #N/A | #N/A | #N/A |
| NAV2-AS3 | 1.30E-05 | 0.10 | 0.05 | 0.002 | 0.380576 | 2.47 | #N/A | #N/A | #N/A | #N/A | #N/A | #N/A |
| PRR27 | 1.63E-05 | 0.26 | 0.07 | 0.008 | 0.480005 | 2.24 | #N/A | #N/A | #N/A | #N/A | #N/A | #N/A |
| SEMA6A-AS1 | 1.74E-25 | 1.21 | 0.73 | 0.406 | 5.10E-21 | 2.17 | #N/A | #N/A | #N/A | #N/A | #N/A | #N/A |
| AF130359.1 | 0.004999 | 0.09 | 0.03 | 0.001 | 1 | 2.13 | #N/A | #N/A | #N/A | #N/A | #N/A | #N/A |
| AL589693.1 | 4.54E-16 | 1.49 | 0.71 | 0.521 | 1.33E-11 | 2.03 | #N/A | #N/A | #N/A | #N/A | #N/A | #N/A |
| AC100800.1 | 1.00E-14 | 0.53 | 0.35 | 0.093 | 2.94E-10 | 2.00 | #N/A | #N/A | #N/A | #N/A | #N/A | #N/A |
| MAP1LC3B2 | 2.30E-06 | 0.21 | 0.08 | 0.009 | 0.067433 | 1.98 | #N/A | #N/A | #N/A | #N/A | #N/A | #N/A |
| AC090673.1 | 2.43E-08 | 0.34 | 0.15 | 0.025 | 0.000713 | 1.97 | #N/A | #N/A | #N/A | #N/A | #N/A | #N/A |
| SPACA9 | 0.000102 | 0.08 | 0.05 | 0.002 | 1 | 1.91 | #N/A | #N/A | #N/A | #N/A | #N/A | #N/A |
| LINC01006 | 1.04E-08 | 0.74 | 0.31 | 0.119 | 0.000305 | 1.91 | #N/A | #N/A | #N/A | #N/A | #N/A | #N/A |
| PLEKHG4B | 2.21E-16 | 0.72 | 0.52 | 0.198 | 6.48E-12 | 1.91 | #N/A | #N/A | #N/A | #N/A | #N/A | #N/A |
| AC108879.1 | 0.004991 | 0.07 | 0.03 | 0.001 | 1 | 1.81 | #N/A | #N/A | #N/A | #N/A | #N/A | #N/A |
| CCDC175 | 6.84E-05 | 0.10 | 0.05 | 0.003 | 1 | 1.77 | #N/A | #N/A | #N/A | #N/A | #N/A | #N/A |
| AC079465.1 | 3.42E-10 | 0.69 | 0.31 | 0.121 | 1.00E-05 | 1.77 | #N/A | #N/A | #N/A | #N/A | #N/A | #N/A |
| AL139294.1 | 6.38E-10 | 0.29 | 0.18 | 0.030 | 1.87E-05 | 1.75 | #N/A | #N/A | #N/A | #N/A | #N/A | #N/A |
| GATA3-AS1 | 7.10E-05 | 0.10 | 0.05 | 0.003 | 1 | 1.73 | #N/A | #N/A | #N/A | #N/A | #N/A | #N/A |
| AC023282.1 | 5.31E-11 | 0.77 | 0.47 | 0.214 | 1.56E-06 | 1.70 | #N/A | #N/A | #N/A | #N/A | #N/A | #N/A |

| | | | | | | | | | | | | |
|---|---|---|---|---|---|---|---|---|---|---|---|---|
| FBN3 | 4.73E-18 | 0.86 | 0.63 | 0.320 | 1.39E-13 | 1.70 | #N/A | #N/A | #N/A | #N/A | #N/A | #N/A |
| AC126177.3 | 0.00499 | 0.07 | 0.03 | 0.001 | 1 | 1.65 | #N/A | #N/A | #N/A | #N/A | #N/A | #N/A |
| AP001783.1 | 1.65E-06 | 0.08 | 0.06 | 0.003 | 0.048571 | 1.65 | #N/A | #N/A | #N/A | #N/A | #N/A | #N/A |
| AC113137.1 | 0.000309 | 0.23 | 0.06 | 0.008 | 1 | 1.63 | #N/A | #N/A | #N/A | #N/A | #N/A | #N/A |
| AC046129.1 | 0.002206 | 0.10 | 0.03 | 0.002 | 1 | 1.56 | #N/A | #N/A | #N/A | #N/A | #N/A | #N/A |
| LINC01426 | 0.00078 | 0.05 | 0.03 | 0.001 | 1 | 1.55 | #N/A | #N/A | #N/A | #N/A | #N/A | #N/A |
| HMGA2-AS1 | 1.83E-09 | 0.42 | 0.22 | 0.059 | 5.38E-05 | 1.53 | #N/A | #N/A | #N/A | #N/A | #N/A | #N/A |
| AC233296.1 | 2.82E-06 | 0.62 | 0.19 | 0.075 | 0.082704 | 1.53 | #N/A | #N/A | #N/A | #N/A | #N/A | #N/A |
| AC124798.1 | 0.003582 | 0.06 | 0.03 | 0.001 | 1 | 1.52 | #N/A | #N/A | #N/A | #N/A | #N/A | #N/A |
| Kcnmb4os2 | #N/A | #N/A | #N/A | #N/A | #N/A | #N/A | 2.28E-24 | 0.175222 | 0.094 | 0.005 | 4.15E-20 | 3.29 |
| 4930425O10Rik | #N/A | #N/A | #N/A | #N/A | #N/A | #N/A | 8.06E-25 | 0.158088 | 0.096 | 0.005 | 1.47E-20 | 3.04 |
| Gm26894 | #N/A | #N/A | #N/A | #N/A | #N/A | #N/A | 2.23E-10 | 0.07392 | 0.033 | 0.001 | 4.06E-06 | 2.44 |
| H2-Ab1 | #N/A | #N/A | #N/A | #N/A | #N/A | #N/A | 1.64E-22 | 0.307022 | 0.123 | 0.021 | 2.99E-18 | 1.80 |
| Micalcl | #N/A | #N/A | #N/A | #N/A | #N/A | #N/A | 1.13E-10 | 0.05015 | 0.035 | 0.001 | 2.06E-06 | 1.76 |
| **IFE Enriched** | | | | | | | | | | | | |
| LINC01956 | 8.04E-28 | -1.08635 | 0.019 | 0.412 | 2.36E-23 | -23.56 | #N/A | #N/A | #N/A | #N/A | #N/A | #N/A |
| PTPRQ | 7.06E-12 | -0.63305 | 0.013 | 0.199 | 2.07E-07 | -9.69 | #N/A | #N/A | #N/A | #N/A | #N/A | #N/A |
| AC087473.1 | 1.82E-16 | -0.74275 | 0.032 | 0.309 | 5.33E-12 | -7.17 | #N/A | #N/A | #N/A | #N/A | #N/A | #N/A |
| EPHA6 | 7.82E-26 | -1.67805 | 0.134 | 0.561 | 2.29E-21 | -7.03 | #N/A | #N/A | #N/A | #N/A | #N/A | #N/A |
| TMCO5A | 1.33E-07 | -0.32892 | 0.006 | 0.128 | 0.003899 | -7.02 | #N/A | #N/A | #N/A | #N/A | #N/A | #N/A |
| LINC02643 | 7.48E-31 | -1.28784 | 0.127 | 0.574 | 2.19E-26 | -5.82 | #N/A | #N/A | #N/A | #N/A | #N/A | #N/A |
| AC093606.1 | 6.18E-09 | -0.39395 | 0.013 | 0.165 | 0.000181 | -5.00 | #N/A | #N/A | #N/A | #N/A | #N/A | #N/A |
| LINC02345 | 3.80E-16 | -0.6976 | 0.051 | 0.328 | 1.12E-11 | -4.49 | #N/A | #N/A | #N/A | #N/A | #N/A | #N/A |
| MIR4500HG | 3.29E-10 | -0.46456 | 0.019 | 0.18 | 9.65E-06 | -4.40 | #N/A | #N/A | #N/A | #N/A | #N/A | #N/A |
| CCDC26 | 1.89E-16 | -1.35279 | 0.185 | 0.523 | 5.55E-12 | -3.82 | #N/A | #N/A | #N/A | #N/A | #N/A | #N/A |
| MAR01 | 2.98E-13 | -1.261 | 0.146 | 0.423 | 8.75E-09 | -3.65 | #N/A | #N/A | #N/A | #N/A | #N/A | #N/A |
| MEF2C-AS1 | 1.64E-16 | -1.04234 | 0.134 | 0.459 | 4.82E-12 | -3.57 | #N/A | #N/A | #N/A | #N/A | #N/A | #N/A |
| AC012349.1 | 4.05E-05 | -0.22043 | 0.006 | 0.09 | 1 | -3.31 | #N/A | #N/A | #N/A | #N/A | #N/A | #N/A |
| LINC01697 | 7.34E-14 | -0.77643 | 0.096 | 0.374 | 2.15E-09 | -3.02 | #N/A | #N/A | #N/A | #N/A | #N/A | #N/A |
| AC027251.1 | 2.43E-05 | -0.19232 | 0.006 | 0.093 | 0.714433 | -2.98 | #N/A | #N/A | #N/A | #N/A | #N/A | #N/A |
| KRT40 | 1.83E-09 | -0.44684 | 0.032 | 0.2 | 5.37E-05 | -2.79 | #N/A | #N/A | #N/A | #N/A | #N/A | #N/A |

| | | | | | | | | | | | | |
|---|---|---|---|---|---|---|---|---|---|---|---|---|
| AF165147.1 | 1.32E-12 | -0.95388 | 0.14 | 0.405 | 3.86E-08 | -2.76 | #N/A | #N/A | #N/A | #N/A | #N/A | #N/A |
| AC092810.3 | 0.001017 | -0.23239 | 0.006 | 0.067 | 1 | -2.60 | #N/A | #N/A | #N/A | #N/A | #N/A | #N/A |
| AP001037.1 | 8.40E-05 | -0.17836 | 0.006 | 0.086 | 1 | -2.56 | #N/A | #N/A | #N/A | #N/A | #N/A | #N/A |
| LAMB4 | 2.25E-27 | -1.2057 | 0.389 | 0.813 | 6.61E-23 | -2.52 | #N/A | #N/A | #N/A | #N/A | #N/A | #N/A |
| LINC01695 | 2.33E-12 | -0.86792 | 0.146 | 0.422 | 6.84E-08 | -2.51 | #N/A | #N/A | #N/A | #N/A | #N/A | #N/A |
| AL353586.1 | 1.90E-15 | -1.02832 | 0.21 | 0.48 | 5.57E-11 | -2.35 | #N/A | #N/A | #N/A | #N/A | #N/A | #N/A |
| AL161725.1 | 4.85E-05 | -0.15746 | 0.006 | 0.089 | 1 | -2.34 | #N/A | #N/A | #N/A | #N/A | #N/A | #N/A |
| DSG1-AS1 | 4.24E-06 | -0.24769 | 0.013 | 0.122 | 0.124518 | -2.32 | #N/A | #N/A | #N/A | #N/A | #N/A | #N/A |
| AL136456.1 | 0.008734 | -0.26207 | 0.006 | 0.052 | 1 | -2.27 | #N/A | #N/A | #N/A | #N/A | #N/A | #N/A |
| NGF-AS1 | 5.90E-14 | -0.68022 | 0.115 | 0.368 | 1.73E-09 | -2.18 | #N/A | #N/A | #N/A | #N/A | #N/A | #N/A |
| PICART1 | 0.000208 | -0.15899 | 0.006 | 0.077 | 1 | -2.04 | #N/A | #N/A | #N/A | #N/A | #N/A | #N/A |
| AC113383.1 | 0.000118 | -0.37181 | 0.019 | 0.104 | 1 | -2.04 | #N/A | #N/A | #N/A | #N/A | #N/A | #N/A |
| AL008633.1 | 7.99E-12 | -1.06815 | 0.248 | 0.472 | 2.35E-07 | -2.03 | #N/A | #N/A | #N/A | #N/A | #N/A | #N/A |
| LINC02595 | 1.39E-13 | -0.64206 | 0.134 | 0.423 | 4.09E-09 | -2.03 | #N/A | #N/A | #N/A | #N/A | #N/A | #N/A |
| NLRP7 | 0.000244 | -0.1631 | 0.006 | 0.074 | 1 | -2.01 | #N/A | #N/A | #N/A | #N/A | #N/A | #N/A |
| AC020916.1 | 3.87E-15 | -1.03402 | 0.312 | 0.603 | 1.14E-10 | -2.00 | #N/A | #N/A | #N/A | #N/A | #N/A | #N/A |
| AC104781.2 | 6.69E-09 | -0.54859 | 0.076 | 0.267 | 0.000196 | -1.93 | #N/A | #N/A | #N/A | #N/A | #N/A | #N/A |
| XAGE2 | 0.000621 | -0.15894 | 0.006 | 0.072 | 1 | -1.91 | #N/A | #N/A | #N/A | #N/A | #N/A | #N/A |
| AC068875.1 | 1.21E-06 | -0.32853 | 0.032 | 0.17 | 0.035387 | -1.75 | #N/A | #N/A | #N/A | #N/A | #N/A | #N/A |
| EYS | 1.82E-07 | -0.73809 | 0.134 | 0.308 | 0.00534 | -1.70 | #N/A | #N/A | #N/A | #N/A | #N/A | #N/A |
| AC092078.2 | 0.000236 | -0.34489 | 0.025 | 0.118 | 1 | -1.63 | #N/A | #N/A | #N/A | #N/A | #N/A | #N/A |
| AC083870.1 | 1.17E-16 | -0.98451 | 0.42 | 0.678 | 3.44E-12 | -1.59 | #N/A | #N/A | #N/A | #N/A | #N/A | #N/A |
| 3110099E03Rik | #N/A | #N/A | #N/A | #N/A | #N/A | #N/A | 1.81E-20 | -0.19934 | 0.017 | 0.151 | 3.30E-16 | -1.77 |
| Agrp | #N/A | #N/A | #N/A | #N/A | #N/A | #N/A | 1.82E-14 | -0.13863 | 0.008 | 0.099 | 3.31E-10 | -1.72 |

**Supplementary Table 14: Dermal condensate and dermis enriched genes identified in either the mouse (Qu et al., 2022) or human where no orthologue was found in the other species.**

| | Human | | | | | | Mouse (Qu et al., 2022) | | | | | |
|---|---|---|---|---|---|---|---|---|---|---|---|---|
| | p_val | avg_log2FC | pct.1 | pct.2 | p_val_adj | UScore | p_val | avg_log2FC | pct.1 | pct.2 | p_val_adj | UScore |
| **Dermal Condensate Enriched** | | | | | | | | | | | | |
| AC004594.1 | 2.71E-201 | 1.595039 | 0.51 | 0.007 | 7.95E-197 | 116.21 | #N/A | #N/A | #N/A | #N/A | #N/A | #N/A |
| AL450332.1 | 2.01E-185 | 2.046375 | 0.497 | 0.011 | 5.90E-181 | 92.46 | #N/A | #N/A | #N/A | #N/A | #N/A | #N/A |
| AC078923.1 | 2.60E-179 | 3.112673 | 0.514 | 0.026 | 7.64E-175 | 61.54 | #N/A | #N/A | #N/A | #N/A | #N/A | #N/A |
| AC078820.2 | 1.08E-53 | 0.50182 | 0.157 | 0.002 | 3.17E-49 | 39.39 | #N/A | #N/A | #N/A | #N/A | #N/A | #N/A |
| LINC00578 | 6.45E-135 | 2.346471 | 0.421 | 0.026 | 1.89E-130 | 37.99 | #N/A | #N/A | #N/A | #N/A | #N/A | #N/A |
| AC009803.1 | 5.29E-58 | 0.193831 | 0.16 | 0.001 | 1.55E-53 | 31.01 | #N/A | #N/A | #N/A | #N/A | #N/A | #N/A |
| AC008056.1 | 1.50E-121 | 0.300664 | 0.332 | 0.004 | 4.40E-117 | 24.96 | #N/A | #N/A | #N/A | #N/A | #N/A | #N/A |
| AC021613.1 | 4.12E-52 | 1.073199 | 0.175 | 0.008 | 1.21E-47 | 23.48 | #N/A | #N/A | #N/A | #N/A | #N/A | #N/A |
| NPS | 1.32E-25 | 0.303212 | 0.071 | 0.001 | 3.88E-21 | 21.53 | #N/A | #N/A | #N/A | #N/A | #N/A | #N/A |
| LINC02227 | 3.51E-94 | 1.379964 | 0.332 | 0.023 | 1.03E-89 | 19.92 | #N/A | #N/A | #N/A | #N/A | #N/A | #N/A |
| AC025437.4 | 1.15E-57 | 0.637783 | 0.18 | 0.007 | 3.38E-53 | 16.40 | #N/A | #N/A | #N/A | #N/A | #N/A | #N/A |
| INHBA-AS1 | 1.86E-30 | 0.309656 | 0.091 | 0.002 | 5.47E-26 | 14.09 | #N/A | #N/A | #N/A | #N/A | #N/A | #N/A |
| LINC02015 | 1.07E-21 | 0.414569 | 0.067 | 0.002 | 3.15E-17 | 13.89 | #N/A | #N/A | #N/A | #N/A | #N/A | #N/A |
| LINC01994 | 1.54E-25 | 0.317994 | 0.08 | 0.002 | 4.54E-21 | 12.72 | #N/A | #N/A | #N/A | #N/A | #N/A | #N/A |
| KIF26B-AS1 | 1.75E-121 | 1.509722 | 0.521 | 0.075 | 5.13E-117 | 10.49 | #N/A | #N/A | #N/A | #N/A | #N/A | #N/A |
| AL591518.1 | 2.66E-43 | 0.85209 | 0.171 | 0.014 | 7.80E-39 | 10.41 | #N/A | #N/A | #N/A | #N/A | #N/A | #N/A |
| LINC01995 | 4.26E-24 | 0.272031 | 0.075 | 0.002 | 1.25E-19 | 10.20 | #N/A | #N/A | #N/A | #N/A | #N/A | #N/A |
| AF064860.1 | 6.54E-73 | 1.112169 | 0.322 | 0.036 | 1.92E-68 | 9.95 | #N/A | #N/A | #N/A | #N/A | #N/A | #N/A |
| AC010230.1 | 4.05E-66 | 1.104793 | 0.313 | 0.045 | 1.19E-61 | 7.68 | #N/A | #N/A | #N/A | #N/A | #N/A | #N/A |
| LINC02596 | 3.71E-16 | 0.160962 | 0.046 | 0.001 | 1.09E-11 | 7.40 | #N/A | #N/A | #N/A | #N/A | #N/A | #N/A |
| C10orf71 | 2.76E-15 | 0.149739 | 0.047 | 0.001 | 8.09E-11 | 7.04 | #N/A | #N/A | #N/A | #N/A | #N/A | #N/A |
| LYPLAL1-DT | 1.80E-43 | 0.921172 | 0.233 | 0.034 | 5.30E-39 | 6.31 | #N/A | #N/A | #N/A | #N/A | #N/A | #N/A |
| Z99756.1 | 9.24E-20 | 0.262647 | 0.067 | 0.003 | 2.71E-15 | 5.87 | #N/A | #N/A | #N/A | #N/A | #N/A | #N/A |
| HES4 | 3.52E-101 | 0.896122 | 0.488 | 0.076 | 1.03E-96 | 5.75 | #N/A | #N/A | #N/A | #N/A | #N/A | #N/A |
| AL096794.1 | 1.22E-25 | 0.574672 | 0.107 | 0.011 | 3.60E-21 | 5.59 | #N/A | #N/A | #N/A | #N/A | #N/A | #N/A |
| LYPLAL1-AS1 | 4.80E-49 | 0.957777 | 0.25 | 0.044 | 1.41E-44 | 5.44 | #N/A | #N/A | #N/A | #N/A | #N/A | #N/A |

| | | | | | | | | | | | | |
|---|---|---|---|---|---|---|---|---|---|---|---|---|
| AC117461.1 | 3.78E-16 | 0.216383 | 0.049 | 0.002 | 1.11E-11 | 5.30 | #N/A | #N/A | #N/A | #N/A | #N/A | #N/A |
| LINC02607 | 4.24E-35 | 0.72144 | 0.189 | 0.026 | 1.25E-30 | 5.24 | #N/A | #N/A | #N/A | #N/A | #N/A | #N/A |
| DEC01 | 3.86E-106 | 1.5182 | 0.641 | 0.189 | 1.13E-101 | 5.15 | #N/A | #N/A | #N/A | #N/A | #N/A | #N/A |
| AC116345.1 | 8.44E-88 | 1.459694 | 0.641 | 0.195 | 2.48E-83 | 4.80 | #N/A | #N/A | #N/A | #N/A | #N/A | #N/A |
| AC021055.1 | 4.31E-19 | 0.151502 | 0.058 | 0.002 | 1.27E-14 | 4.39 | #N/A | #N/A | #N/A | #N/A | #N/A | #N/A |
| AL449403.2 | 6.91E-23 | 0.254637 | 0.086 | 0.005 | 2.03E-18 | 4.38 | #N/A | #N/A | #N/A | #N/A | #N/A | #N/A |
| AC061958.1 | 2.35E-29 | 0.515198 | 0.149 | 0.018 | 6.88E-25 | 4.26 | #N/A | #N/A | #N/A | #N/A | #N/A | #N/A |
| AC097450.1 | 3.34E-19 | 0.309397 | 0.08 | 0.006 | 9.80E-15 | 4.13 | #N/A | #N/A | #N/A | #N/A | #N/A | #N/A |
| AC092958.1 | 9.95E-40 | 0.89758 | 0.242 | 0.059 | 2.92E-35 | 3.68 | #N/A | #N/A | #N/A | #N/A | #N/A | #N/A |
| AC025437.2 | 1.64E-11 | 0.101791 | 0.036 | 0.001 | 4.82E-07 | 3.66 | #N/A | #N/A | #N/A | #N/A | #N/A | #N/A |
| LINC00670 | 1.27E-21 | 0.310657 | 0.093 | 0.008 | 3.72E-17 | 3.61 | #N/A | #N/A | #N/A | #N/A | #N/A | #N/A |
| AC091979.1 | 1.05E-23 | 0.272031 | 0.089 | 0.007 | 3.08E-19 | 3.46 | #N/A | #N/A | #N/A | #N/A | #N/A | #N/A |
| AC011853.2 | 1.92E-26 | 0.249424 | 0.109 | 0.008 | 5.64E-22 | 3.40 | #N/A | #N/A | #N/A | #N/A | #N/A | #N/A |
| AC098588.3 | 3.90E-50 | 1.049457 | 0.452 | 0.147 | 1.15E-45 | 3.23 | #N/A | #N/A | #N/A | #N/A | #N/A | #N/A |
| AP003049.2 | 3.38E-25 | 0.219245 | 0.102 | 0.007 | 9.93E-21 | 3.19 | #N/A | #N/A | #N/A | #N/A | #N/A | #N/A |
| AC022613.1 | 4.60E-16 | 0.202225 | 0.062 | 0.004 | 1.35E-11 | 3.13 | #N/A | #N/A | #N/A | #N/A | #N/A | #N/A |
| AC025437.3 | 9.39E-15 | 0.173667 | 0.053 | 0.003 | 2.76E-10 | 3.07 | #N/A | #N/A | #N/A | #N/A | #N/A | #N/A |
| AC008780.1 | 8.36E-11 | 0.086993 | 0.035 | 0.001 | 2.45E-06 | 3.04 | #N/A | #N/A | #N/A | #N/A | #N/A | #N/A |
| THSD4-AS1 | 7.00E-70 | 1.229058 | 0.599 | 0.245 | 2.05E-65 | 3.00 | #N/A | #N/A | #N/A | #N/A | #N/A | #N/A |
| AC016924.1 | 9.92E-08 | 0.112845 | 0.026 | 0.001 | 0.002911 | 2.93 | #N/A | #N/A | #N/A | #N/A | #N/A | #N/A |
| LINC02223 | 4.24E-21 | 0.295276 | 0.098 | 0.01 | 1.25E-16 | 2.89 | #N/A | #N/A | #N/A | #N/A | #N/A | #N/A |
| AC136424.2 | 1.25E-52 | 0.706315 | 0.335 | 0.082 | 3.66E-48 | 2.89 | #N/A | #N/A | #N/A | #N/A | #N/A | #N/A |
| AC013287.1 | 2.71E-20 | 0.344393 | 0.098 | 0.012 | 7.97E-16 | 2.81 | #N/A | #N/A | #N/A | #N/A | #N/A | #N/A |
| HHIP-AS1 | 6.65E-20 | 0.226603 | 0.082 | 0.007 | 1.95E-15 | 2.65 | #N/A | #N/A | #N/A | #N/A | #N/A | #N/A |
| LINC02836 | 5.24E-15 | 0.209483 | 0.062 | 0.005 | 1.54E-10 | 2.60 | #N/A | #N/A | #N/A | #N/A | #N/A | #N/A |
| ABO | 9.40E-33 | 0.053226 | 0.097 | 0.002 | 2.76E-28 | 2.58 | #N/A | #N/A | #N/A | #N/A | #N/A | #N/A |
| AC064874.1 | 9.46E-27 | 0.459809 | 0.142 | 0.027 | 2.78E-22 | 2.42 | #N/A | #N/A | #N/A | #N/A | #N/A | #N/A |
| AC110023.1 | 1.57E-43 | 0.408532 | 0.259 | 0.044 | 4.60E-39 | 2.40 | #N/A | #N/A | #N/A | #N/A | #N/A | #N/A |
| SOX2-OT | 5.04E-34 | 0.683596 | 0.311 | 0.089 | 1.48E-29 | 2.39 | #N/A | #N/A | #N/A | #N/A | #N/A | #N/A |
| AC025839.1 | 3.36E-09 | 0.152908 | 0.031 | 0.002 | 9.88E-05 | 2.37 | #N/A | #N/A | #N/A | #N/A | #N/A | #N/A |
| FGF10-AS1 | 8.32E-31 | 0.506997 | 0.23 | 0.051 | 2.44E-26 | 2.29 | #N/A | #N/A | #N/A | #N/A | #N/A | #N/A |
| LINC02357 | 8.43E-11 | 0.155159 | 0.042 | 0.003 | 2.47E-06 | 2.17 | #N/A | #N/A | #N/A | #N/A | #N/A | #N/A |

| | | | | | | | | | | | | |
|---|---|---|---|---|---|---|---|---|---|---|---|---|
| DIO3OS | 8.83E-20 | 0.382905 | 0.106 | 0.019 | 2.59E-15 | 2.14 | #N/A | #N/A | #N/A | #N/A | #N/A | #N/A |
| AC092422.1 | 1.26E-27 | 0.852933 | 0.321 | 0.129 | 3.68E-23 | 2.12 | #N/A | #N/A | #N/A | #N/A | #N/A | #N/A |
| AL590999.1 | 3.42E-53 | 0.944332 | 0.49 | 0.231 | 1.00E-48 | 2.00 | #N/A | #N/A | #N/A | #N/A | #N/A | #N/A |
| LINC01416 | 2.90E-31 | 0.906894 | 0.355 | 0.161 | 8.53E-27 | 2.00 | #N/A | #N/A | #N/A | #N/A | #N/A | #N/A |
| AC005358.1 | 6.19E-15 | 0.233395 | 0.075 | 0.009 | 1.82E-10 | 1.94 | #N/A | #N/A | #N/A | #N/A | #N/A | #N/A |
| LINC02456 | 2.07E-13 | 0.144821 | 0.053 | 0.004 | 6.09E-09 | 1.92 | #N/A | #N/A | #N/A | #N/A | #N/A | #N/A |
| AL513487.1 | 1.69E-10 | 0.10725 | 0.035 | 0.002 | 4.96E-06 | 1.88 | #N/A | #N/A | #N/A | #N/A | #N/A | #N/A |
| LINC02224 | 6.15E-19 | 0.410057 | 0.146 | 0.033 | 1.81E-14 | 1.81 | #N/A | #N/A | #N/A | #N/A | #N/A | #N/A |
| AC008056.2 | 7.87E-27 | 0.043779 | 0.082 | 0.002 | 2.31E-22 | 1.79 | #N/A | #N/A | #N/A | #N/A | #N/A | #N/A |
| RFLNA | 5.70E-36 | 0.909346 | 0.477 | 0.263 | 1.67E-31 | 1.65 | #N/A | #N/A | #N/A | #N/A | #N/A | #N/A |
| LINC01579 | 2.09E-32 | 0.204723 | 0.16 | 0.02 | 6.13E-28 | 1.64 | #N/A | #N/A | #N/A | #N/A | #N/A | #N/A |
| AC005224.2 | 2.06E-23 | 0.214671 | 0.122 | 0.016 | 6.06E-19 | 1.64 | #N/A | #N/A | #N/A | #N/A | #N/A | #N/A |
| LINC01122 | 1.14E-21 | 0.598703 | 0.262 | 0.096 | 3.34E-17 | 1.63 | #N/A | #N/A | #N/A | #N/A | #N/A | #N/A |
| AF123462.1 | 4.89E-14 | 0.036166 | 0.044 | 0.001 | 1.43E-09 | 1.59 | #N/A | #N/A | #N/A | #N/A | #N/A | #N/A |
| SEMA6A-AS1 | 2.48E-45 | 0.805341 | 0.612 | 0.313 | 7.27E-41 | 1.57 | #N/A | #N/A | #N/A | #N/A | #N/A | #N/A |
| CYP27C1 | 4.57E-32 | 0.691355 | 0.403 | 0.177 | 1.34E-27 | 1.57 | #N/A | #N/A | #N/A | #N/A | #N/A | #N/A |
| MEP1B | 8.19E-20 | 0.364814 | 0.153 | 0.036 | 2.40E-15 | 1.55 | #N/A | #N/A | #N/A | #N/A | #N/A | #N/A |
| PTPRN2 | 3.16E-41 | 0.579229 | 0.47 | 0.177 | 9.28E-37 | 1.54 | #N/A | #N/A | #N/A | #N/A | #N/A | #N/A |
| AC091939.1 | 3.19E-12 | 0.209349 | 0.066 | 0.009 | 9.35E-08 | 1.54 | #N/A | #N/A | #N/A | #N/A | #N/A | #N/A |
| 2700046A07Rik | #N/A | #N/A | #N/A | #N/A | #N/A | #N/A | 6.02E-71 | 0.771999 | 0.266 | 0.009 | 1.10E-66 | 22.82 |
| Gm11541 | #N/A | #N/A | #N/A | #N/A | #N/A | #N/A | 4.60E-39 | 0.498379 | 0.161 | 0.006 | 8.39E-35 | 13.37 |
| H2-DMb1 | #N/A | #N/A | #N/A | #N/A | #N/A | #N/A | 8.25E-20 | 0.132112 | 0.069 | 0.001 | 1.50E-15 | 9.12 |
| Fgf3 | #N/A | #N/A | #N/A | #N/A | #N/A | #N/A | 1.75E-17 | 0.205921 | 0.075 | 0.003 | 3.18E-13 | 5.15 |
| 6330403K07Rik | #N/A | #N/A | #N/A | #N/A | #N/A | #N/A | 1.23E-128 | 1.468889 | 0.707 | 0.248 | 2.24E-124 | 4.19 |
| Gml2 | #N/A | #N/A | #N/A | #N/A | #N/A | #N/A | 4.62E-09 | 0.108248 | 0.036 | 0.001 | 8.42E-05 | 3.90 |
| H2-DMb2 | #N/A | #N/A | #N/A | #N/A | #N/A | #N/A | 9.85E-14 | 0.078686 | 0.048 | 0.001 | 1.79E-09 | 3.78 |
| Gm26644 | #N/A | #N/A | #N/A | #N/A | #N/A | #N/A | 7.73E-09 | 0.111564 | 0.033 | 0.001 | 0.000141 | 3.68 |
| H2-DMa | #N/A | #N/A | #N/A | #N/A | #N/A | #N/A | 1.07E-25 | 0.459102 | 0.167 | 0.041 | 1.96E-21 | 1.87 |
| **Dermis Enriched** | | | | | | | | | | | | |
| VSIG10L2 | 2.46E-36 | -0.37689 | 0.002 | 0.165 | 7.23E-32 | -31.09 | #N/A | #N/A | #N/A | #N/A | #N/A | #N/A |
| ZNF385D-AS1 | 1.33E-51 | -0.64119 | 0.015 | 0.265 | 3.90E-47 | -11.33 | #N/A | #N/A | #N/A | #N/A | #N/A | #N/A |

| | | | | | | | | | | | | |
|---|---|---|---|---|---|---|---|---|---|---|---|---|
| LINC01725 | 1.47E-54 | -0.96451 | 0.027 | 0.302 | 4.31E-50 | -10.79 | #N/A | #N/A | #N/A | #N/A | #N/A | #N/A |
| AC079298.3 | 7.44E-88 | -1.26963 | 0.058 | 0.484 | 2.18E-83 | -10.59 | #N/A | #N/A | #N/A | #N/A | #N/A | #N/A |
| ZNF385D | 6.06E-234 | -3.31898 | 0.306 | 0.882 | 1.78E-229 | -9.57 | #N/A | #N/A | #N/A | #N/A | #N/A | #N/A |
| CLEC2A | 3.95E-27 | -0.47355 | 0.007 | 0.139 | 1.16E-22 | -9.40 | #N/A | #N/A | #N/A | #N/A | #N/A | #N/A |
| LINC01807 | 3.90E-14 | -0.25939 | 0.002 | 0.071 | 1.15E-09 | -9.21 | #N/A | #N/A | #N/A | #N/A | #N/A | #N/A |
| AL355612.1 | 3.47E-36 | -1.21615 | 0.044 | 0.252 | 1.02E-31 | -6.97 | #N/A | #N/A | #N/A | #N/A | #N/A | #N/A |
| SLC9A2 | 5.99E-38 | -0.9516 | 0.04 | 0.267 | 1.76E-33 | -6.35 | #N/A | #N/A | #N/A | #N/A | #N/A | #N/A |
| LINC01697 | 1.56E-31 | -0.59598 | 0.018 | 0.191 | 4.57E-27 | -6.32 | #N/A | #N/A | #N/A | #N/A | #N/A | #N/A |
| LINC01550 | 8.92E-44 | -0.65044 | 0.033 | 0.279 | 2.62E-39 | -5.50 | #N/A | #N/A | #N/A | #N/A | #N/A | #N/A |
| Z99289.1 | 5.45E-43 | -0.64036 | 0.036 | 0.284 | 1.60E-38 | -5.05 | #N/A | #N/A | #N/A | #N/A | #N/A | #N/A |
| AP000676.5 | 1.16E-28 | -0.84382 | 0.035 | 0.209 | 3.40E-24 | -5.04 | #N/A | #N/A | #N/A | #N/A | #N/A | #N/A |
| AL138689.2 | 1.10E-40 | -0.59907 | 0.033 | 0.267 | 3.24E-36 | -4.85 | #N/A | #N/A | #N/A | #N/A | #N/A | #N/A |
| LINC01362 | 7.63E-34 | -0.79155 | 0.042 | 0.244 | 2.24E-29 | -4.60 | #N/A | #N/A | #N/A | #N/A | #N/A | #N/A |
| AL360178.1 | 2.97E-38 | -0.66651 | 0.04 | 0.267 | 8.72E-34 | -4.45 | #N/A | #N/A | #N/A | #N/A | #N/A | #N/A |
| AL138828.1 | 7.73E-65 | -1.30194 | 0.168 | 0.555 | 2.27E-60 | -4.30 | #N/A | #N/A | #N/A | #N/A | #N/A | #N/A |
| LINC02624 | 4.66E-17 | -0.31112 | 0.007 | 0.096 | 1.37E-12 | -4.27 | #N/A | #N/A | #N/A | #N/A | #N/A | #N/A |
| AC099753.1 | 4.87E-29 | -0.66042 | 0.033 | 0.213 | 1.43E-24 | -4.26 | #N/A | #N/A | #N/A | #N/A | #N/A | #N/A |
| AL133346.1 | 1.48E-42 | -0.97153 | 0.08 | 0.347 | 4.34E-38 | -4.21 | #N/A | #N/A | #N/A | #N/A | #N/A | #N/A |
| EPHA6 | 1.28E-55 | -1.45043 | 0.179 | 0.52 | 3.77E-51 | -4.21 | #N/A | #N/A | #N/A | #N/A | #N/A | #N/A |
| AC099066.2 | 7.54E-23 | -0.38167 | 0.013 | 0.138 | 2.21E-18 | -4.05 | #N/A | #N/A | #N/A | #N/A | #N/A | #N/A |
| SLC7A14-AS1 | 1.76E-11 | -0.24745 | 0.004 | 0.064 | 5.17E-07 | -3.96 | #N/A | #N/A | #N/A | #N/A | #N/A | #N/A |
| MIR100HG | 4.45E-95 | -1.25762 | 0.23 | 0.704 | 1.31E-90 | -3.85 | #N/A | #N/A | #N/A | #N/A | #N/A | #N/A |
| GPC6-AS2 | 2.68E-15 | -0.18432 | 0.004 | 0.081 | 7.87E-11 | -3.73 | #N/A | #N/A | #N/A | #N/A | #N/A | #N/A |
| GPC6-AS1 | 1.38E-23 | -0.29212 | 0.011 | 0.14 | 4.06E-19 | -3.72 | #N/A | #N/A | #N/A | #N/A | #N/A | #N/A |
| LINC02063 | 2.55E-72 | -1.0598 | 0.169 | 0.589 | 7.48E-68 | -3.69 | #N/A | #N/A | #N/A | #N/A | #N/A | #N/A |
| SAMMSON | 4.91E-10 | -0.25566 | 0.004 | 0.057 | 1.44E-05 | -3.64 | #N/A | #N/A | #N/A | #N/A | #N/A | #N/A |
| AC097528.1 | 1.97E-284 | -2.3972 | 0.641 | 0.973 | 5.80E-280 | -3.64 | #N/A | #N/A | #N/A | #N/A | #N/A | #N/A |
| AC016877.1 | 2.12E-37 | -0.60581 | 0.046 | 0.276 | 6.23E-33 | -3.63 | #N/A | #N/A | #N/A | #N/A | #N/A | #N/A |
| SAMD4A | 1.63E-168 | -1.77741 | 0.461 | 0.933 | 4.77E-164 | -3.60 | #N/A | #N/A | #N/A | #N/A | #N/A | #N/A |
| AC093765.3 | 3.92E-33 | -0.71903 | 0.055 | 0.273 | 1.15E-28 | -3.57 | #N/A | #N/A | #N/A | #N/A | #N/A | #N/A |
| C1orf198 | 2.35E-46 | -0.69513 | 0.069 | 0.354 | 6.90E-42 | -3.57 | #N/A | #N/A | #N/A | #N/A | #N/A | #N/A |
| MEF2C-AS1 | 3.43E-59 | -1.21198 | 0.209 | 0.586 | 1.01E-54 | -3.40 | #N/A | #N/A | #N/A | #N/A | #N/A | #N/A |

| | | | | | | | | | | | | |
|---|---|---|---|---|---|---|---|---|---|---|---|---|
| CDHR3 | 4.12E-37 | -0.7282 | 0.069 | 0.311 | 1.21E-32 | -3.28 | #N/A | #N/A | #N/A | #N/A | #N/A | #N/A |
| AC107072.2 | 8.55E-44 | -0.78975 | 0.093 | 0.384 | 2.51E-39 | -3.26 | #N/A | #N/A | #N/A | #N/A | #N/A | #N/A |
| HMGA2-AS1 | 8.47E-24 | -0.32763 | 0.016 | 0.153 | 2.49E-19 | -3.13 | #N/A | #N/A | #N/A | #N/A | #N/A | #N/A |
| AP002989.1 | 1.37E-16 | -0.54283 | 0.024 | 0.133 | 4.03E-12 | -3.01 | #N/A | #N/A | #N/A | #N/A | #N/A | #N/A |
| SLC22A24 | 8.77E-09 | -0.13024 | 0.002 | 0.045 | 0.000257 | -2.93 | #N/A | #N/A | #N/A | #N/A | #N/A | #N/A |
| LINC01695 | 7.21E-24 | -0.54991 | 0.038 | 0.201 | 2.12E-19 | -2.91 | #N/A | #N/A | #N/A | #N/A | #N/A | #N/A |
| ANKFN1 | 1.65E-30 | -0.62448 | 0.058 | 0.269 | 4.84E-26 | -2.90 | #N/A | #N/A | #N/A | #N/A | #N/A | #N/A |
| NLRC5 | 1.45E-17 | -0.51618 | 0.027 | 0.147 | 4.25E-13 | -2.81 | #N/A | #N/A | #N/A | #N/A | #N/A | #N/A |
| SLC22A10 | 4.09E-15 | -0.34371 | 0.013 | 0.1 | 1.20E-10 | -2.64 | #N/A | #N/A | #N/A | #N/A | #N/A | #N/A |
| C1orf21 | 2.18E-69 | -1.28912 | 0.364 | 0.735 | 6.39E-65 | -2.60 | #N/A | #N/A | #N/A | #N/A | #N/A | #N/A |
| AC002454.1 | 2.21E-10 | -0.20189 | 0.005 | 0.064 | 6.50E-06 | -2.58 | #N/A | #N/A | #N/A | #N/A | #N/A | #N/A |
| IPO9-AS1 | 1.61E-49 | -0.98386 | 0.211 | 0.553 | 4.74E-45 | -2.58 | #N/A | #N/A | #N/A | #N/A | #N/A | #N/A |
| KLRF2 | 1.92E-11 | -0.16104 | 0.004 | 0.062 | 5.64E-07 | -2.50 | #N/A | #N/A | #N/A | #N/A | #N/A | #N/A |
| AC022034.1 | 2.36E-11 | -0.15472 | 0.004 | 0.064 | 6.92E-07 | -2.48 | #N/A | #N/A | #N/A | #N/A | #N/A | #N/A |
| AC008522.1 | 8.11E-31 | -0.52684 | 0.058 | 0.271 | 2.38E-26 | -2.46 | #N/A | #N/A | #N/A | #N/A | #N/A | #N/A |
| LINC01060 | 2.11E-11 | -0.66748 | 0.033 | 0.12 | 6.21E-07 | -2.43 | #N/A | #N/A | #N/A | #N/A | #N/A | #N/A |
| AC003991.1 | 1.58E-10 | -0.18631 | 0.005 | 0.065 | 4.64E-06 | -2.42 | #N/A | #N/A | #N/A | #N/A | #N/A | #N/A |
| SEPTIN11 | 1.14E-151 | -1.59603 | 0.632 | 0.951 | 3.35E-147 | -2.40 | #N/A | #N/A | #N/A | #N/A | #N/A | #N/A |
| LINC02284 | 2.70E-20 | -0.3512 | 0.022 | 0.15 | 7.92E-16 | -2.39 | #N/A | #N/A | #N/A | #N/A | #N/A | #N/A |
| SEPTIN9 | 9.04E-69 | -1.08336 | 0.33 | 0.719 | 2.65E-64 | -2.36 | #N/A | #N/A | #N/A | #N/A | #N/A | #N/A |
| AC020611.2 | 1.62E-29 | -0.55091 | 0.064 | 0.274 | 4.74E-25 | -2.36 | #N/A | #N/A | #N/A | #N/A | #N/A | #N/A |
| RGMB-AS1 | 3.55E-15 | -0.17696 | 0.007 | 0.092 | 1.04E-10 | -2.33 | #N/A | #N/A | #N/A | #N/A | #N/A | #N/A |
| AL133415.1 | 1.50E-35 | -0.57187 | 0.084 | 0.335 | 4.39E-31 | -2.28 | #N/A | #N/A | #N/A | #N/A | #N/A | #N/A |
| AC083855.2 | 1.38E-16 | -0.29848 | 0.016 | 0.121 | 4.06E-12 | -2.26 | #N/A | #N/A | #N/A | #N/A | #N/A | #N/A |
| GBP1 | 6.67E-15 | -0.20984 | 0.009 | 0.096 | 1.96E-10 | -2.24 | #N/A | #N/A | #N/A | #N/A | #N/A | #N/A |
| AF165147.1 | 9.28E-24 | -0.80656 | 0.124 | 0.323 | 2.72E-19 | -2.10 | #N/A | #N/A | #N/A | #N/A | #N/A | #N/A |
| SGO1-AS1 | 1.12E-24 | -1.02156 | 0.219 | 0.449 | 3.28E-20 | -2.09 | #N/A | #N/A | #N/A | #N/A | #N/A | #N/A |
| AC007099.1 | 8.77E-24 | -0.49197 | 0.055 | 0.231 | 2.58E-19 | -2.07 | #N/A | #N/A | #N/A | #N/A | #N/A | #N/A |
| AC093765.2 | 3.92E-18 | -0.3799 | 0.029 | 0.155 | 1.15E-13 | -2.03 | #N/A | #N/A | #N/A | #N/A | #N/A | #N/A |
| LHFPL6 | 2.09E-67 | -1.04548 | 0.428 | 0.803 | 6.13E-63 | -1.96 | #N/A | #N/A | #N/A | #N/A | #N/A | #N/A |
| MEF2C-AS2 | 4.19E-26 | -0.49676 | 0.067 | 0.261 | 1.23E-21 | -1.94 | #N/A | #N/A | #N/A | #N/A | #N/A | #N/A |
| AC068725.1 | 1.04E-14 | -0.21239 | 0.011 | 0.099 | 3.06E-10 | -1.91 | #N/A | #N/A | #N/A | #N/A | #N/A | #N/A |

| | | | | | | | | | | | | |
|---|---|---|---|---|---|---|---|---|---|---|---|---|
| LINC01950 | 4.58E-14 | -0.49503 | 0.04 | 0.153 | 1.35E-09 | -1.89 | #N/A | #N/A | #N/A | #N/A | #N/A | #N/A |
| CELF2-AS2 | 4.20E-08 | -0.08944 | 0.002 | 0.042 | 0.001234 | -1.88 | #N/A | #N/A | #N/A | #N/A | #N/A | #N/A |
| AL360178.2 | 7.55E-12 | -0.17193 | 0.007 | 0.076 | 2.22E-07 | -1.87 | #N/A | #N/A | #N/A | #N/A | #N/A | #N/A |
| KIAA1217 | 1.21E-44 | -0.93592 | 0.337 | 0.656 | 3.56E-40 | -1.82 | #N/A | #N/A | #N/A | #N/A | #N/A | #N/A |
| BX284613.2 | 5.58E-18 | -0.73681 | 0.071 | 0.175 | 1.64E-13 | -1.82 | #N/A | #N/A | #N/A | #N/A | #N/A | #N/A |
| AL713852.1 | 1.32E-16 | -0.21161 | 0.013 | 0.11 | 3.86E-12 | -1.79 | #N/A | #N/A | #N/A | #N/A | #N/A | #N/A |
| AC004551.1 | 6.25E-10 | -0.14507 | 0.005 | 0.061 | 1.84E-05 | -1.77 | #N/A | #N/A | #N/A | #N/A | #N/A | #N/A |
| AC106845.1 | 2.43E-34 | -0.64334 | 0.16 | 0.437 | 7.13E-30 | -1.76 | #N/A | #N/A | #N/A | #N/A | #N/A | #N/A |
| AL365214.2 | 8.28E-21 | -0.37 | 0.04 | 0.189 | 2.43E-16 | -1.75 | #N/A | #N/A | #N/A | #N/A | #N/A | #N/A |
| MIR193BHG | 3.55E-23 | -0.53613 | 0.087 | 0.276 | 1.04E-18 | -1.70 | #N/A | #N/A | #N/A | #N/A | #N/A | #N/A |
| LINC02802 | 1.68E-25 | -0.39842 | 0.058 | 0.246 | 4.94E-21 | -1.69 | #N/A | #N/A | #N/A | #N/A | #N/A | #N/A |
| PWRN1 | 4.59E-17 | -0.37874 | 0.033 | 0.147 | 1.35E-12 | -1.69 | #N/A | #N/A | #N/A | #N/A | #N/A | #N/A |
| AC099792.1 | 8.50E-10 | -0.18989 | 0.007 | 0.062 | 2.50E-05 | -1.68 | #N/A | #N/A | #N/A | #N/A | #N/A | #N/A |
| HLA-E | 1.53E-15 | -0.32828 | 0.026 | 0.133 | 4.48E-11 | -1.68 | #N/A | #N/A | #N/A | #N/A | #N/A | #N/A |
| AC005550.2 | 4.08E-17 | -0.2349 | 0.015 | 0.107 | 1.20E-12 | -1.68 | #N/A | #N/A | #N/A | #N/A | #N/A | #N/A |
| AC093523.1 | 4.56E-07 | -0.15715 | 0.004 | 0.042 | 0.013374 | -1.65 | #N/A | #N/A | #N/A | #N/A | #N/A | #N/A |
| PARP15 | 4.20E-14 | -0.24684 | 0.018 | 0.113 | 1.23E-09 | -1.55 | #N/A | #N/A | #N/A | #N/A | #N/A | #N/A |
| AC020637.1 | 4.77E-12 | -0.57429 | 0.067 | 0.178 | 1.40E-07 | -1.53 | #N/A | #N/A | #N/A | #N/A | #N/A | #N/A |
| AC090994.1 | 1.84E-15 | -0.3377 | 0.033 | 0.148 | 5.40E-11 | -1.51 | #N/A | #N/A | #N/A | #N/A | #N/A | #N/A |
| SPRY4-AS1 | 1.43E-18 | -0.55523 | 0.093 | 0.253 | 4.20E-14 | -1.51 | #N/A | #N/A | #N/A | #N/A | #N/A | #N/A |
| AP000331.1 | 1.33E-21 | -0.40194 | 0.062 | 0.232 | 3.91E-17 | -1.50 | #N/A | #N/A | #N/A | #N/A | #N/A | #N/A |
| Abca8a | #N/A | #N/A | #N/A | #N/A | #N/A | #N/A | 3.78E-23 | -0.4328 | 0.054 | 0.278 | 6.88E-19 | -2.23 |
| Lockd | #N/A | #N/A | #N/A | #N/A | #N/A | #N/A | 7.08E-18 | -0.5006 | 0.063 | 0.254 | 1.29E-13 | -2.02 |
| 5033430I15Rik | #N/A | #N/A | #N/A | #N/A | #N/A | #N/A | 4.52E-10 | -0.18141 | 0.009 | 0.095 | 8.24E-06 | -1.91 |
| Fam46a | #N/A | #N/A | #N/A | #N/A | #N/A | #N/A | 6.31E-38 | -0.78331 | 0.278 | 0.651 | 1.15E-33 | -1.83 |
| Agtr1a | #N/A | #N/A | #N/A | #N/A | #N/A | #N/A | 6.29E-10 | -0.15809 | 0.009 | 0.093 | 1.15E-05 | -1.63 |
| Sgol2a | #N/A | #N/A | #N/A | #N/A | #N/A | #N/A | 4.73E-10 | -0.32763 | 0.03 | 0.138 | 8.62E-06 | -1.51 |

#### Supplementary information – Samples and statistics

##### Figure 1e – Hair follicle stage

Number of samples used for each area per age. Skin samples were taken from 44 individual human fetal specimens.

| | | Anatomical Site | | | | | | | |
|---|---|---|---|---|---|---|---|---|---|
| | | Back | Cheek | Chin | Eyebrow | Forehead | Front | Scalp | Upper Lip |
| Weeks Estimated Gestational Age | 10 | 4 | 1 | 1 | 2 | 3 | | 3 | 2 |
| | 11 | 1 | 1 | 1 | 1 | 1 | 1 | 1 | 2 |
| | 12 | 7 | 6 | 6 | 6 | 7 | 4 | 7 | 7 |
| | 13 | 6 | 2 | 1 | 1 | 2 | 4 | 4 | 3 |
| | 14 | 6 | 5 | 4 | 5 | 5 | 6 | 6 | 4 |
| | 15 | 1 | 1 | 1 | 1 | 1 | 0 | 1 | 1 |
| | 16 | 7 | 3 | 4 | 3 | 6 | 5 | 6 | 4 |
| | 17 | 3 | 2 | 2 | 2 | 2 | 3 | 2 | 2 |
| | 18 | 1 | | | | | 1 | | |
| | 19 | 2 | 1 | 1 | 1 | 1 | 1 | 1 | 1 |

**Figure 2a – Placode diameter**

i. Ordinary one-way ANOVA between anatomical sites

**F (8, 156) = 0.9025, P = 0.5160**

ii. Tukey's multiple comparisons test

| | | Mouse | Human | | | | | | |
|---|---|---|---|---|---|---|---|---|---|
| | | Back | Back | Cheek | Chin | Eyebrow | Forehead | Front | Scalp |
| Human | Back | >0.9999 | | | | | | | |
| | Cheek | >0.9999 | 0.9987 | | | | | | |
| | Chin | 0.9962 | 0.9777 | >0.9999 | | | | | |
| | Eyebrow | 0.9873 | 0.9404 | 0.9997 | >0.9999 | | | | |
| | Forehead | >0.9999 | >0.9999 | 0.9937 | 0.9425 | 0.8705 | | | |
| | Front | 0.9984 | 0.9872 | >0.9999 | >0.9999 | >0.9999 | 0.9617 | | |
| | Scalp | >0.9999 | >0.9999 | >0.9999 | 0.9976 | 0.989 | 0.9998 | 0.9992 | |
| | Upper Lip | 0.8617 | 0.6511 | 0.9706 | 0.9992 | 0.9998 | 0.4912 | 0.9963 | 0.8191 |

iii. Number of samples of each age per area. Skin samples were taken from 33 individual human fetal specimens and 12 mouse specimens.

| | | Anatomical Site | | | | | | | |
|---|---|---|---|---|---|---|---|---|---|
| | | Back | Cheek | Chin | Eyebrow | Forehead | Front | Scalp | Upper Lip |
| Weeks Estimated Gestational Age | 10 | | | | 1 | | | | 2 |
| | 11 | | | | | | | | 1 |
| | 12 | | 4 | 3 | 5 | 2 | | 4 | 7 |
| | 13 | 1 | 2 | 1 | 1 | 2 | 3 | 3 | 3 |
| | 14 | 5 | 5 | 4 | 5 | 5 | 5 | 5 | 4 |
| | 15 | 1 | 1 | 1 | 1 | 1 | | 1 | 1 |
| | 16 | 5 | 3 | 4 | 3 | 6 | 5 | 6 | 3 |
| | 17 | 2 | 2 | 2 | 2 | 2 | 3 | 2 | 2 |
| | 18 | 1 | | | | | 1 | | |
| | 19 | 2 | 1 | 1 | 1 | 1 | 1 | 1 | 1 |
| Mouse | E13.75 | 1 | | | | | | | |
| | E14 | 7 | | | | | | | |
| | E15 | 4 | | | | | | | |

**Figure 2b – Placode density**

i. Ordinary one-way ANOVA between anatomical sites

**$F (8,56) = 8.872$, $P = <0.0001$**

ii. Tukey's multiple comparisons test

| | | Mouse | Human | | | | | | |
|---|---|---|---|---|---|---|---|---|---|
| | | Back | Back | Cheek | Chin | Eyebrow | Forehead | Front | Scalp |
| Human | Back | >0.9999 | | | | | | | |
| | Cheek | 0.0204 | 0.0817 | | | | | | |
| | Chin | <0.0001 | 0.0002 | 0.3581 | | | | | |
| | Eyebrow | <0.0001 | 0.0005 | 0.6298 | >0.9999 | | | | |
| | Forehead | 0.8593 | 0.965 | 0.611 | 0.0051 | 0.0149 | | | |
| | Front | >0.9999 | >0.9999 | 0.0727 | 0.0002 | 0.0006 | 0.9304 | | |
| | Scalp | 0.82 | 0.9493 | 0.6663 | 0.0066 | 0.0189 | >0.9999 | 0.9072 | |
| | Upper Lip | 0.0623 | 0.1684 | >0.9999 | 0.4626 | 0.7174 | 0.7617 | 0.1444 | 0.8041 |

iii. Number of samples of each age per area. Skin samples were taken from 11 individual human fetal specimens and 8 mouse specimens.

| | | Anatomical Site | | | | | | | |
|---|---|---|---|---|---|---|---|---|---|
| | | Back | Cheek | Chin | Eyebrow | Forehead | Front | Scalp | Upper Lip |
| Weeks Estimated Gestational Age | 10 | | | | 1 | | | | |
| | 11 | | | | | | | | |
| | 12 | | 3 | 3 | 2 | 1 | | 1 | 1 |
| | 13 | | 1 | 1 | 1 | 1 | | 1 | 1 |
| | 14 | 4 | 3 | 2 | 3 | 3 | 3 | 3 | 2 |
| | 15 | | | | | | | | |
| | 16 | 1 | 1 | 1 | 1 | 1 | 1 | 1 | 1 |
| | 17 | 1 | 1 | 1 | 1 | 1 | 1 | 1 | 1 |
| | 18 | | | | | | | | |
| | 19 | | | | | | | | |
| Mouse | E13.75 | 8 | | | | | | | |

**Figure 2c – Hair fibre width**

**i.** Ordinary one-way ANOVA between human anatomical sites

**F (7,28) = 3.203, P = 0.0127**

**ii.** Tukey's multiple comparisons test

| | Back | Cheek | Chin | Eyebrow | Forehead | Front | Scalp |
|---|---|---|---|---|---|---|---|
| Cheek | >0.9999 | | | | | | |
| Chin | >0.9999 | >0.9999 | | | | | |
| Eyebrow | 0.6302 | 0.131 | 0.0505 | | | | |
| Forehead | 0.8331 | 0.3501 | 0.1809 | 0.9941 | | | |
| Front | 0.9997 | 0.9988 | 0.9994 | 0.2635 | 0.4509 | | |
| Scalp | 0.9998 | 0.9971 | 0.9853 | 0.181 | 0.5317 | 0.9583 | |
| Upper Lip | 0.9365 | 0.6572 | 0.4733 | 0.9504 | 0.9998 | 0.6352 | 0.8828 |

**iii.** Number of samples of each age per area. Skin samples were taken from 11 individual specimens.

| | | Anatomical Site | | | | | | | |
|---|---|---|---|---|---|---|---|---|---|
| | | Back | Cheek | Chin | Eyebrow | Forehead | Front | Scalp | Upper Lip |
| Weeks Estimated Gestational Age | 10 | | | | | | | | |
| | 11 | | | | | | | | |
| | 12 | | | | | | | | |
| | 13 | | | | | | | | |
| | 14 | | | | 2 | | | | |
| | 15 | | | | | | | | |
| | 16 | | | 1 | 3 | 5 | | 3 | 2 |
| | 17 | | 2 | 2 | 2 | 2 | | 2 | 2 |
| | 18 | | | | | | | | |
| | 19 | 1 | 1 | 1 | 1 | 1 | 1 | 1 | 1 |

**Figure 2d – Dermal Condensate area**

i. Two-way ANOVA

**Between HF stage: F (6,301) = 124.4, P = <0.0001**

**Between anatomical sites: F (7,301) = 2.416, P = 0.0202**

ii. Tukey’s multiple comparisons tests between human anatomical sites

| | Back | Cheek | Chin | Eyebrow | Forehead | Front | Scalp |
|---|---|---|---|---|---|---|---|
| Back | | | | | | | |
| Cheek | 0.7768 | | | | | | |
| Chin | 0.9811 | 0.9978 | | | | | |
| Eyebrow | 0.286 | 0.9994 | 0.8953 | | | | |
| Forehead | 0.9752 | 0.9947 | >0.9999 | 0.8014 | | | |
| Front | 0.9821 | 0.2846 | 0.6227 | 0.0471 | 0.5342 | | |
| Scalp | 0.3016 | >0.9999 | 0.9438 | >0.9999 | 0.8602 | 0.0408 | |
| Upper Lip | >0.9999 | 0.8513 | 0.9885 | 0.4825 | 0.9864 | 0.9952 | 0.5226 |

iii. Number of samples of each age per area. Skin samples were taken from 15 individual specimens.

| | | Anatomical Site | | | | | | | |
|---|---|---|---|---|---|---|---|---|---|
| Weeks Estimated Gestational Age | | Back | Cheek | Chin | Eyebrow | Forehead | Front | Scalp | Upper Lip |
| | 10 | | | | | | | | |
| | 11 | | | | | | | | |
| | 12 | | | 1 | 2 | 3 | | 1 | 2 |
| | 13 | | | | | 1 | 1 | 1 | |
| | 14 | 1 | 1 | 1 | 1 | 2 | 1 | 2 | 1 |
| | 15 | | | | | 1 | | 1 | |
| | 16 | 3 | 1 | 1 | 1 | 4 | 2 | 4 | 1 |
| | 17 | | | | | | | | |
| | 18 | 1 | | | | | 1 | | |
| | 19 | 2 | 1 | 1 | 1 | 1 | 1 | 1 | 1 |

**Figure 2e (top) – Sebaceous gland area**

**i.** Ordinary one-way ANOVA between human anatomical sites

**F (7,34) = 0.8887, P = 0.5260**

**ii.** Tukey's multiple comparisons test

| | Back | Cheek | Chin | Eyebrow | Forehead | Front | Scalp |
|---|---|---|---|---|---|---|---|
| Back | | | | | | | |
| Cheek | 0.9871 | | | | | | |
| Chin | >0.9999 | 0.9989 | | | | | |
| Eyebrow | 0.5929 | 0.9753 | 0.6016 | | | | |
| Forehead | 0.9018 | >0.9999 | 0.9563 | 0.9786 | | | |
| Front | 0.9986 | >0.9999 | >0.9999 | 0.9596 | 0.9996 | | |
| Scalp | 0.9789 | >0.9999 | 0.9976 | 0.865 | 0.9997 | >0.9999 | |
| Upper Lip | 0.9021 | >0.9999 | 0.9583 | 0.9973 | >0.9999 | 0.9992 | 0.9994 |

**iii.** Number of samples of each age per area. Skin samples were taken from 14 individual specimens.

| | | Anatomical Site | | | | | | | |
|---|---|---|---|---|---|---|---|---|---|
| | | Back | Cheek | Chin | Eyebrow | Forehead | Front | Scalp | Upper Lip |
| Weeks Estimated Gestational Age | 10 | | | | | | | | |
| | 11 | | | | | | | | |
| | 12 | | | | | | | | |
| | 13 | | | | | | | | |
| | 14 | | | | 3 | 1 | | | |
| | 15 | | | | 1 | | | | |
| | 16 | | | 1 | 3 | 5 | | 4 | 2 |
| | 17 | | 2 | 2 | 2 | 2 | | 2 | 2 |
| | 18 | 1 | | | | | 1 | | |
| | 19 | 1 | 1 | 1 | 1 | 1 | 1 | 1 | 1 |

**Figure 2e (bottom) – Sebaceous gland density**

**i.** Ordinary one-way ANOVA between human anatomical sites

**F (7,34) = 1.712, P = 0.1393**

**ii.** Tukey's multiple comparisons test

| | Back | Cheek | Chin | Eyebrow | Forehead | Front | Scalp |
|---|---|---|---|---|---|---|---|
| Back | | | | | | | |
| Cheek | 0.9708 | | | | | | |
| Chin | 0.9889 | >0.9999 | | | | | |
| Eyebrow | >0.9999 | 0.7642 | 0.8335 | | | | |
| Forehead | 0.9981 | 0.4242 | 0.4684 | 0.9904 | | | |
| Front | >0.9999 | 0.9954 | 0.9992 | 0.9997 | 0.9779 | | |
| Scalp | 0.9787 | >0.9999 | >0.9999 | 0.6366 | 0.2326 | 0.998 | |
| Upper Lip | 0.98 | >0.9999 | >0.9999 | 0.7204 | 0.3256 | 0.998 | >0.9999 |

**iii.** Number of samples of each age per area. Skin samples were taken from 14 individual specimens.

| | | Anatomical Site | | | | | | | |
|---|---|---|---|---|---|---|---|---|---|
| | | Back | Cheek | Chin | Eyebrow | Forehead | Front | Scalp | Upper Lip |
| Weeks Estimated Gestational Age | 10 | | | | | | | | |
| | 11 | | | | | | | | |
| | 12 | | | | | | | | |
| | 13 | | | | | | | | |
| | 14 | | | | 3 | 1 | | | |
| | 15 | | | | 1 | | | | |
| | 16 | | | 1 | 3 | 5 | | 4 | 2 |
| | 17 | | 2 | 2 | 2 | 2 | | 2 | 2 |
| | 18 | 1 | | | | | 1 | | |
| | 19 | 1 | 1 | 1 | 1 | 1 | 1 | 1 | 1 |

**Figure 2f (bottom left) – Epidermal cell density**

i. Two-way ANOVA

**Between anatomical sites: F (7,68) = 5.777, P = <0.0001**

**Between ages: F (10,68) = 2.322, P = 0.0204**

ii. Tukey's multiple comparisons test between anatomical sites

| | Back | Cheek | Chin | Eyebrow | Forehead | Front | Scalp |
|---|---|---|---|---|---|---|---|
| Cheek | 0.0820 | | | | | | |
| Chin | 0.0009 | 0.9408 | | | | | |
| Eyebrow | 0.0331 | >0.9999 | 0.9862 | | | | |
| Forehead | 0.9808 | 0.4069 | 0.0170 | 0.2336 | | | |
| Front | 0.6756 | 0.6423 | 0.0434 | 0.4263 | 0.9992 | | |
| Scalp | 0.5731 | 0.6900 | 0.0518 | 0.4724 | 0.9974 | >0.9999 | |
| Upper Lip | 0.0017 | 0.9717 | >0.9999 | 0.9955 | 0.0284 | 0.0697 | 0.0825 |

iii. Tukey's multiple comparisons test between weeks estimated gestational age

| | | Human | | | | | | | | | |
|---|---|---|---|---|---|---|---|---|---|---|---|
| | | 9 | 10 | 11 | 12 | 13 | 14 | 15 | 16 | 17 | 18 |
| Human | 10 | >0.9999 | | | | | | | | | |
| | 11 | >0.9999 | >0.9999 | | | | | | | | |
| | 12 | >0.9999 | >0.9999 | >0.9999 | | | | | | | |
| | 13 | >0.9999 | >0.9999 | >0.9999 | >0.9999 | | | | | | |
| | 14 | 0.9605 | 0.6249 | 0.9819 | 0.3153 | 0.2212 | | | | | |
| | 15 | >0.9999 | >0.9999 | >0.9999 | >0.9999 | >0.9999 | 0.6508 | | | | |
| | 16 | 0.9996 | 0.9982 | >0.9999 | 0.9992 | 0.9686 | 0.7522 | 0.9932 | | | |
| | 17 | 0.7103 | 0.2944 | 0.7448 | 0.2587 | 0.1524 | 0.9734 | 0.3265 | 0.4950 | | |
| | 18 | 0.8203 | 0.5648 | 0.8581 | 0.5586 | 0.3891 | 0.9961 | 0.5208 | 0.7771 | >0.9999 | |
| | 19 | 0.9901 | 0.9194 | 0.9985 | 0.8713 | 0.7132 | >0.9999 | 0.8957 | 0.9950 | 0.9229 | 0.9820 |

**iv.** Number of samples of each age per area. Skin samples were taken from 31 individual specimens.

| Weeks Estimated Gestational Age | Anatomical Site | | | | | | | |
|---|---|---|---|---|---|---|---|---|
| | Back | Cheek | Chin | Eyebrow | Forehead | Front | Scalp | Upper Lip |
| 9 | 1 | | | | | | | |
| 10 | 3 | | | | | | 2 | |
| 11 | 1 | | | | | 1 | | |
| 12 | 5 | 1 | 1 | 1 | 2 | 2 | 3 | 1 |
| 13 | 4 | | | | 1 | 2 | 2 | |
| 14 | 5 | 1 | 1 | 1 | 2 | 3 | 2 | 1 |
| 15 | 1 | | | | 1 | | 1 | |
| 16 | 5 | 1 | 1 | 1 | 4 | 3 | 4 | 1 |
| 17 | 3 | | | | | | | |
| 18 | 1 | | | | | 1 | | |
| 19 | 2 | 1 | 1 | 1 | 1 | 1 | 1 | 1 |

**Figure 2f (bottom right) – Epidermal cell proliferation**

**i.** Two-way ANOVA

**Between anatomical sites: F (7,67) = 0.2011, P = 0.9842**

**Between ages: F (10,67) = 5.623, P = <0.0001**

**ii.** Tukey's multiple comparisons test between anatomical sites

| | Back | Cheek | Chin | Eyebrow | Forehead | Front | Scalp |
|---|---|---|---|---|---|---|---|
| Cheek | >0.9999 | | | | | | |
| Chin | >0.9999 | >0.9999 | | | | | |
| Eyebrow | >0.9999 | >0.9999 | >0.9999 | | | | |
| Forehead | 0.9979 | 0.9972 | >0.9999 | 0.9992 | | | |
| Front | >0.9999 | >0.9999 | >0.9999 | >0.9999 | 0.9887 | | |
| Scalp | >0.9999 | >0.9999 | >0.9999 | >0.9999 | 0.9998 | 0.9997 | |
| Upper Lip | 0.9989 | >0.9999 | 0.9990 | >0.9999 | 0.9794 | >0.9999 | 0.9975 |

**iii.** Tukey's multiple comparisons test between weeks estimated gestational age

| | | Human | | | | | | | | | |
|---|---|---|---|---|---|---|---|---|---|---|---|
| | | 9 | 10 | 11 | 12 | 13 | 14 | 15 | 16 | 17 | 18 |
| Human | 10 | 0.1248 | | | | | | | | | |
| | 11 | 0.9201 | 0.8405 | | | | | | | | |
| | 12 | 0.0718 | >0.9999 | 0.6696 | | | | | | | |
| | 13 | 0.1928 | >0.9999 | 0.9424 | 0.9971 | | | | | | |
| | 14 | 0.8368 | 0.1615 | >0.9999 | 0.0018 | 0.1840 | | | | | |
| | 15 | 0.2212 | >0.9999 | 0.9393 | >0.9999 | >0.9999 | 0.5231 | | | | |
| | 16 | 0.8376 | 0.1327 | >0.9999 | 0.0008 | 0.1404 | >0.9999 | 0.4789 | | | |
| | 17 | 0.9870 | 0.2466 | >0.9999 | 0.0907 | 0.3823 | 0.9998 | 0.5060 | 0.9998 | | |
| | 18 | >0.9999 | 0.1380 | 0.9920 | 0.0470 | 0.1981 | 0.9653 | 0.2920 | 0.9658 | >0.9999 | |
| | 19 | 0.9699 | 0.0599 | >0.9999 | 0.0008 | 0.0683 | 0.9972 | 0.2596 | 0.9969 | >0.9999 | 0.9996 |

**iv.** Number of samples of each age per area. Skin samples were taken from 31 individual specimens.

| | | Anatomical Site | | | | | | | |
|---|---|---|---|---|---|---|---|---|---|
| | | Back | Cheek | Chin | Eyebrow | Forehead | Front | Scalp | Upper Lip |
| Weeks Estimated Gestational Age | 9 | 1 | | | | | | | |
| | 10 | 3 | | | | | | 2 | |
| | 11 | 1 | | | | | 1 | | |
| | 12 | 4 | 1 | 1 | 1 | 2 | 2 | 3 | 1 |
| | 13 | 4 | | | | 1 | 2 | 2 | |
| | 14 | 5 | 1 | 1 | 1 | 2 | 3 | 2 | 1 |
| | 15 | 1 | | | | 1 | | 1 | |
| | 16 | 5 | 1 | 1 | 1 | 4 | 3 | 4 | 1 |
| | 17 | 3 | | | | | | | |
| | 18 | 1 | | | | | 1 | | |
| | 19 | 2 | 1 | 1 | 1 | 1 | 1 | 1 | 1 |

**Figure 2f (top left) – Dermal cell density**

**i.** Two-way ANOVA

**Between anatomical sites: F (7,68) = 6.275, P = <0.0001**

**Between ages: F (10,68) = 1.954, P = 0.0525**

**ii.** Tukey’s multiple comparisons test between anatomical sites

| | Back | Cheek | Chin | Eyebrow | Forehead | Front | Scalp |
|---|---|---|---|---|---|---|---|
| Cheek | 0.7248 | | | | | | |
| Chin | 0.0003 | 0.2511 | | | | | |
| Eyebrow | 0.9942 | 0.9974 | 0.0551 | | | | |
| Forehead | >0.9999 | 0.8012 | 0.0010 | 0.9968 | | | |
| Front | 0.9993 | 0.9259 | 0.0024 | >0.9999 | 0.9999 | | |
| Scalp | >0.9999 | 0.8509 | 0.0010 | 0.9991 | >0.9999 | >0.9999 | |
| Upper Lip | 0.0003 | 0.2629 | >0.9999 | 0.0587 | 0.0011 | 0.0027 | 0.0012 |

**iii.** Tukey’s multiple comparisons test between weeks estimated gestational age

| | | Human | | | | | | | | | |
|---|---|---|---|---|---|---|---|---|---|---|---|
| | | 9 | 10 | 11 | 12 | 13 | 14 | 15 | 16 | 17 | 18 |
| Human | 10 | 0.2605 | | | | | | | | | |
| | 11 | 0.4836 | >0.9999 | | | | | | | | |
| | 12 | 0.0936 | 0.9999 | >0.9999 | | | | | | | |
| | 13 | 0.0402 | 0.9643 | 0.9910 | 0.9966 | | | | | | |
| | 14 | 0.2674 | >0.9999 | >0.9999 | 0.9390 | 0.5526 | | | | | |
| | 15 | 0.0587 | 0.9710 | 0.9878 | 0.9969 | >0.9999 | 0.8161 | | | | |
| | 16 | 0.0428 | 0.9759 | 0.9954 | 0.9979 | >0.9999 | 0.3891 | >0.9999 | | | |
| | 17 | 0.3912 | >0.9999 | >0.9999 | 0.9997 | 0.9690 | >0.9999 | 0.9693 | 0.9809 | | |
| | 18 | 0.2416 | >0.9999 | >0.9999 | >0.9999 | >0.9999 | 0.9996 | >0.9999 | >0.9999 | >0.9999 | |
| | 19 | 0.1040 | 0.9999 | >0.9999 | >0.9999 | 0.9996 | 0.9659 | 0.9991 | >0.9999 | 0.9996 | >0.9999 |

**iv.** Number of samples of each age per area. Skin samples were taken from 31 individual specimens.

| | | Anatomical Site | | | | | | | |
|---|---|---|---|---|---|---|---|---|---|
| | | Back | Cheek | Chin | Eyebrow | Forehead | Front | Scalp | Upper Lip |
| Weeks Estimated Gestational Age | 9 | 1 | | | | | | | |
| | 10 | 3 | | | | | | 2 | |
| | 11 | 1 | | | | | 1 | | |
| | 12 | 5 | 1 | 1 | 1 | 2 | 2 | 3 | 1 |
| | 13 | 4 | | | | 1 | 2 | 2 | |
| | 14 | 5 | 1 | 1 | 1 | 2 | 3 | 2 | 1 |
| | 15 | 1 | | | | 1 | | 1 | |
| | 16 | 5 | 1 | 1 | 1 | 4 | 3 | 4 | 1 |
| | 17 | 3 | | | | | | | |
| | 18 | 1 | | | | | 1 | | |
| | 19 | 2 | 1 | 1 | 1 | 1 | 1 | 1 | 1 |

**Figure 2f (top right) – Dermal cell proliferation**

**i.** Two-way ANOVA

**Between anatomical sites: F (7,67) = 2.170, P = 0.0479**

**Between ages: F (10,67) = 4.749, P = <0.0001**

**ii.** Tukey's multiple comparisons test between anatomical sites

| | Back | Cheek | Chin | Eyebrow | Forehead | Front | Scalp |
|---|---|---|---|---|---|---|---|
| Cheek | 0.6403 | | | | | | |
| Chin | 0.3711 | >0.9999 | | | | | |
| Eyebrow | 0.0654 | 0.9768 | 0.9980 | | | | |
| Forehead | 0.9891 | 0.9517 | 0.8043 | 0.3217 | | | |
| Front | 0.6188 | 0.9988 | 0.9749 | 0.6412 | 0.9956 | | |
| Scalp | 0.4996 | 0.9992 | 0.9785 | 0.6504 | 0.9905 | >0.9999 | |
| Upper Lip | 0.6629 | >0.9999 | >0.9999 | 0.9730 | 0.9586 | 0.9992 | 0.9995 |

**iii.** Tukey's multiple comparisons test between weeks estimated gestational age

| Human | | Human | | | | | | | | | |
|---|---|---|---|---|---|---|---|---|---|---|---|
| | | 9 | 10 | 11 | 12 | 13 | 14 | 15 | 16 | 17 | 18 |
| Human | 10 | 0.2282 | | | | | | | | | |
| | 11 | 0.9998 | 0.3247 | | | | | | | | |
| | 12 | 0.8831 | 0.3529 | 0.9905 | | | | | | | |
| | 13 | 0.8882 | 0.4563 | 0.9915 | >0.9999 | | | | | | |
| | 14 | >0.9999 | 0.0002 | 0.9998 | 0.0131 | 0.0615 | | | | | |
| | 15 | 0.9454 | 0.7761 | 0.9985 | >0.9999 | >0.9999 | 0.5877 | | | | |
| | 16 | 0.9988 | 0.0061 | >0.9999 | 0.4500 | 0.6828 | 0.8165 | 0.9792 | | | |
| | 17 | >0.9999 | 0.0089 | 0.9989 | 0.3542 | 0.3971 | >0.9999 | 0.7245 | 0.9700 | | |
| | 18 | >0.9999 | 0.0051 | 0.9510 | 0.1541 | 0.1763 | 0.9895 | 0.4268 | 0.7287 | 0.9999 | |
| | 19 | >0.9999 | 0.0031 | >0.9999 | 0.1972 | 0.3744 | >0.9999 | 0.8444 | 0.9968 | >0.9999 | 0.9674 |

**iv.** Number of samples of each age per area. Skin samples were taken from 31 individual specimens.

| | | Anatomical Site | | | | | | | |
|---|---|---|---|---|---|---|---|---|---|
| | | Back | Cheek | Chin | Eyebrow | Forehead | Front | Scalp | Upper Lip |
| Weeks Estimated Gestational Age | 9 | 1 | | | | | | | |
| | 10 | 3 | | | | | | 2 | |
| | 11 | 1 | | | | | 1 | | |
| | 12 | 4 | 1 | 1 | 1 | 2 | 2 | 3 | 1 |
| | 13 | 4 | | | | 1 | 2 | 2 | |
| | 14 | 5 | 1 | 1 | 1 | 2 | 3 | 2 | 1 |
| | 15 | 1 | | | | 1 | | 1 | |
| | 16 | 5 | 1 | 1 | 1 | 4 | 3 | 4 | 1 |
| | 17 | 3 | | | | | | | |
| | 18 | 1 | | | | | 1 | | |
| | 19 | 2 | 1 | 1 | 1 | 1 | 1 | 1 | 1 |

**Figure 2g – Distribution of proliferation in hair follicles**

**i.** Ordinary one-way ANOVA between stage 2/3 placodes and the top and bottom of stage 4+ placodes

**F (2,48) = 7.620, P = 0.0013**

**ii.** Tukey's multiple comparisons test

| | Stage 2/3 Entire Placode | Stage 4+ Top Half |
|---|---|---|
| Stage 4+ Top Half | 0.9977 | |
| Stage 4+ Bottom Half | 0.0050 | 0.0038 |

**iii.** Number of samples of each age. 14 individual specimens were used.

| | | # of samples |
|---|---|---|
| Weeks Estimated Gestational Age | | |
| | 11 | |
| | 12 | 1 |
| | 13 | 2 |
| | 14 | 4 |
| | 15 | 1 |
| | 16 | 4 |
| | 17 | 1 |
| | 18 | |
| | 19 | 1 |
